# Spatial gene expression maps in vertebrate limbs display conserved and regenerative species-specific features within connective tissue

**DOI:** 10.64898/2025.12.15.694399

**Authors:** Conor L. McMann, Chanyoung Park, Jennifer K. Cloutier, Peter W. Reddien

## Abstract

Regeneration is widespread but sparsely distributed throughout the animal kingdom. Identifying factors that differentiate regenerative and non-regenerative organisms could enable approaches for improving regenerative outcomes in non-regenerative species. Constitutive adult positional information can be required for regeneration, but has been poorly characterized across animals. Here, we generated positional gene expression atlases for the limbs of one regenerative (axolotl) and one non-regenerative (mouse) vertebrate. Regional gene expression signatures in both species are highly overlapping and mirror multiple developmental positional information patterns, particularly along the primary limb axis. These expression signatures are largely harbored in connective tissue, including diverse fibroblast types, in both organisms. We also identified species-specific regional expression patterns, including for *Proxima*, a novel gene encoding a secreted factor with strong positional expression in axolotl. Positional gene expression similar to developmental patterns also exists between forelimbs and hindlimbs and along anterior-posterior and dorsal-ventral limb axes, although some of these regional expression signatures are stronger in axolotl than in mouse. This work establishes regional atlases of adult vertebrate limbs and suggests that the connective tissue of regenerative and non-regenerative vertebrate limbs share a conserved signature of positional memory, with some signatures more apparent in the regenerative species.

## Introduction

Most major animal phyla contain species capable of regenerating lost body parts^1–3^. Analyses suggest that some regenerative mechanisms have shared evolutionary origins^4–8^. However, the properties that differentiate organisms capable and incapable of regeneration remain poorly understood. One approach to understanding the evolutionary basis for regeneration loss and/or gain is to identify mechanisms that enable regeneration in model systems and to determine the distribution of such mechanisms across species.

Adult positional information – factors that influence the regional identity and organization of cells^9,10^ – is a central component of regeneration in planarians^11^, acoels^4,5^, hydra^12^, and axolotls^13–16^. Several indications suggest that adult positional information or memory of developmental position might be harbored in connective tissue (CT) in vertebrates.

For example, dermal and intestinal fibroblasts can influence epithelial morphology in a regionally dependent manner^17–27^. Human fibroblasts display different gene expression signatures based on organ and body position, sometimes mirroring developmental origin, as reflected by expression of particular *Hox* genes^28–31^. In mouse digit tip regeneration, fibroblasts contribute substantially to the blastema^32,33^ and incorporate into the new digit tip based on positional origin^34^. In the zebrafish pectoral fin, fibroblasts and osteoblasts show regional expression of genes involved in developmental patterning^35^. In axolotls, CT incorporates into regenerating limb tissue in a manner that reflects its positional origin, a behavior not found for muscle or Schwann cells^36,37^.

Chromatin profiling of axolotl CT has revealed position-specific signatures within the limb that reflect developmental patterning programs^38^. Posterior expression of *Hand2* in uninjured axolotl limb CT is required for proper patterning in the regenerating limb^16^.

In planarians, position control genes (PCGs) display constitutive, regional expression in adults and regulate the pattern of tissue identity^11,39^. PCGs include members of canonical developmental signaling pathways and PCG expression patterns are primarily harbored in muscle^40^. Resetting of positional information after injury is required for planarian regeneration after amputation^11,41–44^. A similar adult positional information system is present in acoels^5^ and, like planarians, it is primarily harbored in muscle^4^.

Current models indicate that acoels are members of a sister clade to all other bilaterians, having diverged from the rest of the Bilateria over 550 million years ago^5,45–49^. The similarities between these two clades raise the possibility that a similar system of adult positional information was present at the base of Bilateria^4^. In planarians, muscle serves as the primary CT^50^ in addition to being the main source of PCG expression. This is consistent with CT-like cell types producing extracellular matrix (ECM) components and harboring adult positional information potentially being a unifying attribute of regeneration across many clades^50^.

In this work, we developed single-cell spatial gene expression atlases of the adult limbs of two well-studied model vertebrate systems: one capable (axolotl) and the other incapable (mouse) of limb regeneration. Comparison of spatial features in two tetrapod systems, separated by hundreds of millions of years of evolution and with different regenerative capacity, might enable the identification of vertebrate positional memory molecules and patterns, as well as yield insight into the evolution of positional memory in tetrapods. We developed spatial cell-type transcriptome atlases of these vertebrate limbs and examined the adult, constitutive expression of these developmental factors, together with any other regionally expressed genes across limb axes. Regenerative and non-regenerative vertebrate limbs shared key characteristics, including regional expression of a broad set of developmental TFs that recapitulated developmental patterns, a general lack of constitutive regional expression of developmental signaling factors, and a prominent role for CT in the regional expression of developmental patterning factors. Certain spatial expression features were more robust in axolotl limbs than in mouse limbs and some regional expression patterns were species-specific.

These data are consistent with a model in which CT similarly expresses genes associated with developmental positional information in adult regenerative and non-regenerative vertebrates.

## Results

### Generation of spatial atlases of gene expression across adult vertebrate limbs

To assess the scope and diversity of region-restricted gene expression within adult vertebrate limbs, we generated spatial limb gene expression datasets for axolotl and mouse limbs. We collected diverse tissue from every layer of forelimbs and hindlimbs, and also segmented hindlimbs along proximal-distal (PD), dorsal-ventral (DV), and anterior-posterior (AP) axes (Fig. 1A, see Methods). We performed single-cell RNA sequencing (scRNA-seq) on each limb segment to generate spatial limb cell-type transcriptome atlases (Fig. 1B, Supplementary Fig. 1A-C, Supplementary Data 1). The resultant vertebrate limb atlases are composed of approximately 86,000 mouse and 197,000 axolotl cells. We also generated bulk RNA-sequencing (RNA-seq) datasets of skin – samples which included epidermis, dermis, and hypodermis while excluding skeletal muscle and deep connective tissue – taken from proximal, medial, and distal limb segments for each species. We chose to limit this bulk RNA-sequencing dataset to skin because it includes many cell types, including dermal CT, and because limb cell-type composition is varied across the PD axis; using full limb regions would strongly bias differentially expressed genes toward markers of non-uniformly distributed cell types^51^. *Tig1*, which is associated with proximal limb identity in axolotl^15^, displayed enriched proximal/medial expression, demonstrating that the bulk RNA-seq dataset can be used to detect meaningful PD expression differences (Supplementary Fig. 1D).

**Figure 1.**
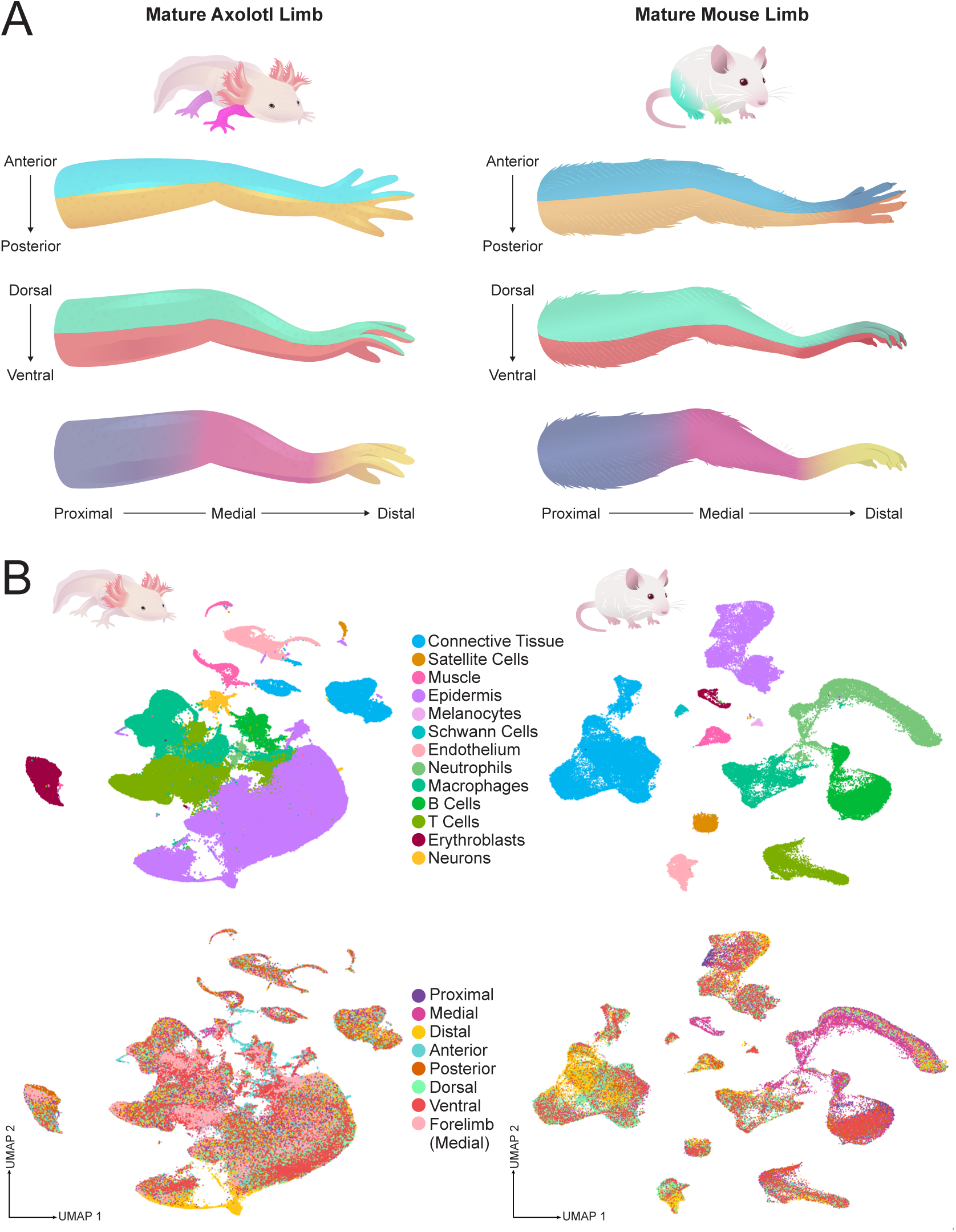
Generation of spatial atlases of the limbs of mouse and axolotl. (A) Schematic illustrating the axes of adult limbs of axolotl (left) and mouse (right). For tissue collection, medial sections of forelimbs were taken for comparison to hindlimbs. Hindlimbs were segmented along the PD axis into proximal, medial, and distal segments by bisecting the knee and ankle joints. Hindlimbs were also segmented along the DV and AP axes by surgically bisecting the tissue at the midline of each axis and separately harvesting tissue from each side. For axolotl DV and AP segmentation, only the medial portion was used to reduce PD variability within these tissues. (B) UMAP representation of scRNA-seq atlases of axolotl (left) and mouse (right) limbs, colored by cell type (top) and position of origin (bottom).

### A CT transcriptome atlas of the adult vertebrate limb

Given the functional role of CT in positional memory, we analyzed this tissue in detail. In both organisms, subclustering revealed a diverse set of CT subtypes, including multiple fibroblast types, osteoblasts, tenocytes, and chondrocytes (Fig. 2A-B, Supplementary Fig. 2A-B, Supplementary Data 2). Fibroblasts comprised the majority of CT in both datasets (Fig. 2A-B). Whereas it is known that fibroblasts are heterogeneous, this heterogeneity is just beginning to be dissected^28,52–58^. Subclustering revealed seven distinct limb fibroblast classes in axolotl and eight in mouse. Each fibroblast class displayed unique expression profiles that included genes encoding distinct core ECM components, secreted matrix-associated factors, and TFs (Fig 2C). Mouse CT additionally included two characterized fibroblast subtypes – myofibroblasts and dermal papillary fibroblasts^59^ that also had distinct ECM and TF profiles (Supplementary Fig. 2B). Given that ECM production is a core function of fibroblasts^52,60^, these specific ECM expression profiles suggest distinct functional roles for these fibroblast populations within limbs.

**Figure 2.**
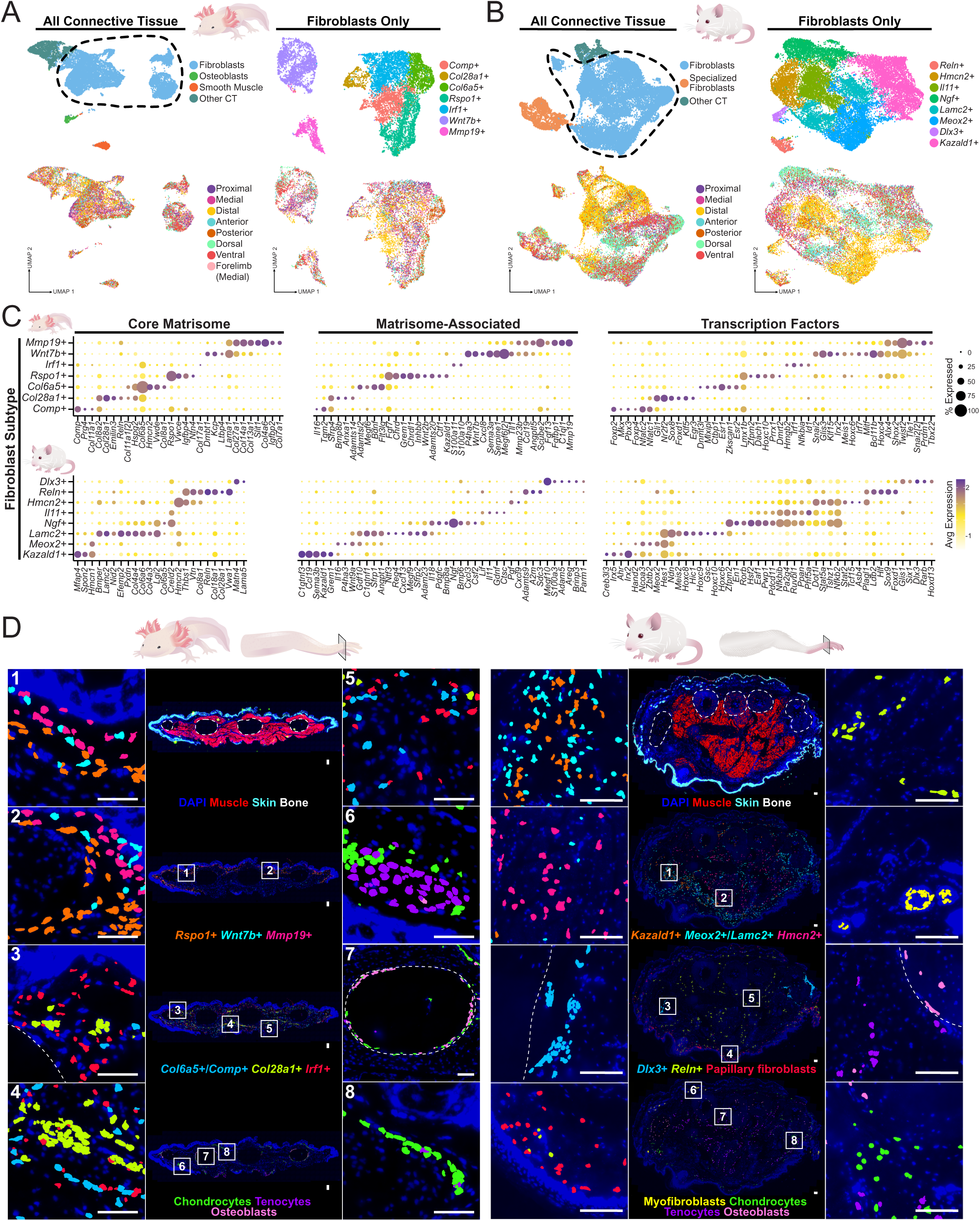
Spatial and expression heterogeneity of vertebrate limb fibroblasts. (A-B) UMAP of all CT cells (left) and subclustered fibroblasts (right) from axolotl (A) and mouse (B) limbs, colored by subtype (top) and position of origin (bottom). (C) Expression of genes encoding core matrisome components (left), matrisome-associated factors (center), and TFs (right) that differentiate fibroblast subtypes in the axolotl (top) and mouse (bottom) limb. (D) Spatial localization of fibroblast and CT subtypes within distal cross-section of the axolotl and mouse distal limb with MERFISH. Zoomed insets (left and right) are labelled on image of full cross-section (center). Dotted white lines outline bone. Scale bars, 100 µm.

We used MERFISH^61,62^, a spatial transcriptomics method used to visualize the expression of hundreds of genes simultaneously, to localize fibroblast types within full sections of mouse and axolotl distal limbs (Fig. 2D). In axolotl, *Mmp19*+, *Wnt7b*+, and *Rspo1*+ fibroblasts formed loose dermal layers, with *Mmp19*+ cells closest to the epidermis and *Rspo1*+ cells farthest, often projecting deeper into the dermis. *Comp*+, *Col6a5*+, and *Irf1*+ fibroblasts, on the other hand, were more internal and were integrated within muscle, with *Irf1*+ fibroblasts also found in the dermis (Fig. 2D). *Col28a1*+ fibroblasts were found in tight bundles scattered through limb tissue layers, as was the case for *Reln*+ fibroblasts in the mouse limb (Fig. 2D). Mouse *Kazald1*+, *Meox2*+, and *Lamc2*+ fibroblasts were intermingled throughout the dermis, whereas *Hmcn2*+ fibroblasts were primarily localized within the muscle and *Dlx3*+ fibroblasts around bone (Fig. 2D). In both organisms, chondrocytes were present in and around bone and tenocytes were more often muscle-associated, although cartilage was also sometimes muscle-associated in axolotl limbs (Fig. 2D). Mouse dermal papillary fibroblasts were observed in their expected location in the upper dermis^63,64^ and myofibroblasts were also found around hair follicles (Fig. 2D). The localization and myofibroblast-like gene expression of these cells is consistent with a dermal sheath cell identity^65^. Overall, these data suggest that a large degree of expression and functional fibroblast diversity exists in vertebrate limbs.

Axolotl *Col28a1*+ fibroblasts and mouse *Reln*+ fibroblasts expressed similar ECM and TF-encoding genes, including those encoding structural ECM components (e.g., collagens) and several genes associated with neural and/or glia function (e.g., *Reln*, *Col28a1*, and *Sox9*), suggesting a possible affiliation with neurites (Supplementary Fig. 2C). *Sox9* and *Col28a1* expression have previously been associated with endoneurial mesenchymal cells, which intermingle with axons and glial cells in nerve bundles^53^, supporting this possibility. Axolotl *Rspo1*+ fibroblasts and mouse *Kazald1*+ fibroblasts also expressed multiple orthologous genes, particularly those encoding core ECM components (Supplementary Fig. 2C). Both axolotl *Comp*+ fibroblasts and mouse *Dlx3*+ fibroblasts displayed enriched expression of markers previously associated with synovial fibroblasts^66–68^, which are found in the mesenchymal lining between joints, suggesting a similar functional identity in their respective limb tissue contexts (Supplementary Fig. 2D). Beyond these cases of similarity, there was unexpectedly little overlap in ECM and TF markers between fibroblast populations of mouse and axolotl.

Whereas most broad classes of cell types within the limb (e.g., immune cells, skin, endothelium) were easily identifiable using markers generally shared between organisms (Supplementary Fig. 1A), this was not possible with fibroblasts subtypes. Broad phyletic sampling will be an important direction to undertake for understanding of the evolution of these limb fibroblast types.

### Major hindlimb- and forelimb-distinguishing developmental factors reprise expression constitutively in adult vertebrate limb CT

In vertebrate limb development, hindlimb outgrowth and identity are largely controlled by the TF genes *Tbx4* and *Pitx1*, whereas formation of the forelimb is primarily guided by the TF gene *Tbx5*^69–76^. In axolotl, the top overall differentially expressed genes genome-wide between adult forelimb and hindlimb CT genome-wide were *Tbx5* (in the forelimb) and *Tbx4* (in the hindlimb) (Fig. 3A, Supplementary Data 3). *Pitx1* also showed prominent hindlimb-enriched expression (Fig. 3A, Supplementary Data 3). The expression of *Tbx5, Tbx4*, and *Pitx1* was predominantly in CT, indicating CT-specificity in the regional retention of hindlimb-versus-forelimb identity factors (Fig. 3B-D).

**Figure 3.**
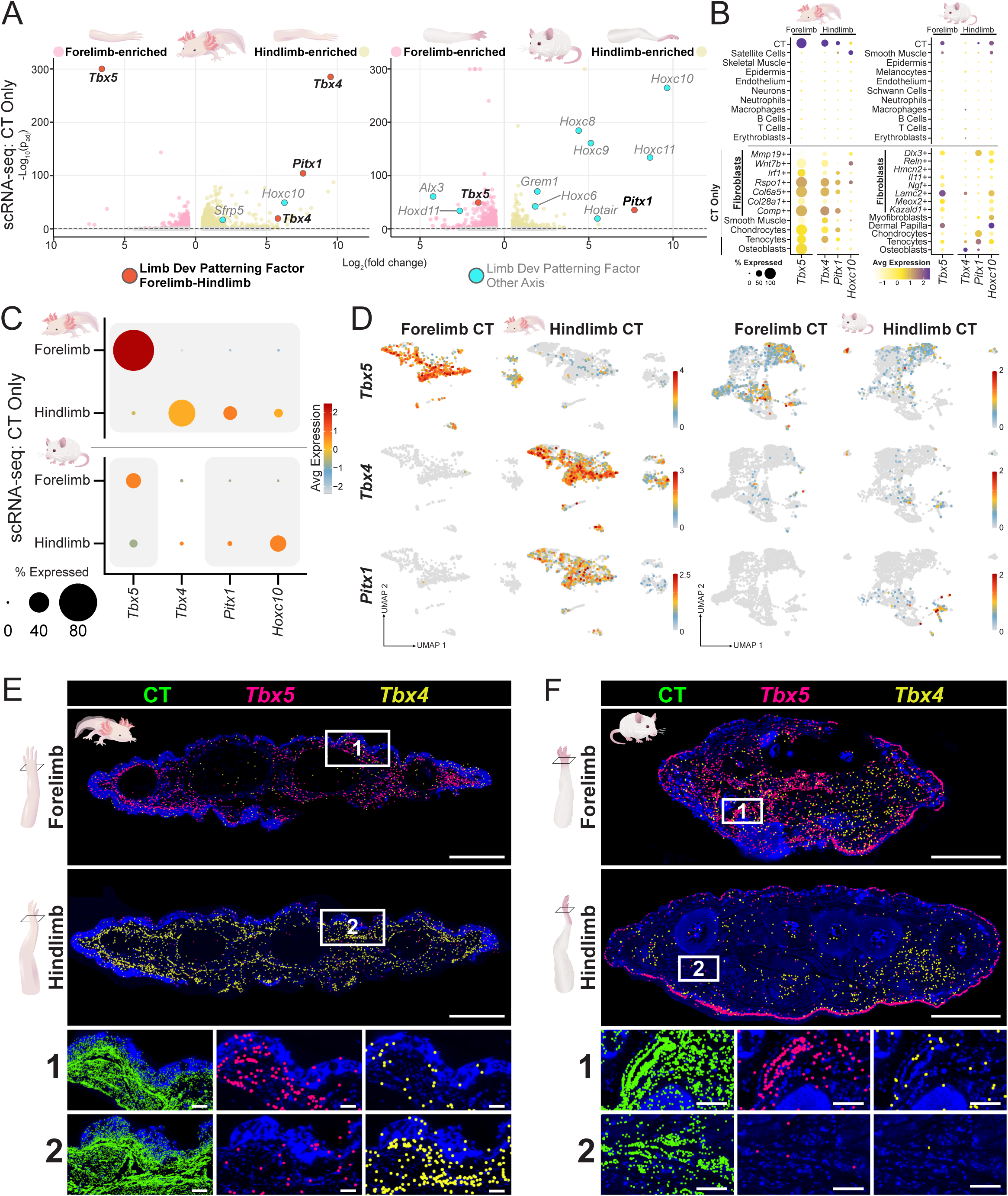
Expression of developmental forelimb and hindlimb factors in adult vertebrate limbs. (A) Volcano plot depicting top differentially expressed genes between forelimb and hindlimb CT in axolotl (left) and mouse (right). Dotted line represents cutoff at p_adj_<0.05. The lower scoring *Tbx4* locus from the axolotl genome annotation was excluded from further analysis (see Supplementary Data 16). (B) Expression of *Tbx5*, *Tbx4*, and *Pitx1* within different cell types present in scRNA-seq atlases (top) and CT subtypes (bottom) for axolotl (left) and mouse (right). Only medial tissue from each limb is represented for better comparison. (C) Expression of *Tbx5*, *Tbx4*, and *Pitx1* within hindlimb and forelimb CT in axolotl (top) and mouse (bottom). Grey backdrop indicates a statistically significant difference in expression between hindlimb and forelimb CT (p_adj_<0.05). (D) UMAP visualization of *Tbx5*, *Tbx4*, and *Pitx1* expression within hindlimb and forelimb CT in axolotl (left) and mouse (right). (E) MERFISH visualization of *Tbx4* and *Tbx5* expression in axolotl (top) and mouse (bottom) forelimb and hindlimb CT. CT was visualized via expression of *Pdgfra*, *Dpt*, and *Col6a1* in both organisms. This pool additionally included *Col4a1* in mouse. Scale bars, 1mm for full cross-section, 100 µm for insets.

Expression of these genes was also widely expressed across axolotl CT subsets, indicating broad retention of forelimb-versus-hindlimb positional memory across the diverse cell types of the CT (Fig. 3B, Supplementary Fig. 3A).

In mouse, *Tbx5* was also constitutively expressed and was significantly more expressed in forelimb over hindlimb CT (Fig. 3A). Mouse *Pitx1* was significantly more expressed in hindlimb than forelimb CT, but it was expressed at lower levels overall compared to the expression of *Pitx1* in axolotl hindlimb CT (Fig. 3A-D). By contrast, *Tbx4 was* only lowly expressed in mouse CT and did not show a significant expression difference between forelimb and hindlimb (Fig 3A,C, Supplementary Data 3). Mouse *Tbx5* expression was somewhat widespread within CT, whereas *Pitx1* expression was driven by a small population of *Dlx3+* fibroblasts and tenocytes (Fig. 3B,D, Supplementary Fig. 3A, Supplementary Fig. 4, Supplementary Data 4). With MERFISH in both mouse and axolotl, *Tbx5* expression was widespread across fibroblast types and expression was strikingly higher throughout forelimb CT than in hindlimb CT (Fig. 3E). MERFISH also showed notably higher axolotl *Tbx4* expression in hindlimb CT than in forelimb CT, but this expression difference was not observed for mouse *Tbx4* (Fig. 3E). Visium HD, which enabled the spatial localization of transcripts within the full endogenous mouse limb across different positions of forelimb and hindlimb, also showed forelimb *Tbx5* expression and comparable forelimb and hindlimb *Tbx4* expression in mouse (Supplementary Fig. 3B). Overall, both organisms showed constitutive *Tbx5* and *Pitx1* expression in line with developmental expression patterns, although this expression was more widespread in axolotl. Axolotl CT also displayed hindlimb *Tbx4* expression (Fig. 3B-E).

Some *HoxC* cluster genes also showed hindlimb-enriched expression, although more prominently in mouse than axolotl (Fig. 3A). These genes were expressed in CT, but not exclusively (Fig. 3B, Supplementary Fig. 3C). *Hoxc10* displayed hindlimb-enriched expression in both mouse and axolotl by both scRNA-seq and MERFISH (Fig. 3A-C, Supplementary Fig. 3D). *Hoxc10* is expressed in developing axolotl hindlimbs^77^ and heterozygous *Hoxc10 - Hoxc9* deletion causes hindlimb defects in humans^78^. In general, *HoxC* cluster genes (specifically *c9*-*c13*) show more clear expression in developing hindlimbs than forelimbs^79^, mirroring the adult expression of several *HoxC* genes found here, particularly in mouse.

Computational analysis of Visium HD data yielded further insight into differentially expressed genes between forelimb and hindlimb mouse paw tissue. Although this method did not yield robust results for CT, likely because of lower transcript density, it was able to robustly characterize differences between forepaw and hindpaw epidermis (Supplementary Data 5). *Hoxa7* and *Hoxa9* were expressed specifically in hindlimb epidermis; this was corroborated by MERFISH (Supplementary Fig. 3E-F, Supplementary Data 5).

Together, these data suggest that CT in the limbs of both axolotl and mouse constitutively express major TFs associated with forelimb-versus-hindlimb specification throughout CT cell types, with some species-specific differences.

### *Hox* genes in mouse and axolotl limbs recapitulate PD developmental expression

*Hox* genes are famous for their role in patterning the primary body axis^80–82^. In the evolution of tetrapod limbs, this pattern-regulating system was re-deployed for patterning the major (PD) axis of limb outgrowth: for the hindlimb, group 10 *Hox* genes promote proximal limb identity^83^, group 11 *Hox* genes promote medial limb identity^83–86^, and group 12 and 13 *Hox* genes promote distal limb identity^87–90^. The forelimb additionally utilizes group 9 *Hox* genes for promotion of proximal arm identity^89,91^. *HoxA* and *HoxD* clusters are particularly important for PD limb developmental patterning^88,89,92,93^. Some limb developmental *Hox* genes have positionally restricted expression in the axolotl limb blastema during regeneration^94^. The human dermis is known to express certain *Hox* genes in a positionally restricted manner^29–31^ and some evidence suggests this may be true for axolotl as well^38^. However, the cell-type specificity of *Hox* gene expression in the adult limb and how expression compares between regenerative and non-regenerative adult limbs is poorly understood.

In both axolotl and mouse bulk skin RNA-seq data, some group 10 *Hox* genes, predominantly members of the *HoxA* and *HoxD* clusters, showed constitutive proximal/medial limb expression, some group 11 genes showed medial/distal expression, and group 13 genes displayed distal expression, largely mirroring *Hox* expression during limb development (Fig. 4A, Supplementary Data 6). These group 10- 13 *Hox* genes showed strong selectivity in their expression to CT, with some minor expression in satellite cells and smooth muscle (Fig. 4B). Expression of these *Hox* genes was seen in many different CT subtypes in both organisms, including diverse fibroblast types spanning from dermis to deeper tissues, as well as in tenocytes, chondroblasts, and osteoblasts (Fig. 4B). For example, *Hoxa13* was detectable by MERFISH in most CT subsets in distal mouse and axolotl limb tissue (Supplementary Fig. 5A).

**Figure 4.**
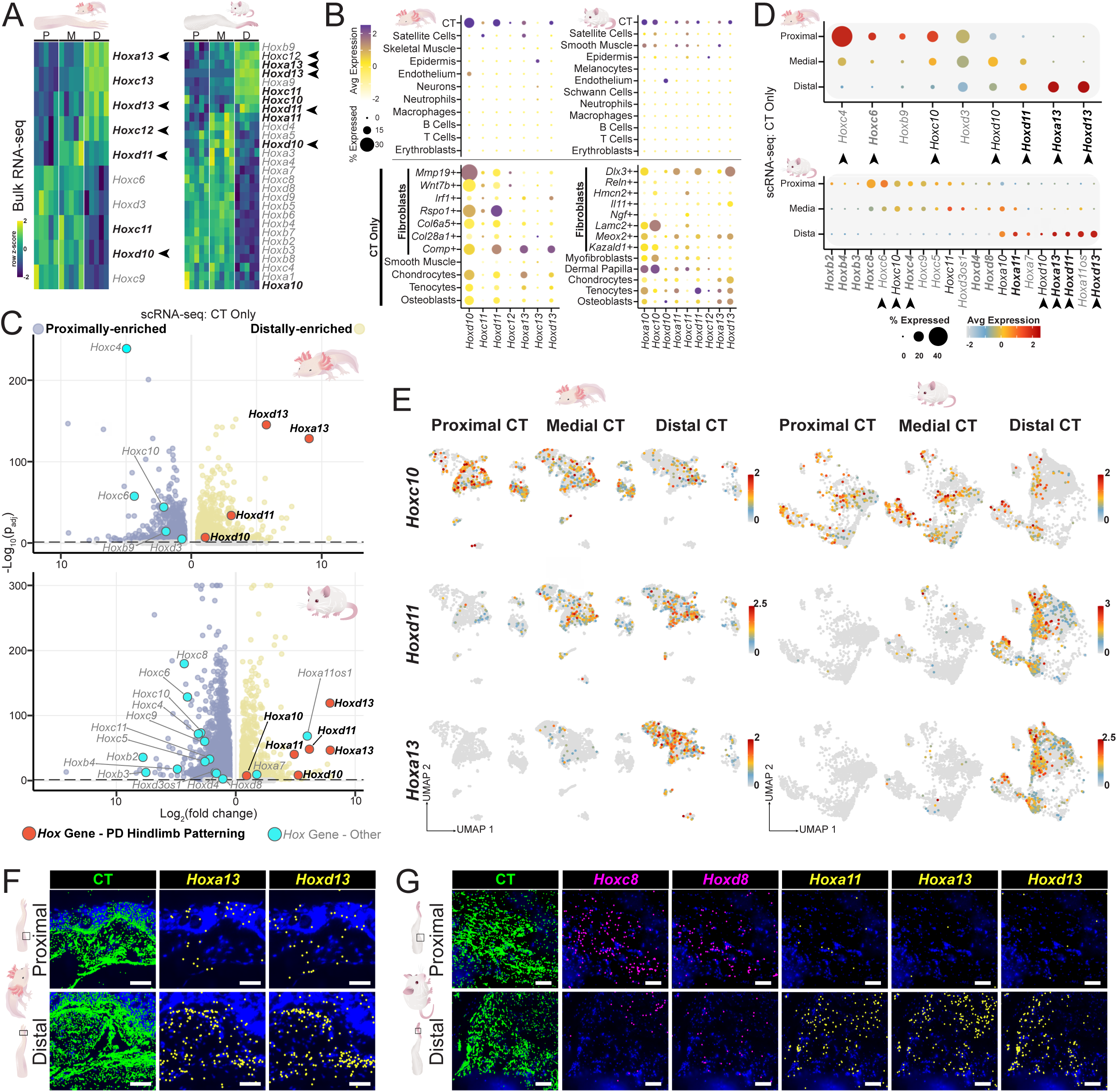
PD expression of *Hox* genes in adult vertebrate limbs. (A) Heatmap of bulk RNA-seq *Hox* gene expression data in skin along the PD axis in the axolotl (left) and mouse (right) hindlimb. All genes shown displayed PD-expression bias in at least one positional comparison (proximal versus distal, proximal versus medial, medial versus distal) by DESeq2, p_adj_<0.05. Bold genes indicate *Hox* genes with known involvement in vertebrate developmental hindlimb PD patterning. Arrows indicate genes with similar expression patterns in both mouse and axolotl datasets. (B) Expression of *Hox* genes with known involvement in vertebrate developmental hindlimb PD patterning that showed adult PD-expression bias within different cell types present in scRNA-seq atlases (top) and within different CT subsets (bottom) for axolotl (left) and mouse (right). (C) Volcano plot depicting top differentially expressed genes between proximal, medial, and distal CT in axolotl (top) and mouse (bottom). For genes that appeared in multiple comparisons (proximal versus distal, proximal versus medial, medial versus distal), only data from the lowest p-value result was used. Dotted line represents cutoff at p_adj_<0.05. (D) Expression of *Hox* genes significantly differentially expressed within CT along the PD axis in axolotl (top) and mouse (bottom) hindlimb CT. Grey backdrop indicates a statistically significant difference in expression in CT from at least two positions along the PD axis (p_adj_<0.05). (E) Expression of *Hoxc10*, *Hoxd11*, and *Hoxa13* within proximal, medial, and distal CT of axolotl (left) and mouse (right) hindlimb CT. (F) MERFISH visualization of *Hoxa13* and *Hoxd13* expression within proximal and distal CT in axolotl hindlimb. (G) MERFISH visualization of *Hoxc8, Hoxd8, Hoxa11*, *Hoxa13,* and *Hoxd13* expression within proximal and distal CT in mouse hindlimbs.CT was visualized via expression of *Pdgfra*, *Dpt*, and *Col6a1* in both organisms. This pool additionally included *Col4a1* in mouse. Scale bars, 100 µm.

In the scRNA-seq CT datasets from both organisms, many of the top differentially expressed genes along the PD axis were *Hox* genes, pointing to a prominence of these genes in the constitutive regional gene expression signature of adult limb CT (Fig. 4C-D, Supplementary Data 7). *Hox* expression patterns within the scRNA-seq CT data were often similar to that of the bulk skin RNA-seq data, e.g., *Hoxd11* expression was medial/distal and *Hoxa13* and *Hoxd13* were expressed distally in the CT of both organisms (Fig. 4C-E). *Hoxa13* and *Hoxd13* also showed prominent distal CT expression by MERFISH in both organisms (Fig. 4F-G, Supplementary Fig. 5B). In mouse, *Hoxa11* displayed distal expression, whereas *Hoxd8* and *Hoxc8* displayed expression in proximal CT by MERFISH, mirroring patterns found in both scRNA-seq and bulk RNA-seq data (Fig. 4G, Supplementary Fig. 5B).

Analysis of the scRNA-seq CT datasets also revealed PD-biased expression of *Hox* genes not identified as PD-biased in bulk skin RNA-seq. For example, *Hoxc4*, *Hoxc6*, and *Hoxc10* were expressed proximally in CT of both organisms; other *Hox* genes showed species-specific patterns, although these typically did not involve canonical limb developmental *Hox* genes (Fig. 4C-E). *Hox* genes from paralog groups 1-9 that showed PD-biased expression in either assay were more often expressed proximally/medially than distally (Fig. 4A-E). These non-canonical limb *Hox* genes with PD-expression bias in the adult limb were also less specifically expressed in CT in both organisms, suggesting that CT-specificity is predominant for *Hox* genes involved in limb patterning during development (Supplementary Fig. 5C). Some of the strongest expression of non-canonical hindlimb *Hox* genes was in the epidermis (e.g., *Hoxa7*, *Hoxa9*), leading us to investigate *Hox* PD-expression differences in epidermal cells from scRNA-seq data.

Epidermal subclustering yielded diverse populations, including various keratinocyte populations in both organisms, hair follicle cells in mouse, and amphibian-specific gland cells^95^ in axolotl (Supplementary Fig. 6A,B, Supplementary Data 8). These cells displayed PD-biased expression of several *Hox* genes, although primarily non-canonical limb *Hox* genes (e.g., *Hoxc13* in axolotl, *Hoxa9* in mouse) (Fig. 4A-B, Supplementary Fig. 5C, Supplementary Fig. 6C-D). The distal-enriched skin expression of some of these *Hox* genes was visible in both species by MERFISH and Visium (Supplementary Fig. 6E). Differentially expressed *Hox* genes on the PD axis were generally expressed in a higher percentage of cells in CT clusters and at higher levels in axolotl than in mouse CT (Fig. 4B-D). Differential expression of other cell types (i.e., smooth muscle, endothelium, satellite cells, Schwann cells) within scRNA-seq datasets for both organisms failed to reveal prominent positionally-biased expression of *Hox* genes, particularly for developmental limb *Hox* genes (Supplementary Data 9, 10).

In summary, the adult limb CT of both organisms harbors a positionally restricted *Hox* gene expression map that largely recapitulates developmental expression patterns. This map is largely specific to limb CT and is distributed throughout CT cell types.

### Many limb developmental patterning TFs show PD-biased expression in adult limbs, but most signaling factors do not

In addition to regional *Hox* gene expression, vertebrate PD limb developmental patterning involves the regional expression of additional signaling factors and TFs. FGF^90,96–106^ and WNT^90,102,105,107–110^ signaling in the apical ectodermal ridge (AER), the ectodermally-derived signaling center at the distal edge of the limb bud^111–113^, drives distal hindlimb outgrowth. FGF expression in the AER is regulated by genes encoding the TFs *Sp6*, *Sp8*, and *Sp9*^114,115^. Axolotls have an unconventional AER, with some FGF and WNT signaling relegated to distal mesenchyme^14,116–119^. Distal outgrowth is partially coordinated with other limb axes by the expression of *Grem1*, which encodes a BMP antagonist, in the distal mesoderm^99,120–122^. Proximal limb identity, both during development and regeneration is regulated by retinoic acid (RA) signaling^90,100,123,124^, with RA synthesized proximally by enzymes encoded by *Aldh1a2*^125,126^ and *Rdh10*^127–129^ and degraded distally by an enzyme encoded by *Cyp26b1*^130–132^. *Meis1* and *Meis2* are RA-responsive TF-encoding genes and are prominently implicated in specifying proximal limb identity, both during development and regeneration^90,99,100,133–136^. Recent evidence suggests that *Meis* may be proximally expressed in axolotl limbs, both in uninjured tissue and in regenerating blastemas^38,132^.We sought to test whether these prominent TFs and signaling components were constitutively expressed in adult vertebrate limbs and if so, in which cell types.

Unlike the *Hox* genes described above, *Wnt* and *Fgf* genes largely showed no overt PD expression bias in bulk RNA-seq skin datasets of both organisms, although some *Wnt* and *Fgf* genes showed proximal/medial enrichment in expression in mouse bulk RNA-seq (Fig. 5A, Supplementary Fig. 7E). In both species, *Wnt* and *Fgf* genes were typically lowly expressed, although some showed epidermal or CT expression in the scRNA-seq data (Supplementary Fig. 7A,D). Some PD-bias in expression existed within CT and epidermis scRNA-seq data, although rarely in ways that mirrored developmental expression patterns (Supplementary Fig. 7B,F-G). In axolotl, *Aldh1a2*, *Rdh10*, and *Cyp26b1* showed no PD-expression bias in the adult limb (Fig. 5A). In mouse, these genes either displayed no significant PD bias or displayed PD bias counter to developmental expectations (Fig. 5A). These findings are generally inconsistent with a model in which major PD limb developmental signaling factors maintain their regional expression as a mechanism of positional memory. This aligns with previous work demonstrating that some key developmental signaling factors are upregulated only during limb regeneration^14,16,137^.

**Figure 5.**
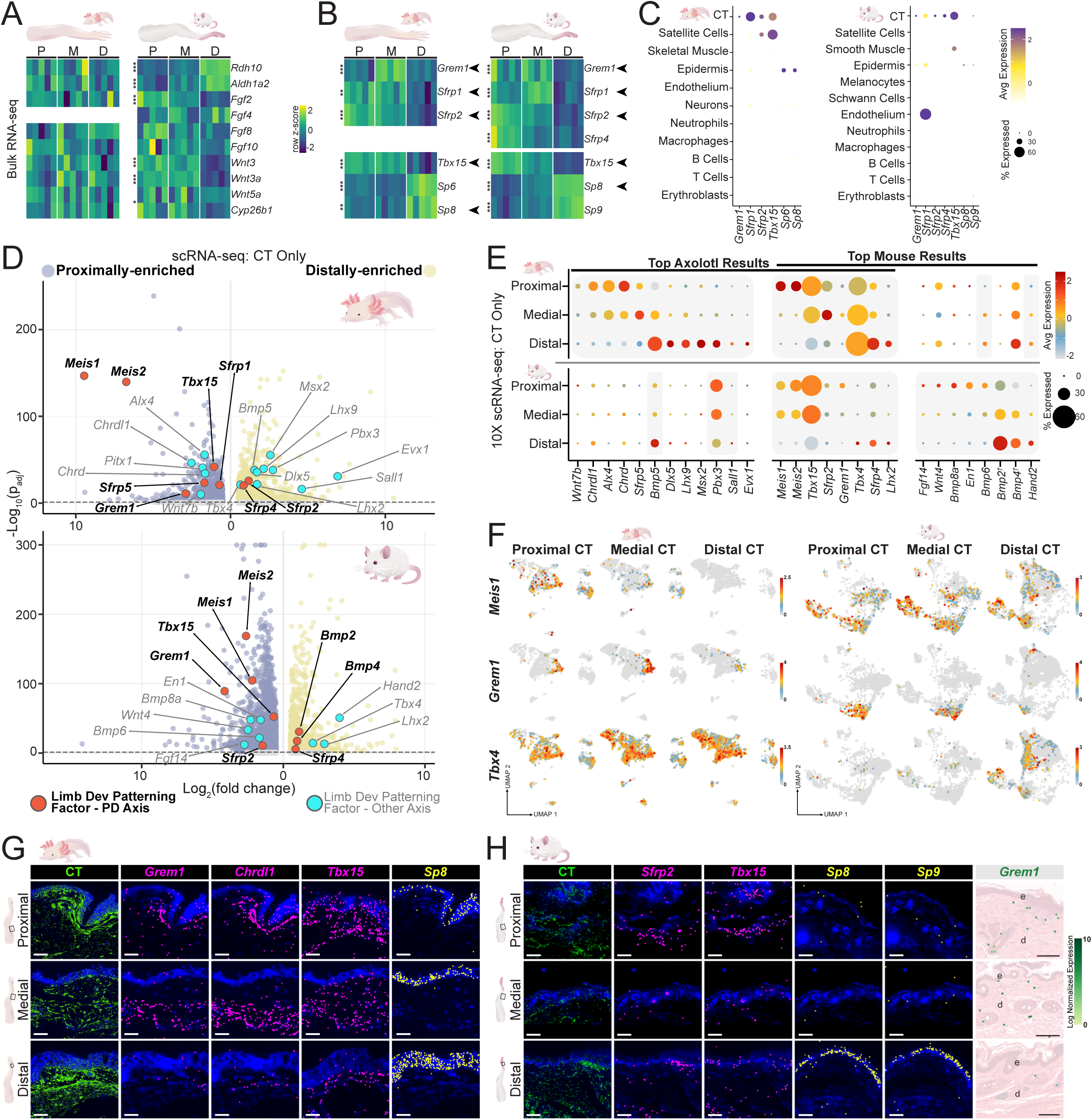
PD expression of developmental patterning factors in adult vertebrate limbs. (A) Heatmap of PD bulk RNA-seq expression of genes encoding signaling factors expressed in the AER or involved in RA-based proximal limb specification in the axolotl (left) and mouse (right) hindlimb. (B) Heatmap of PD bulk RNA-seq expression of signaling factors (top) and TFs (bottom) with roles in limb PD patterning that showed PD-biased expression patterns in axolotl (left) and mouse (right). Arrows indicate genes with similar expression patterns in both mouse and axolotl datasets. (C) Expression of genes from (B) within different cell types present in scRNA-seq atlases for axolotl (left) and mouse (right). (D) Volcano plot depicting top differentially expressed genes between proximal, medial, and distal CT in axolotl (left) and mouse (right). For genes that appeared in multiple comparisons (proximal versus distal, proximal versus medial, medial versus distal), only data from the lowest p-value result was used. Dotted line represents cutoff at p_adj_<0.05. *Hox* genes involved in PD developmental limb patterning are shown in Fig. 4C. . (E) Expression of genes labelled in (D) within CT along the PD axis in axolotl (top) and mouse (bottom) hindlimbs. Grey backdrop indicates a statistically significant difference in expression in CT from at least two positions along the PD axis (p_adj_<0.05). (F) Expression of *Meis1*, *Grem1*, and *Tbx4* within proximal, medial, and distal CT of axolotl (left) and mouse (right) hindlimb CT. (G) MERFISH visualization of *Grem1*, *Chrdl1*, and *Tbx15* in proximal, medial, and distal hindlimb CT as well as expression of *Sp8* within proximal, medial, and distal skin in axolotl hindlimb. (H) MERFISH visualization of *Sfrp2* and *Tbx15* expression in proximal, medial, and distal hindlimb CT as well as expression of *Sp8* and *Sp9* within proximal, medial, and distal skin in mouse hindlimbs. Visium HD visualization of *Grem1* expression in hindlimb dermis (far right). CT was visualized via expression of *Pdgfra*, *Dpt*, and *Col6a1* in both organisms. This pool additionally included *Col4a1* in mouse. e=epidermis, d=dermis. Scale bars, 100 µm. *=p_adj_<0.05, **=p_adj_<0.01, ***=p_adj_<0.001. ^†^=best axolotl BLAST hit.

In both species, multiple members of the *Sfrp* gene family, which encode WNT antagonists with roles in limb developmental patterning^138–140^, displayed proximal/medial expression in bulk RNA-seq skin datasets and were expressed broadly in CT cell types (Fig. 5B-C, Supplementary Fig. 7C). *Sfrp2* was proximally/medially expressed in CT scRNA-seq data in both organisms and was visibly enriched in proximal CT in mouse MERFISH (Fig. 5D-E, H). Additionally, *Grem1* showed distinct proximal/medial expression in the bulk-RNA seq skin data, was expressed primarily in CT, and showed proximal/medial expression in CT scRNA-seq data in both organisms (Fig. 5B-F). The proximal/medial expression bias of *Grem1* in limb CT was also visible by MERFISH in axolotl and with Visium HD in mouse (Fig. 5G-H). Whereas *Sfrp* genes were expressed across many CT populations in both species, *Grem1* was expressed specifically in mouse *Kazald1*+ fibroblasts and axolotl *Rspo1*+ fibroblasts (Fig. 5F, Supplementary Fig. 7C). These genes will be interesting targets for future mechanistic work to ascertain whether they have any functional role in positional memory.

Among TFs, *Meis1* and *Meis2* genes were notable in showing proximal/medial-enriched expression in the scRNA-seq CT data of both organisms (Fig. 5D-F). Interestingly, whereas this observation arose from comparisons within CT cells, *Meis1* and *Meis2* were actually most prominently expressed in the axolotl epidermis; in mouse, these genes were primarily expressed in CT, with minor epidermal expression (Supplementary Fig. 7D). Bulk RNA-seq of skin revealed distal expression of *Sp6* and *Sp8* in axolotl and of *Sp8* and *Sp9* in mouse (Fig. 5B). These genes were expressed primarily in the epidermis in both species (Fig. 5C). Epidermal scRNA-seq data displayed distal expression of *Sp8* in both mouse and axolotl, as well as *Sp9* in mouse (Supplementary Fig. 7F-G). *Sp8* showed clear distal skin-enriched expression in both species by MERFISH, as did *Sp9* in mouse skin (Fig. 5G-H).

We additionally sought to systematically assess other TFs and signaling factors that have less well-characterized roles in developmental limb PD axis formation. We found several such genes with PD-biased expression in bulk RNA-seq skin and scRNA-seq datasets, many with CT-enriched expression (Fig. 5D-F, Supplementary Fig. 7C-F).

*Tbx15* is essential for proper proximal limb formation in development^141^. *Tbx15* showed proximal/medial expression in bulk RNA-seq datasets, CT scRNA-seq datasets, and by MERFISH in both organisms (Fig. 5B-E, G-H). *Chrdl1*, which encodes a secreted BMP antagonist, has been implicated in limb patterning^142^ and was previously shown to be expressed proximally in the adult axolotl limb expression in bulk RNA-seq data for both mouse and axolotl and was expressed in the CT of both organisms (Supplementary Fig. 7D-E). *Chrdl1* also displayed proximal/medial expression in axolotl scRNA-seq CT data and with MERFISH (Fig. 5D-E,G). Differential expression analysis with the scRNA-seq datasets failed to reveal prominent PD-expression bias for most of these limb-developmental PD-patterning factors in other non-CT cell types (Supplementary Data 9, 10).

In summary, expression of core limb PD-patterning signaling components – *Fgf*, *Wnt*, and RA – did not mirror their developmental roles in the adult limbs of either organism. However, several other genes associated with limb development – including the TFs *Meis1, Meis2, Sp8*, and *Tbx15*, as well as genes encoding BMP- (*Grem1*, *Chrdl1*) and WNT-related signaling factors (*Sfrp2*) – displayed CT-driven PD-biased expression patterns, in many cases recapitulating expression during development and regeneration. Several of these were shared between the regenerative and non-regenerative vertebrate limbs.

### Limb atlases reveal PD-biased expression signatures of TFs, signaling, and ECM-modifying factors

In principle, genes involved in adult limb positional memory need not have roles in limb development. We therefore characterized PD-biased expression for additional genes across the genome, regardless of any prior linkage to limb development. We ranked each gene that showed significant PD-expression bias by the padj-value of differential expression in PD bulk skin RNA-seq and isolated CT scRNA-seq datasets. Many of the top-ranked genes in both datasets were the limb-developmental factors that were discussed above; these genes were used to set rank thresholds of interest for each species (Supplementary Fig. 8A). We first identified high-ranking genes in each species that were predicted to be involved in processes that could influence patterning processes – signaling, ECM modification, or transcriptional regulation – or were genes known to have a limb-development role, but are less well characterized than those described above (Fig. 6A-C). Some genes that met these criteria showed similar expression patterns in both species, although the expression patterns of more than half of the identified genes were species-specific (Fig. 6A-C). The expression of identified genes was most often primarily in CT in both species (Supplementary Fig. 8B). *Tgfbr2*, which encodes a TGF-β receptor, displayed proximal expression in axolotl and mouse skin bulk RNA-seq, in axolotl scRNA-seq CT data, and also in axolotl MERFISH and mouse Visium HD spatial data (Fig. 6A-D, Supplementary Fig. 8C). *Hpse2*, which encodes an ECM-remodeling factor, showed distal expression bias in mouse bulk RNA-seq data, in CT scRNA-seq data for both species, as well as in distal CT in axolotl MERFISH and mouse Visium HD data (Fig. 6A-D, Supplementary Fig. 8C). Several other high-ranking genes in each species displayed similar expression patterns that mirrored sequencing results in axolotl MERFISH and mouse Visium HD spatial data (e.g., *Ntn1*, *Pianp, Tll2* in axolotl and *Adamts2, Trps1,* and *Rbp7* in mouse), typically specific to either CT or epidermis (Fig. 6D, Supplementary Fig. 8C-D). These data demonstrate that, in addition to genes with roles in limb developmental patterning and differentiated epidermal function, many additional genes involved in signaling, ECM modification, or transcriptional regulation have regional PD expression within adult axolotl and mouse limbs.

**Figure 6.**
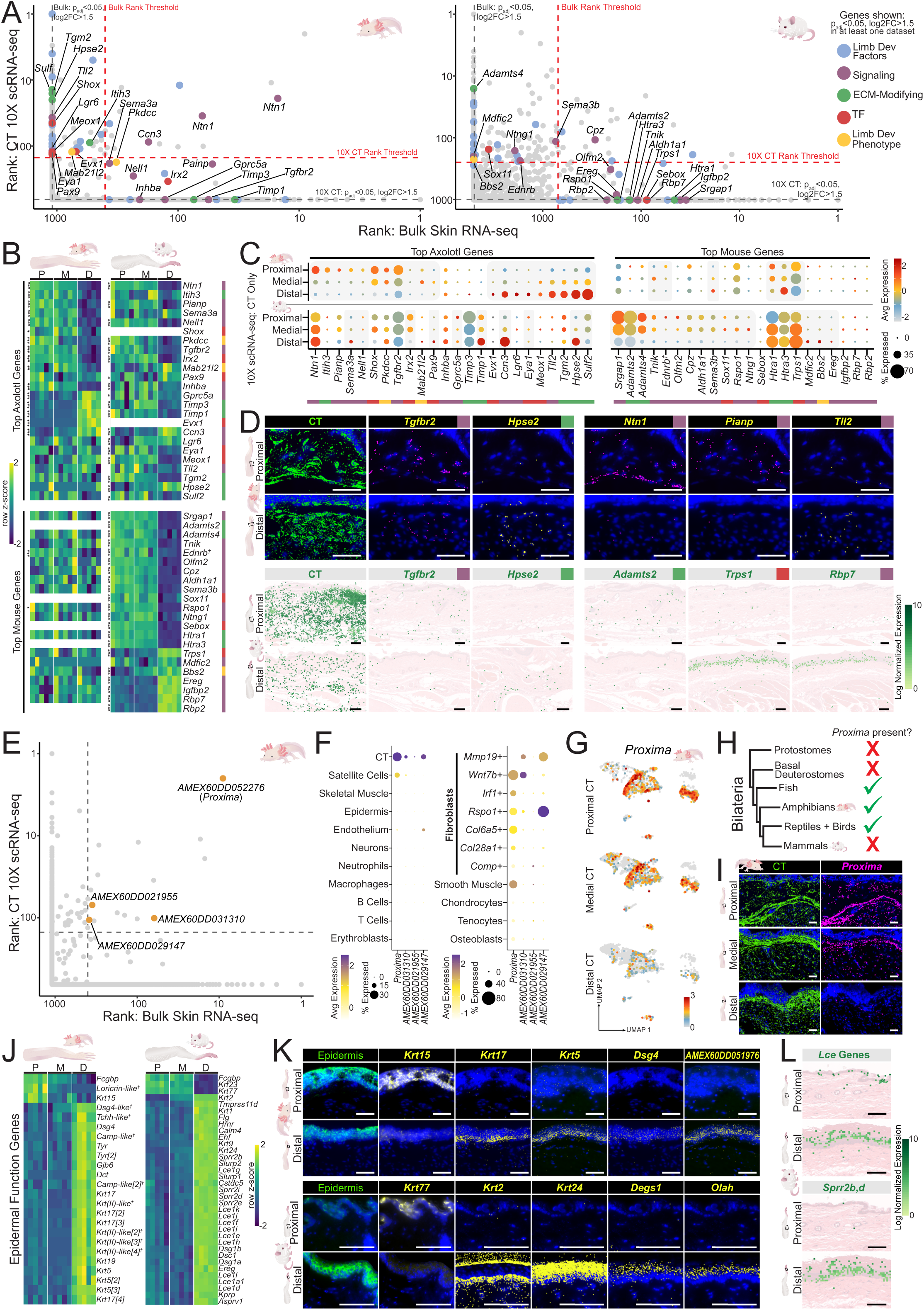
Genome-wide characterization of PD-biased expression across vertebrate hindlimbs. (A) Plot displaying the rank of each gene in differential expression analysis in the bulk skin RNA-seq dataset (x-axis) and CT 10X scRNA-seq dataset (y-axis) for axolotl (left) and mouse (right) hindlimbs. Genes were sorted by p_adj_ and for genes that appeared in multiple comparisons (proximal versus distal, proximal versus medial, medial versus distal); only data from the lowest p-value result was used. Genes were only included if p_adj_<0.05 and log_2_FC>1.5 for at least one positional comparison. Dotted line represents rank thresholds for identifying genes of interest for positional information roles, set by ranks of discussed developmental factors (axolotl: x-axis intercept = 400, y-axis intercept =150; mouse: x-axis intercept = 700; y-axis intercept=250), shown in Supplementary Fig. 8A and see text. Genes of certain functional categories are labelled, shown in the key (far right). (B) Heatmap of bulk skin RNA-seq expression of genes from (A) along the PD axis in axolotl (left) and mouse (right) hindlimbs. (C) Expression of select genes from (A) in axolotl (top) and mouse (bottom) hindlimb CT. Grey backdrop indicates a statistically significant difference in expression in CT from at least two positions along the PD axis (p_adj_<0.05). (D) MERFISH visualization of select genes from (A) in axolotl (top) and mouse (bottom) proximal and distal limb tissue. *Tgfbr2* and *Hpse2* patterns are shared between organisms, whereas other patterns are species specific. CT marked by *Pdgfra*, *Dpt*, and *Col6a1* in axolotl. CT in mouse Visium HD marked by *Pdgfra*, *Col3a1*, *Dpt*, *Col4a1*, *Dcn*, *Col15a1*, *Pi16*, *Tagln*, and *Acta2*. (E) Axolotl rank plot from (A) with top unannotated genes labelled. (F) Expression of genes from (E) within different cell types present in scRNA-seq atlas (left) and CT subsets (right) in axolotl limb tissue. (G) UMAP representation of CT from proximal, medial, and distal axolotl CT showing *Proxima* expression. (H) Presence or absence of *Proxima* orthologs in bilaterian clades; for a more complete phylogeny, see Supplementary Fig. 9F. (I) MERFISH visualization of *Proxima* expression in proximal, medial, and distal axolotl CT. CT marked by *Pdgfra*, *Dpt*, and *Col6a1.* (J) Heatmap of bulk skin RNA-seq expression of highly differentially expressed genes with predicted differentiated epidermal function along the PD axis in the axolotl (left) and mouse (right) hindlimb. (K) MERFISH visualization of expression of highly differentially expressed genes along the PD axis in proximal and distal axolotl (top) and mouse (bottom) skin. Epidermis is marked by *Lgals7* in mouse and *Scel* in axolotl. (L) Visium HD visualization of *Lce* (*Lce1a1*, *Lce1d*, *Lce1e, Lce1f, Lce1k,* and *Lce1l* are shown) (top) and *Sprr* (bottom) genes in proximal and distal mouse skin. All scale bars are 100 µm. The colored box next to gene names corresponds to the functional category labelled in (A). *=p_adj_<0.05, **=p_adj_<0.01, ***=p_adj_<0.001. ^†^=best axolotl BLAST hit.

### *Proxima* is a vertebrate-specific gene, lost in mammals, with PD expression bias in axolotl

Given the prominent role of salamander-specific gene *Prod1* in axolotl limb patterning^13,143–145^, we sought to expand our search for positional information factors to genes that encoded uncharacterized or novel proteins that displayed PD-enriched expression. Four unannotated axolotl genes were ranked above the established thresholds for both bulk RNA-seq and scRNA-seq CT differential expression (Fig. 6E, Supplementary Fig. 9, Supplementary Data 11). Of these uncharacterized axolotl genes, *AMEX60DD052276* stood out as the gene with the single most significantly PD-biased expression when considering rank in both datasets, outranking all previously discussed limb development (e.g., *Hox*) genes (Fig. 6E). *AMEX60DD052276* displayed proximal/medial expression bias and was primarily expressed in fibroblast and other CT cell types (Fig. 6F-G, Supplementary Fig. 9A-D). *AMEX60DD052276* encodes a protein with no recognizable domains but that is predicted to have a signal peptide and to be secreted (Supplementary Fig. 9E). We named this gene *Proxima*. Orthologs of *Proxima* were present in the genomes of representative species from all major vertebrate clades, but absent in the genomes of all mammals and non-vertebrate clades (Fig. 6H, Supplementary Fig. 9F, Supplementary Data 11). This suggests that *Proxima* emerged with vertebrates and remained broadly conserved, with the notable loss of this gene in the evolution of mammals. Alignments of the predicted structures encoded by the orthologs of *Proxima* from different vertebrate clades suggest the existence of a conserved C-terminal domain and N-terminal structural features that are missing in the basal vertebrate (lamprey) ortholog (Supplementary Fig. 9G). In MERFISH data, *Proxima* displayed clear proximal/medial expression enrichment, with expression particularly notable in dermal CT (Fig. 6I).

### PD specialization of the epidermis

We also sought to characterize PD-biased expression more broadly in limb tissue, without specific focus on CT. Among the top 200 differentially expressed genes in the skin bulk RNA-seq datasets of both organisms, a large fraction of the genes were expressed distally in epidermis and are predicted to have roles in mature epidermal function (Supplementary Fig. 10A). Both axolotl and mouse distal limb skin expressed several specific keratin-encoding genes and genes encoding structural and anti-microbial peptides (*Tchh*-like^146^ and *Camp*-like^147^ genes in axolotl, *Lce*^148,149^ and *Sprr*^150^ genes in mouse) (Fig. 6J). MERFISH showed both proximal- and distal-specific expression of a subset of these genes in the skin of axolotl and mouse (Fig. 6K, Supplementary Fig. 10B). MERFISH also showed distal-enriched expression of lipid metabolism genes *Olah* and *Degs1* in mouse skin and of a transcript *AMEX60DD051976*, which is annotated as an uncharacterized lncRNA in axolotl (Fig. 6K, Supplementary Fig. 10B). Visium HD data displayed distal expression of various *Lce* and *Sprr* genes in mouse skin (Fig. 6L, Supplementary Fig. 10B). We also analyzed isolated epidermis in the scRNA-seq data to identify other genes that displayed PD-expression bias within limb skin. This analysis revealed new expression patterns in mouse skin that were also visualized with Visium HD (Supplementary Fig. 10C).

Additionally, many of the epidermal function genes identified in bulk skin RNA-seq data in both organisms showed similar expression patterns in epidermis scRNA-seq data, often being expressed more in keratinocytes than in other epidermal populations (Supplementary Fig. 10D-E). Computational analysis of the Visium HD dataset enabled the discovery of yet more genes with clear PD-biased expression within mouse epidermis, including an *Lce* gene with strong proximal enrichment (Supplementary Fig. 10F, Supplementary Data 5). Previously, few genes had been shown to be specifically expressed in hand/foot skin previously in mammals^22,23^; this work reveals a spatial atlas of specialized limb epidermis across species.

Several distal epidermal markers in axolotl, when visualized by MERFISH, localized to a specific patch of skin on the anterior, ventral epidermis in both fore and hindlimb (Supplementary Fig. 11A). These markers were also co-expressed in a small epidermal subcluster in the scRNA-seq dataset, suggesting that this skin patch constitutes a distinct skin cell state (Supplementary Fig. 11B-C). These cells showed highly enriched expression of genes shown to be specifically expressed to the nuptial pad (a specialized patch of epidermis that some male amphibians develop associated with breeding^151^) in frog samples^152^ (Supplementary Fig. 11C). To our knowledge, this is the first description of a nuptial pad-like cell state in axolotl. Of note, this domain has spatial specificity in all three limb axes, present only in distal, anterior, ventral axolotl skin – positional specificity that could prove quite useful for characterizing regeneration-patterning phenotypes in future work.

### DV limb developmental factor expression is more pronounced in axolotl limb tissue than in mouse

In addition to the proximal-distal axis, tissues are organized asymmetrically on the DV and AP vertebrate limb axes, with positional information of these axes essential for regeneration^14,16,153–156^. In development, dorsal limb identity is regulated by *Lmx1b* and *Wnt7a*^157–161^, whereas ventral identity is regulated by *En1* and *Bmp*^160,162–165^. We identified the top differentially and constitutively expressed genes between adult dorsal and ventral CT in each organism (Fig. 7A, Supplementary Data 12). *Wnt7a* and *Bmp* genes were generally not among the top differentially expressed genes in either organism, similar to the results with signaling factors for the PD axis (Fig. 7A, Supplementary Data 12). By contrast, axolotl *Lmx1b* was prominently expressed in dorsal CT, recapitulating its developmental expression domain (Fig. 7A-C). MERFISH showed notably higher expression of *Lmx1b* in dorsal axolotl CT than in ventral axolotl CT, though *Lmx1b* was also expressed uniformly in epidermis, consistent with sequencing results (Fig. 7B,D). Some evidence of *Lmx1b* expression in uninjured dorsal axolotl skin has been previously reported *Lmx1b* is expressed specifically in the dorsal nail-associated mesenchyme of the mouse digit tip^34,167^.

**Figure 7.**
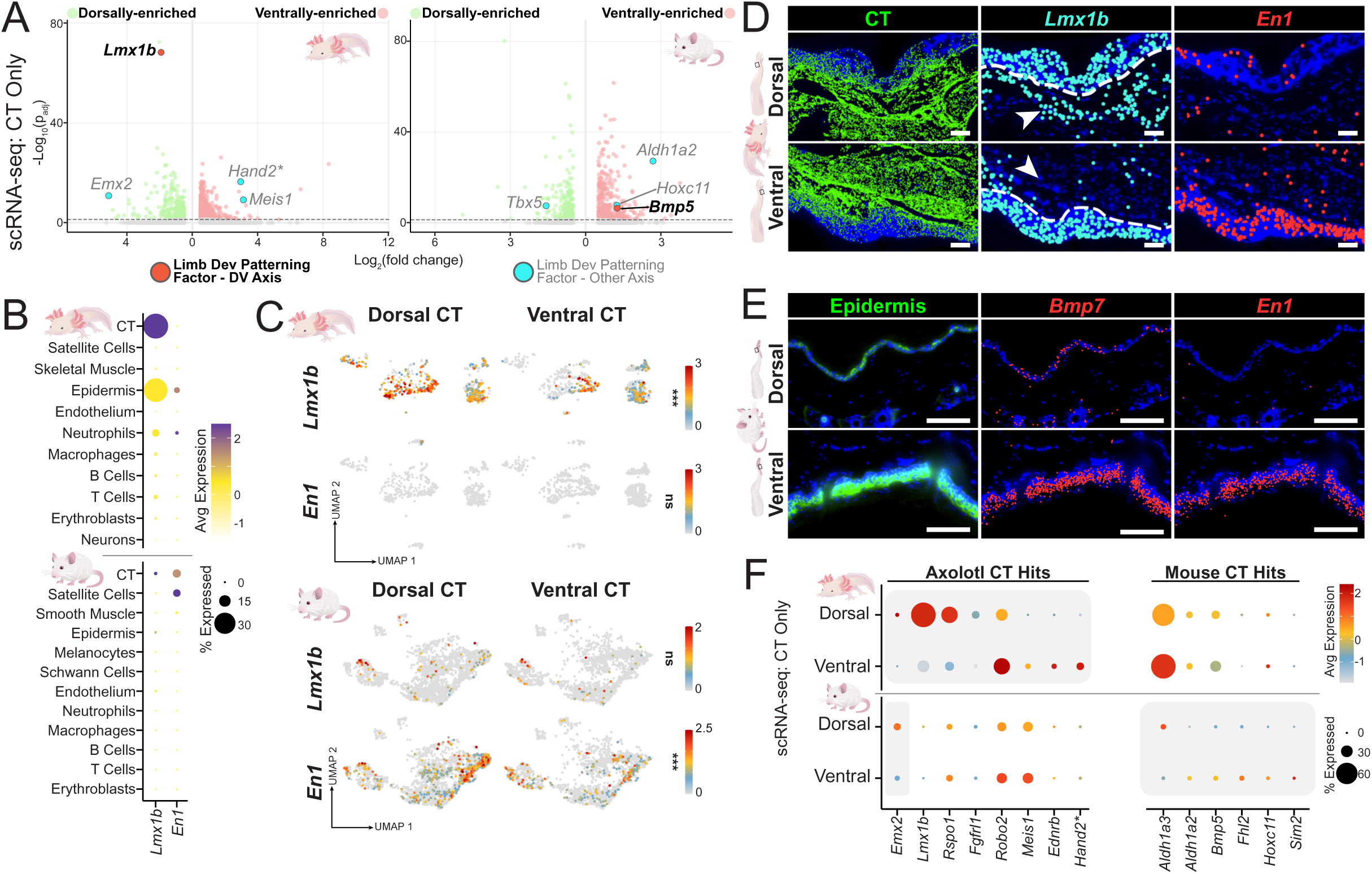
Characterization of DV expression patterns in vertebrate limbs. (A) Volcano plot depicting top differentially expressed genes between dorsal and ventral CT in axolotl (left) and mouse (right). Dotted line represents cutoff atp_adj_<0.05. (B) Expression of genes with known roles in DV-limb developmental patterning within different cell types present in scRNA-seq atlases for axolotl (top) and mouse (bottom). (C) UMAP visualization of expression of *Lmx1b* and *En1* in axolotl (top) and mouse (bottom) CT. (D) MERFISH visualization of *En1* and *Lmx1b* in dorsal and ventral axolotl distal hindlimb tissue. CT marked by *Pdgfra*, *Dpt*, and *Col6a1.* Dotted white line divides epidermis from dermal CT. Arrowheads point to dorsal (top) and ventral (bottom) CT, which have different levels of *Lmx1b* expression. (E) MERFISH visualization of *En1* and *Bmp7* in dorsal and ventral mouse distal hindlimb tissue. (F) Expression of genes with connection to developmental limb patterning that show DV-biased expression within CT along the DV axis in axolotl (top) and mouse (bottom) hindlimb CT. Grey backdrop indicates a statistically significant difference in expression in CT from at least two positions along the PD axis (p_adj_<0.05). Scale bars, 100 µm. The axolotl *Hand2\** locus refers to a distinct gene in the axolotl genome assembly from the canonical *Hand2* referenced in other figures (see Supplementary Data 16).

*En1* showed no clear recapitulation of developmental pattern in adult CT in both organisms, being largely absent in axolotl CT and present in mouse CT, but not ventrally biased in expression (Fig. 7B-C). However, mouse bulk RNA-seq data – generated by separately collecting tissue (including skin, skeletal muscle, and deeper fascia) from the dorsal and ventral limb (see Methods) – did show ventral-specific *En1* expression (Supplementary Fig. 12A). This pattern was also robust in MERFISH and Visium HD data, which showed ventral-specific *En1* expression in the epidermis of both axolotl and mouse (Fig. 7D-E, Supplementary Fig. 12B). Mouse ventral epidermis also specifically expressed *Bmp7* by MERFISH and Visium HD (Fig. 7E, Supplementary Fig. 12B). These data suggest that ventral skin in both organisms expresses markers of developmental ventral identity.

To broaden our search for DV positional information candidate genes, we sought all genes with DV-biased expression and that have predicted roles in signaling, ECM modification, or transcriptional regulation (Fig. 7F). The expression patterns observed were rarely shared between species, although these genes were often specifically expressed in CT and widely among CT subtypes in both organisms (Fig. 7F, Supplementary Fig. 12C). These data demonstrate that, whereas skin in both organisms shows some DV-biased gene expression of limb DV developmental factors, only axolotl limb dorsal CT displayed *Lmx1b* expression, recapitulating developmental pattern.

### AP-patterning factors recapitulate developmental expression patterns in axolotl, but not mouse, CT

During limb development, anterior identity is regulated by *Alx4*^168–171^ and posterior identity is regulated by *Shh*^172–176^ signaling and the TF gene *Hand2*^16,177–179^. Certain Hox genes can also contribute to AP pattern, especially of digits in the limb autopod, with *Hoxd13* and *Hoxa13* promoting posterior identity^87,180^. *Hand2* has recently been characterized as maintaining posterior identity in adult axolotl limb tissue and is essential for proper regeneration, with *Alx4* and *Hand2* reporters showing anterior and posterior limb expression, respectively^16^. We identified the top differentially expressed genes between anterior and posterior CT in both organisms (Fig. 8A, Supplementary Data 13). In axolotls, *Shh* is known to be upregulated substantially during regeneration, with expression being low or absent prior to injury^14,16,181,182^, which was consistent with our findings (Fig. 8A, Supplementary Data 13). By contrast, axolotl *Hand2* and *Alx4* recapitulated their developmental expression bias, displaying posterior and anterior enrichment in expression, respectively (Fig. 8A-C). Other TFs noted to be differentially expressed across axolotl AP CT^16^ also displayed AP expression bias in our scRNA-seq datasets: *Hoxd13* showed posterior expression, and *Alx1*, *Lhx2*, and *Lhx9* displayed anterior expression. Each of these genes were expressed specifically and broadly in axolotl CT, with the exception of *Alx1* which was fairly specific to *Rspo1*+ fibroblasts (Fig. 8B-C, Supplementary Fig. 13B). In mouse, only *Hand2* displayed CT-specific expression among these genes (Fig. 8B-C, Supplementary Fig. 13B). However, *Hand2* expression was not clearly AP-biased in mouse; whether its CT expression reflects any element of AP positional memory remains uncertain and, if present, its significance is not clearly suggested by available data. In axolotl, posterior-enriched expression of *Hand2* and *Hoxd13* and anterior-enriched expression of *Alx1*, *Lhx2*, and *Lhx9* was broad across CT types by MERFISH (Fig. 8D, Supplementary Fig. 13A). Although AP-biased expression of these genes has been previously shown^16^, their specificity and distribution within limb tissue had not been previously characterized. *Grem1* has also been shown to have posterior-biased expression in axolotl limb CT^16^. Whereas *Grem1* was generally less expressed distally, as previously discussed, its distal expression in axolotl was posteriorly-biased by MERFISH (Supplementary Fig. 13A). *Hoxd11* was also posteriorly-biased in expression by MERFISH, consistent with our sequencing results (Fig. 8A, D). With the exception of *Hoxd11* and *Hoxd13*, none of these genes were found to have significant AP bias in mouse CT (Fig. 8A-C, E).

**Figure 8.**
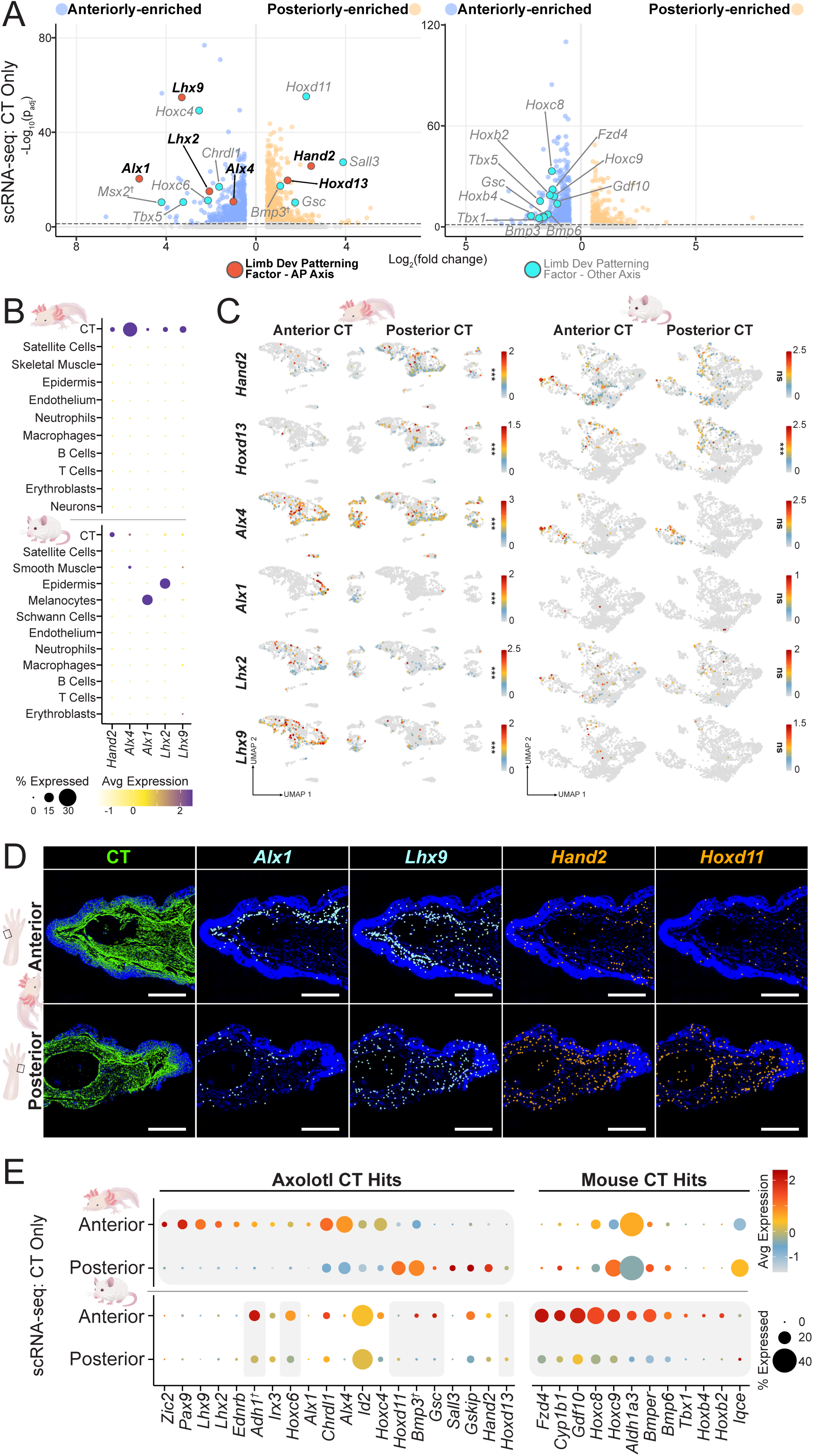
Characterization of AP expression patterns in vertebrate limbs. (A) Volcano plot depicting top differentially expressed genes between anterior and posterior CT in axolotl (left) and mouse (right). Dotted line represents cutoff at p_adj_<0.05. (B) Expression of genes with known roles in AP-limb patterning within different cell types present in scRNA-seq atlases for axolotl (top) and mouse (bottom). (C) UMAP visualization of expression of genes from (B) in axolotl (left) and mouse (right) CT. (D) MERFISH visualization of *Alx1, Lhx9, Hand2, and Hoxd11* in anterior and posterior distal hindlimb tissue. CT marked by *Pdgfra*, *Dpt*, and *Col6a1.* (E) Expression of genes with connection to developmental limb patterning that show AP-biased expression within CT along the AP axis in axolotl (top) and mouse (bottom) hindlimb CT. Grey backdrop indicates a statistically significant difference in expression in CT from at least two positions along the PD axis (p_adj_<0.05). Scale bars, 500 µm. ^†^=best axolotl BLAST hit.

We sought to identify all genes with AP-biased expression that encode proteins with predicted roles in signaling, ECM modification, or transcriptional regulation in both species (Fig. 8E), as well as genes that encode uncharacterized or novel proteins (Supplementary Data 14). The identified genes were often expressed in CT in both species, but infrequently shared expression patterns between species (Fig. 8E, Supplementary Fig. 13B). These findings demonstrate that axolotl CT displays strong signatures of AP positional identity that mirror development, whereas these signatures appear to be either vestigial or altogether absent in mouse CT.

### Spatial expression patterns are reestablished during regeneration

Having characterized spatial gene expression across the developmental axes of uninjured vertebrate limbs, we next sought to understand how these genes are expressed during regeneration. Axolotl limbs that had regenerated, but not fully grown to final scale – taken at 58 and 68 days post-amputation (dpa) – allowed imaging of full, patterned limbs across every major axis using MERFISH (Fig. 9A). The tissue of the newly regenerated axolotl limb proved to be highly compatible with hybridization-based visualization of gene expression, a phenomenon also observed in newly regenerated tissue of other organisms^183^. Position-specific gene expression was robustly reestablished and a multigene, patterned expression map of a regenerative vertebrate limb was visualized (Fig. 9A).

**Figure 9.**
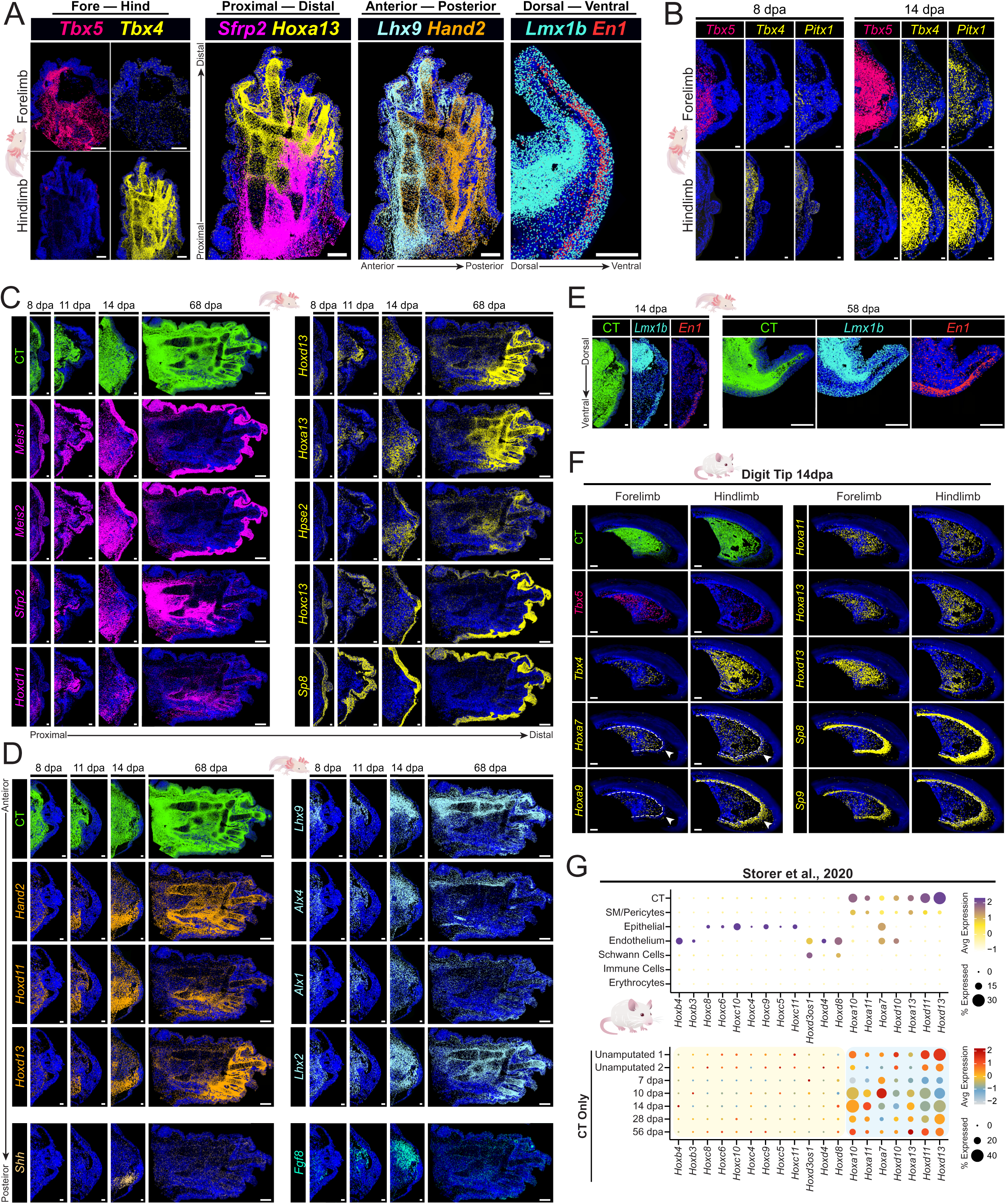
Positionally restricted gene expression during and after regeneration. (A) MERFISH data for genes with positionally restricted gene expression patterns representing the limb positional axes in adult regenerated axolotl limbs. All images are from 68 dpa except the dorsal-ventral image, which is from 58 dpa. (B) MERFISH visualization of *Tbx4* and *Tbx5* expression in early juvenile forelimb and hindlimb blastemas. (C) MERFISH data for genes with PD-biased expression in uninjured tissue during and after axolotl hindlimb regeneration. 68 dpa images for *Hoxa13* and *Sfrp2* are identical to that shown in (A), but in single-channel form. (D) MERFISH data for genes with AP-biased expression in uninjured tissue during and after axolotl limb regeneration. 8, 14, and 11 dpa images are of forelimb blastemas, whereas 68 dpa images are from regenerated hindlimbs. *Hoxd11* and *Hoxd13* 68 dpa images are identical to those shown in (C), recolored to emphasize posteriorly-biased expression. (E) MERFISH data for *Lmx1b* and *En1* during and after axolotl limb regeneration. Both 14 dpa and 58 dpa images are of forelimbs. (F) MERFISH data for genes with positionally restricted expression in uninjured mouse tissue in 14 dpa regenerating forelimb and hindlimb mouse digit tips. Dotted white line divides epidermis from dermal CT for genes with epidermal expression. Arrowheads point to epidermal gene expression that differs between forelimb and hindlimb regenerating digits. (G) Expression of *Hox* genes that displayed PD-biased expression in uninjured mouse CT within different cell types (top) and CT from different regenerative time points (bottom) present in the scRNA-seq mouse digit tip regeneration datasets from Storer et al., 2020^33^. Images of 58 and 68 dpa axolotl limbs are from adult axolotls. Images of 8 dpa and 11 dpa axolotl limbs are from juvenile regenerating axolotls.CT was visualized via expression of *Pdgfra*, *Dpt*, and *Col6a1* in both organisms. This pool additionally included *Col4a1* in mouse. Scale bars, 100 µm for 8 dpa and 11 dpa axolotl images and 500 µm for 58 dpa and 68 dpa axolotl images. Scale bars 100 µm for mouse 14 dpa images.

Differences in CT expression between forelimb and hindlimb that were visible in the intact limb — forelimb-specific *Tbx5* expression and hindlimb-specific *Tbx4*, *Pitx1*, and *Hoxc10* expression — were robustly present after regeneration. These patterns were present in blastemas as early as 8 dpa and were maintained throughout the course of regeneration (Fig. 9A-B, Supplementary Fig. 14A).

For the PD axis, regenerated limbs displayed overlapping but distinct expression patterns for previously visualized genes with distal expression bias in CT, including for *Hoxa13*, *Hoxd13*, and *Hpse2* (Fig. 9A,C, Supplementary Fig. 14B). This approach also enabled visualization of expression for some other genes that showed PD-biased expression within CT in the sequencing results, including proximally-biased expression of *Meis1*, *Meis2*, *Hoxd11,* and *Sfrp2* (Fig. 9A,C, Supplementary Fig. 14B). The *Meis* genes also displayed strong uniform expression in limb epidermis, consistent with sequencing results (Supplementary Fig. 7D). Genes with distal expression in the epidermis, such as *Sp8* and *Hoxc13,* also displayed clear positionally-biased expression along the full regenerated PD limb axis (Fig. 9A,C, Supplementary Fig. 14B). Many of these expression patterns were poorly resolved at 8 dpa, although some showed PD-biased expression by 11 dpa (e.g., *Sfrp2*, *Hoxd13*, *Hoxa13*).

The AP axis was also visualized with single-cell spatial resolution across the full regenerated PD limb axis. *Hand2*, *Hoxd11*, and *Hoxd13*, which were expressed posteriorly in intact limb sequencing data, displayed distinct but overlapping posterior expression domains within CT (Fig. 8A, Fig. 9A,D, Supplementary Fig. 14C).

Conversely, *Lhx9*, *Alx4*, *Alx1*, and *Lhx2,* genes that were expressed anteriorly in limb sequencing data, showed distinct but overlapping expression domains in regenerated anterior CT (Fig. 8, Fig. 9A,D, Supplementary Fig. 14C). These AP-biased gene expression patterns were typically identifiable as early as 8 dpa (Fig. 9D). In contrast to the TFs that displayed perduring AP-biased expression throughout regeneration, *Shh* and *Fgf8* – signaling factors known to be important for specifying posterior and anterior limb tissue, respectively^14^ – displayed more transient AP-biased expression, with the strongest expression signature observed at 14dpa and absent at 68dpa (Fig. 9D, Supplementary Fig. 14C). *Fgf8* appeared to be expressed earlier than *Shh*, with *Shh* expression lasting longer (still visible at 58dpa), consistent with prior experiments quantifying transcript levels^14^ (Fig. 9D, Supplementary Fig. 14C).

For the DV axis, *Lmx1b* was expressed more in dorsal CT than in ventral CT in the regenerated limb, consistent with data from uninjured axolotl limbs (Fig. 9A,E). Similarly, *En1* displayed higher expression in regenerated ventral epidermis than regenerated dorsal epidermis, recapitulating the expression pattern in the uninjured tissue (Fig. 9A,E). These expression patterns were observable at 14dpa (Fig. 9E).

Although mice are incapable of limb regeneration, they can regenerate distal digit tips^184–186^. We assessed whether regenerating digit tips maintain forelimb- and hindlimb-specific gene expression and whether they express distal-biased limb genes. *Tbx5* was more strongly expressed in regenerating forelimb than in regenerating hindlimb digit tip CT, consistent with expression patterns in uninjured mouse tissue (Fig. 9F). Notably, mouse *Tbx4*, which did not show a clear expression bias between uninjured limb types, did display stronger expression in the regenerating hindlimb digit tip than in the regenerating forelimb digit tip, specifically in CT, strengthening the evidence for fore-versus-hindlimb positional memory in mice (Fig. 9F). *Hoxa7* and *Hoxa9*, which were specifically expressed in uninjured hindlimb paw epidermis (Supplementary Fig. 3E-F) showed hindlimb-specific expression in regenerating mouse digit tip epidermis, consistent with a distinct hindlimb positional memory that is retained during and regeneration (Fig. 9F). *Hoxa13* and *Hoxd13*, which were expressed distally in uninjured mouse CT, were expressed in regenerating digit tip CT (Fig. 9F). *Sp8*, *Sp9*, *Hoxc11*, *Hoxc12*, as well as several epidermal function genes with robust distal expression in uninjured epidermis, also displayed expression in regenerating digit tip epidermis (Fig. 9F, Supplementary Fig. 14D).

To further corroborate the observation that regenerating mouse digit tips expressed genes with distal-biased expression in uninjured tissue, we assessed two independent mouse regenerating digit tip scRNA-seq datasets^32,33^ (Supplementary Fig. 15A,D). In both datasets, *Hox* genes with proximal expression in uninjured CT did not show notable expression in regenerating digit tip CT, whereas *Hox* genes with distal expression in uninjured CT showed robust expression in regenerating CT across regeneration timepoints (Fig. 9G, Supplementary Fig. 15E). Similarly, non-*Hox* developmental factors that displayed distal expression in uninjured CT showed higher expression in regenerating digit tip CT than factors that had proximal expression in uninjured CT across timepoints (Supplementary Fig. 15B,F). Regenerating epithelial tissue in these datasets also expressed several genes we found to be expressed specifically in distal uninjured skin in sequencing data (Supplementary Fig. 15C,G).

## Discussion

The underlying properties that distinguish regenerative and non-regenerative organisms remain a fundamental problem. Positional information that guides the identity and pattern of new tissue to precisely replace lost tissue can be essential for regeneration.

Constitutively active and regional adult gene expression can constitute positional information and is sometimes referred to as positional memory. The positional information landscapes in regenerative species remain poorly characterized outside of select species and the identity and roles of cell types responsible for harboring this information are poorly understood. In principle, differences in the presence or makeup of adult positional information could underlie evolved differences in regenerative capacity.

We comprehensively characterized constitutive, positionally biased gene expression across all adult limb axes of mouse and axolotl, constituting positional atlases of gene expression for both regenerative and non-regenerative vertebrate limbs. Numerous key components of positional information in development reprised their expression domains in adult tissue in both organisms. Prominently, *Tbx5* was expressed in the forelimb and *Pitx1* and *Hoxc10* in the hindlimb of both organisms. Axolotls additionally displayed hindlimb-specific *Tbx4* expression. Mice did not overtly display hindlimb-specific *Tbx4* expression in the uninjured limb, but stronger *Tbx4* expression was seen in regenerating hindlimb digit tips compared to those from the forelimb. Expression of these factors was predominantly in the diverse fibroblasts and other CT of the limb.

Given the major developmental specification roles of these TFs, we hypothesize that their prominent constitutive expression represents some form of fore-versus-hindlimb positional memory. The functional relevance of these genes in maintaining and restoring regenerative identity in both species will be of interest to interrogate in the future.

Many *Hox* genes showed PD-biased expression in both organisms, often paralleling classic *Hox* gene expression patterns observed during limb development and consistent with dermal expression of Hox genes in human skin^29–31^. This Hox gene expression was largely specific to CT and widespread within CT subtypes. Non-*Hox* TF-encoding genes with important roles in PD developmental limb patterning – *Meis1*, *Meis2*, *Tbx15*, *Sp8* – also displayed PD-restricted expression patterns in both axolotl and mouse limbs.

Whereas genes encoding the best-characterized PD developmental signaling factors – BMP, FGF, WNT, RA – largely did not show biased expression that mirrored development, several genes that encode regulators of these pathways – *Grem1, Sfrp2,* and *Chrdl1* – showed similar proximal expression in both vertebrate limbs, with expression in CT.

Taking a broader look at genes not known to be involved in PD developmental limb patterning, we found several other genes with constitutive PD-biased, CT-enriched expression involved in processes that might impact positional identity. Whereas some of these expression patterns were shared between species – e.g., *Tgfbr2* and *Hpse2* – many more were species-specific. Among PD-biased gene expression, featured was a previously uncharacterized gene in axolotl that we named *Proxima*. *Proxima* is expressed broadly in proximal and medial axolotl limb CT, and encodes a predicted novel secreted protein found broadly in vertebrates, but lost in mammals.

Regarding the DV axis, *Lmx1b*, a key dorsal identity regulator during limb development, showed prominent dorsal expression in axolotl CT but not in mouse, whereas *En1*, a key regulator of ventral identity, showed prominent expression in ventral epidermis in both organisms. Along the AP axis, axolotl CT displayed clear posterior enrichment in the expression of the critical developmental posterior-specifying TF *Hand2*, which has an established posterior role in axolotl regeneration, and anterior enrichment in the expression of the anterior-specifying TF *Alx4*, along with other axial enrichment of expression for developmental TFs previously shown to have AP bias in axolotl limb tissue (*Hoxd13*, *Alx1*, *Lhx2*, and *Lhx9*)^16^. By contrast, the mouse limb AP axis displayed little to none of these key AP axial biases. In summary, much of the positionally biased gene expression in adult limbs appears similar between mouse and axolotl limb tissue; certain features, especially for the AP and DV axes are lacking in mouse (Fig. 10, Supplementary Data 15). Because the axolotl genome is large and recently sequenced, assembly and annotations are still being refined (e.g., *HoxC* gene models), which merits continued inquiry. This work used the AmbMex60DD genome assembly; upon mapping to another recent genome assembly (UKY_AmexF1_1, see Methods), the top gene expression pattern results, displayed in the model figure (Fig. 10), all remained significant (Supplementary Data 15).

**Figure 10.**
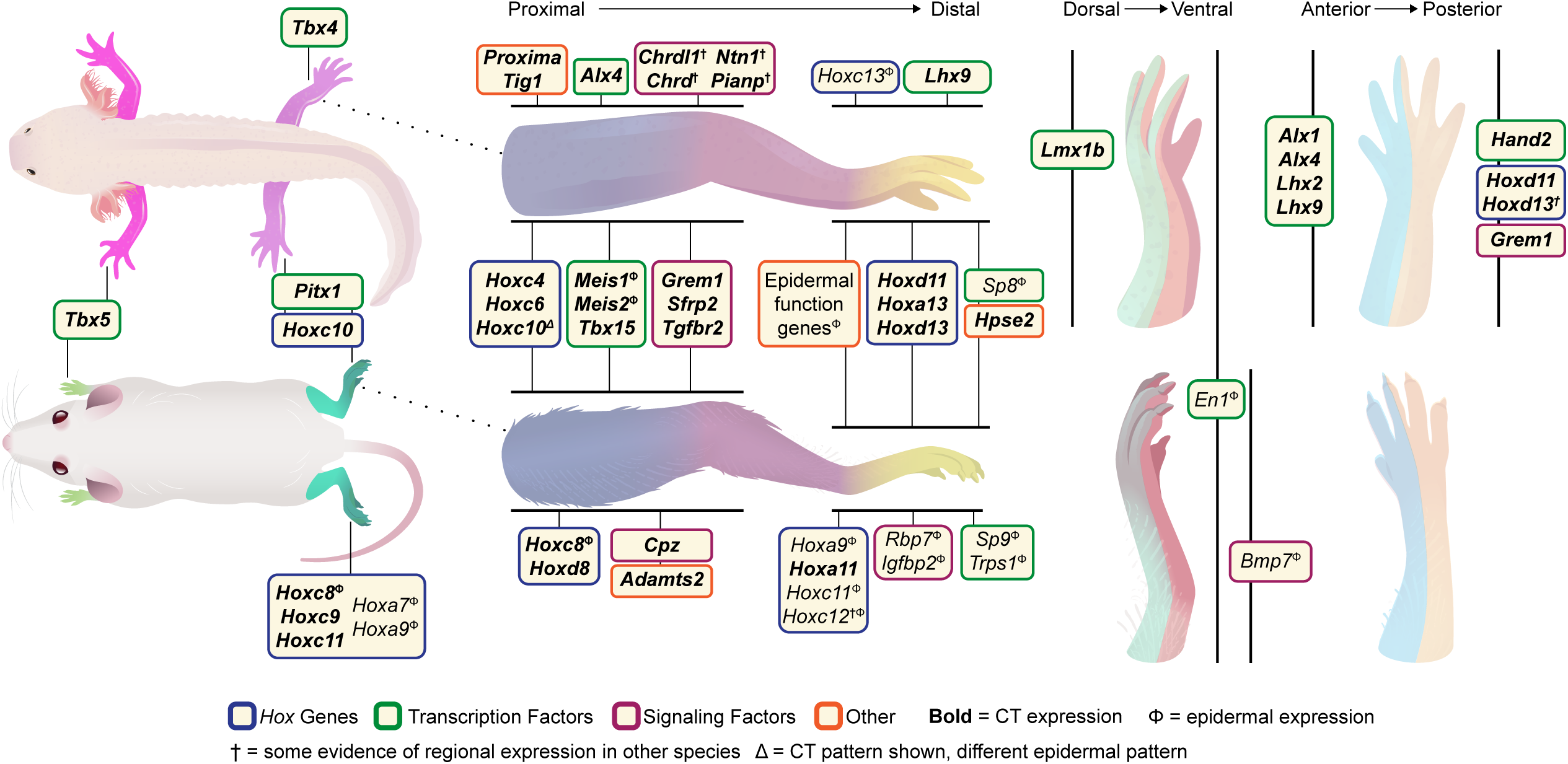
Model figure of vertebrate limb positional gene expression. Schematic representation of a model of positional gene expression across the vertebrate limb. All discussed axes represented. Genes are classified by functional category and tissue of expression enrichment (see key). This positional map model can continue to be refined going forward with future detailed inquiries into the function and properties of each gene presented. Genes were included if (1) there exists strong evidence of an expression pattern and the gene has been previously characterized as important for developmental limb patterning, or if (2) gene is involved in a process that could influence positional information and expression pattern has very strong evidence. For a more detailed summary of the evidence regarding each gene expression pattern shown, see Supplementary Data 15. Note that, whereas *Tbx4* expression was not overtly different in uninjured mouse forelimbs and hindlimbs, its expression appeared hindlimb-specific in regenerating digit tips (see Figure 9).

We additionally used MERFISH to explore positional gene expression during regeneration. Many of the discussed expression patterns were detectable within the first two weeks of axolotl limb regeneration, with AP-biased patterns typically being reestablished in a visibly obvious way earlier than PD-biased patterns. Looking at positional gene expression in limbs in the later stages of regeneration that were fully patterned but not yet fully rescaled (58 and 68 dpa) enabled the simultaneous visualization of striking forelimb-versus-hindlimb, PD, AP, and DV-biased gene expression patterns in the full limb for the first time, fully reestablished after tissue loss. Examination of gene expression in mouse digit tips with MERFISH as well as in pre-existing blastema scRNA-seq datasets demonstrated that forelimb- and hindlimb-specific gene expression differences – both in CT and epidermal populations – were also reestablished during regeneration in mouse, as well as the expression of genes shown to be specifically expressed in distal mouse limb tissue. This ability of newly regenerating tissue to also express positionally restricted genes in the same way as uninjured tissue demonstrates a form of molecular positional memory that is reestablished in newly formed cells after tissue loss in both organisms.

This work also contributed to a broader understanding of PD-biased gene expression in the epidermis in the vertebrate limb. Distal skin in both organisms displayed similar expression specificity, potentially associated with unique structural and immune roles for distal limb skin in its interface with the environment. We generated an online resource that generates expression data for any gene in our scRNA-seq atlas for each organism and provides selected interactive MERFISH data (digilimb.wi.mit.edu).

A majority of the limb development-related genes with adult positional expression bias were those encoding TFs, consistent with the possibility that CT might predominantly harbor positional memory in vertebrates cell-autonomously through TF expression.

These TFs could then be utilized to activate signaling factors in a regional manner during regeneration for patterning in regeneration. For example, posterior limb fibroblasts in axolotl express *Hand2,* but only begin expressing the signaling factor *Shh* in the regenerating limb^16^. This diverges from the case in planarians and acoels, in which many genes proposed to constitute adult positional information encode signaling factors that can influence surrounding cells, potentially related to the extensive cell turnover observed in these systems.

Including *Hand2*, only a handful of genes with constitutive regional expression have been well characterized as functionally influencing positional identity in axolotls. Future work characterizing the functional roles of the many gene expression patterns identified in this work will be essential for better understanding how regional expression patterns in uninjured tissue influence patterning outcomes in regeneration. In mouse, although the gene expression patterns identified cannot be present for full limb regeneration, these patterns could have other roles, such as in maintaining regional attributes of tissues and cells (e.g., epidermal characteristics), influencing local repair processes, in patterning during digit tip regeneration, or could not have an adult role but simply remain actively maintained after development. In at least one case, regional gene signatures of fibroblasts were shown to influence regional epidermal marker expression^31^. Better understanding of the function of regional gene expression in vertebrate tissues can be facilitated by future work detailing the role of TFs and secreted factors described here in adult tissue maintenance, repair, and regeneration. To better understand whether and how changes in positional gene expression are associated with regenerative ability in evolution, wider phylogenetic sampling, including within clades displaying variable regenerative capacities across species, will be important.

The observation that both acoels and planarians, bilatieran clades separated by hundreds of millions of years of evolution, display a robust system of adult positional information suggests that adult positional information maintenance in uninjured tissues may be an ancestral characteristic of the Bilateria, and therefore potentially present in many extant bilaterian clades. In hydra, a cnidarian, constitutive expression of Wnt pathway genes is associated with primary axis regeneration, consistent with the possibility that adult tissue positional memory pre-dates the Bilateria^12^. The scope of shared expression patterns mirroring developmental patterns in both the regenerative and non-regenerative adult vertebrate limb is consistent with the possibility that the last common ancestor of tetrapods had adult positional information across limbs and that similarities described here are explained by homology. Key differences between the observed mouse and axolotl CT expression patterns, particularly along the AP and DV axes, could be the result of loss in mammals or gain in amphibians. Broader species sampling will be of interest to address these possibilities.

In both regenerative and developmental contexts, CT has been tied to positional information maintenance. Among CT cell types, fibroblast heterogeneity characterization remains nascent. This work demonstrates that even fibroblasts within the same tissues display differences in the expression profiles of genes encoding ECM components and TFs, suggesting a complex diversity. Whereas many fibroblast subtypes in axolotl did not have clear counterparts in mouse and vice versa, a few subtypes did have clear similarity. Fibroblast subtypes might have diverged significantly between these two clades and some core functional subtypes likely share evolutionary ancestry and retain similarity.

Within the positional atlas of uninjured limbs obtained in this work, most genes involved in patterning during limb development that showed adult positional bias – e.g., limb *Hox* genes, *Tbx4, Tbx5, Tbx15, Pitx1, Sfrp2, Grem1, Meis1, Meis2, Lmx1b, Hand2, Alx4*, *Alx1*, *Lhx2*, *Lhx9* – displayed strong and enriched expression in CT (Fig. 10). In other words, whereas some of these genes have some expression in other tissues, most regional adult expression of development genes is restricted to CT. Furthermore, many of the regionally expressed genes with a strong CT-bias showed similar expression patterns between mouse and axolotl. Many of the genes expressed regionally across limb axes in CT were expressed throughout distinct CT types, although expression was often stronger and more widespread within axolotl CT versus mouse CT. In planarians, muscle harbors adult positional information and serves as the CT of the animals. These data are consistent with the hypothesis that diverse CT cell types maintain positional memory in adult tissues broadly across bilaterians.

One approach to improving regenerative outcomes is to understand mechanisms underlying the phenomenon in robustly regenerative species and to determine whether these mechanisms are present or absent in non-regenerative species. This work suggests that the broad landscape of positionally biased gene expression in the mammalian limb is partially similar to that in the regenerative axolotl limb, particularly along the proximal-distal limb axis. Some of these components could serve as a positional memory system to guide induced regenerative outcomes in the future, providing an endogenous blueprint for tissue identity. On the other hand, the reduced or missing elements of positional expression in mice could point to attributes contributing to a restriction of mammalian regenerative capacity.

**##Supplementary Figure 1. Quality control metrics for spatial limb atlases and *Tig1* expression in axolotl limb.**

(A) Expression of canonical cell type markers within the scRNA-seq spatial limb atlases, informing which clusters represent which cell types in the axolotl (top) and mouse (bottom) limb atlases. Expression of markers for cell types prominently present in only one organism atlas was still displayed for both organisms. (B) Distribution of genes per cell in each sample contributing to the axolotl (top) and mouse (bottom) limb atlases. (C) Distribution of UMIs per cell in each sample contributing to the axolotl (top) and mouse (bottom) limb atlases. (D) Expression of *Tig1* in bulk RNA-seq axolotl hindlimb skin dataset. ***=padj<0.001

**Supplementary Figure 2. Subclustering of non-fibroblast CT and expression of fibroblast subtype markers between species.**

(A) UMAP of all CT (center) and subclustered CT subsets (left and right) from axolotl limb, colored by subtype. Expression of genes encoding canonical CT subset markers (far left), core matrisome components (center left), matrisome-associated factors (center right), and TFs (far right) in these CT subsets (bottom). (B) UMAP of all CT (center) and subclustered specialized fibroblasts (left) and other CT subsets (right) from mouse limb, colored by subtype. Expression of genes encoding canonical CT subset markers (far left), core matrisome components (center left), matrisome-associated factors (center right), and TFs (far right) in these CT subsets (bottom). (C) Expression of mouse fibroblast subtype markers within axolotl fibroblast subtypes (top) and axolotl fibroblast subtype markers within mouse fibroblast subtypes (bottom). The fibroblast subtype marked by a set of genes is labelled at the top of each gene set. Expression of marker genes from fibroblast subtypes that have strong signatures of shared gene expression are highlighted in blue (axolotl *Col28a1*+ fibroblasts and *Reln*+ fibroblasts) and orange (axolotl *Rspo1*+ fibroblasts and *Kazald1*+ fibroblasts). *Rspo1+* axolotl fibroblasts did not express the best BLAST match for mouse *Kazald1*, but expressed a *Kazald1* paralog (see digilimb.wi.mit.edu). The *Col28a1* gene shown is best BLAST hit in the axolotl genome to mouse Col28a1 (see Supplementary Data 16), however, another *Col28a1* paralog is more highly expressed in the axolotl *Col28a1*+ fibroblasts (see Fig. 2C). (D) Expression of synovial fibroblast markers in axolotl (top) and mouse (bottom) limb fibroblast subtypes. Grey highlight emphasizes subtypes that most express these markers (axolotl *Comp*+ fibroblasts and mouse *Dlx3+* fibroblasts). ^†^=best mouse match.

**##Supplementary Fig. 3. Expression of forelimb and hindlimb developmental factors within mouse and axolotl CT.**

- MERFISH visualization of *Tbx5* expression within different CT subsets in the distal axolotl (left) and mouse (right) forelimb. Scale bars, 10 µm. Axolotl CT subsets were marked as follows: *Rspo1*+ fibroblasts - *Rspo1*, *Wnt7b*+ fibroblasts - *Bcl11b*, *Mmp19*+ fibroblasts - *Mmp19*, *Col6a5*+ fibroblasts - *Col6a5*, *Col28a1*+ fibroblasts - *Col28a1*, *Irf1*+ fibroblasts - *Mgp*, Chondrocytes - *Cilp2*, Tenocytes - *Tnmnd*, Osteoblasts - *Panx3*. Mouse CT subsets were marked as follows: *Kazald1*+ fibroblasts - *Cyp2f2*, *Meox2*+ fibroblasts - *Meox2*, *Hmcn2*+ fibroblasts - *Hmcn2*, *Dlx3*+ fibroblasts - *Htra4*, *Reln*+ fibroblasts - *Reln*, dermal papillary fibroblasts - *Hhip*, Chondrocytes - *Col2a1*,Tenocytes - *Kera*, Osteoblasts - *Dmp1*. (B) Visium HD visualization of *Tbx4* and *Tbx5* expression in mouse forelimb and hindlimb dermis. (C) Expression of *Hoxc* genes within different cell types present in scRNA-seq atlases (top) and CT subtypes (bottom) for axolotl (left) and mouse (right). Only medial tissue from hindlimb is represented. (D) MERFISH visualization of *Hoxc10* expression in axolotl (left) and mouse (right) forelimb and hindlimb CT. (E) Visium HD visualization of genes with forelimb or hindlimb epidermis expression bias, shown within Visium HD images (top) and computationally processed Visium HD UMAPs (bottom). (F) MERFISH visualization of *Hoxa7* and *Hoxa9* transcripts within forelimb and hindlimb epidermis. Epidermis is marked by *Lgals7.* CT was visualized via expression of *Pdgfra*, *Dpt*, and *Col6a1* in both organisms. This pool additionally included *Col4a1* in mouse. Scale bars are 100 µm.

**##Supplementary Fig. 4. Clustering and characterization of mouse forelimb and hindlimb 10X Chromium GEM-X scRNA-seq data.**

(A) UMAP representation of 10X Chromium GEM-X scRNA-seq datasets of mouse limb tissue, colored by cell type (left) and limb of origin (right). (B) Expression of canonical cell type markers within these limb datasets, informing which clusters represent which cell types. (C) Distribution of genes (left) and UMIs (right) per cell in each sample contributing to the 10X Chromium GEM-X scRNA-seq datasets. (D) UMAP of subclustered CT cells in 10X Chromium GEM-X scRNA-seq mouse limb datasets colored by subtype (left) and position of origin (right). (E) Expression of CT subtype markers within CT clusters in 10X Chromium GEM-X scRNA-seq mouse limb datasets.

**##Supplementary Fig. 5. *Hox* gene expression in hindlimb datasets.**

(A) MERFISH visualization of *Hoxa13* expression within different CT subsets in the distal axolotl (left) and mouse (right) limb. Scale bars, 10 µm. Axolotl CT subsets were marked as follows: *Rspo1*+ fibroblasts - *Rspo1*, *Wnt7b*+ fibroblasts - *Bcl11b*, *Mmp19*+ fibroblasts - *Mmp19*, *Col6a5*+ fibroblasts - *Col6a5*, *Col28a1*+ fibroblasts - *Col28a1*, *Irf1*+ fibroblasts - *Mgp*, Chondrocytes - *Cilp2*, Tenocytes - *Tnmnd*, Osteoblasts - *Panx3*. Mouse CT subsets were marked as follows: *Kazald1*+ fibroblasts - *Cyp2f2*, *Meox2*+ fibroblasts - *Meox2*, *Hmcn2*+ fibroblasts - *Hmcn2*, *Dlx3*+ fibroblasts - *Htra4*, *Reln*+ fibroblasts - *Reln*, dermal papillary fibroblasts - *Hhip*, Chondrocytes - *Col2a1*,Tenocytes - *Kera*, Osteoblasts - *Dmp1*. (B) MERFISH visualization *Hox* genes shown in Fig. 4 within medial limb tissue of axolotl (top) and mouse (middle and bottom). CT was visualized via expression of *Pdgfra*, *Dpt*, and *Col6a1* in both organisms. This pool additionally included *Col4a1* in mouse. Scale bars, 100 µm. (C) Expression of PD-biased non-canonical limb development *Hox* genes within different cell types present in scRNA-seq atlases (top) and CT subtypes (bottom) for axolotl (left) and mouse (right).

**##Supplementary Fig. 6. Epidermal subclustering and *Hox* gene expression.**

(A-B) UMAP of subclustered epidermal cell types colored by subtype (top) and positional origin (middle) within the axolotl (A) and mouse (B) scRNA-seq limb datasets. Expression of canonical marker genes used to confirm identity of epidermal subsets (bottom). (C) Expression of *Hox* genes that showed PD-biased expression within isolated epidermis of axolotl (top) and mouse (bottom) hindlimb. Grey backdrop indicates a statistically significant difference in expression in epidermis from at least two positions along the PD axis (p_adj_<0.05). (D) Expression of *Hox* genes shown in (C) within epidermal subsets of axolotl limb (left) and mouse limb (right). (E) Expression of skin-expressed *Hox* genes in proximal, medial, and distal limb skin of axolotl (left) and mouse (right) visualized by MERFISH or Visium HD (far right *Hoxa9* image). e=epidermis, d=dermis. Scale bars, 100 µm.

**##Supplementary Fig. 7. Further characterization developmental patterning factor gene expression.**

(A) Expression of genes from Fig. 5A within different cell types present in scRNA-seq atlases (top) and within CT (bottom) for the axolotl (left) and mouse (right) limb. (B) Expression of genes from Fig. 5A along the PD axis in axolotl (top) and mouse (bottom) hindlimb CT. (C) Expression of genes from Fig. 5B within CT subsets for the axolotl (left) and mouse (right) hindlimb. (D) Expression of genes from 5D within different cell types present in scRNA-seq atlases (left) and within CT subsets (right) for the axolotl (top) and mouse (mouse) limb. (E) Expression of genes from 5D in the bulk skin RNA-seq PD datasets of axolotl (left) and mouse (right) hindlimb skin. (F) Expression of developmental signaling factors that display PD-biased expression in isolated epidermis of axolotl (left) and mouse (right) hindlimb. (G) Expression of genes from (F) within epidermal subsets of axolotl limb (left) and mouse limb (right). Grey backdrop indicates a statistically significant difference in expression in epidermis from at least two positions along the PD axis (p_adj_<0.05). *=p_adj_<0.05, **=p_adj_<0.01, ***=p_adj_<0.001.

**##Supplementary Fig. 8. Further characterization of expression of PD-biased gene expression in vertebrate limbs.**

(A) Plot from Fig. 6A with previously discussed developmental limb patterning factors labelled. The rank of these factors was used to set thresholds of interest in Fig. 6A. (B) Expression genes from 6A within different cell types present in scRNA-seq atlases (left) and within CT subsets (right) for the axolotl (top) and mouse (mouse) limb. (C) MERFISH visualization of genes shown in Fig. 6D in axolotl (top) and mouse (bottom) medial hindlimb tissue. CT marked by *Pdgfra*, *Dpt*, and *Col6a1* in axolotl. CT in mouse Visium HD marked by *Pdgfra*, *Col3a1*, *Dpt*, *Col4a1*, *Dcn*, *Col15a1*, *Pi16*, *Tagln*, and *Acta2*. (D) Visium HD visualization of more genes from Fig. 6A that show differential PD expression in mouse limb tissue. Scale bars, 100 µm. ^†^=best axolotl BLAST hit.

**##Supplementary Fig. 9. Further characterization of PD-biased unannotated axolotl genes.**

(A) Heatmap of bulk skin RNA-seq expression of unannotated axolotl genes from Fig. 6E along the PD axis in the axolotl hindlimb. (B) Expression of unannotated axolotl genes from Fig. 6E in proximal, medial, and distal hindlimb CT in axolotl. (C) UMAP representation of expression of unannotated axolotl genes from Fig. 6E in axolotl CT. (D) MERFISH visualization of *Proxima* expression within different CT subsets in the proximal axolotl limb tissue. Scale bars, 10 µm. Axolotl CT subsets were marked as follows: *Rspo1*+ fibroblasts - *Rspo1* and *Grem2*, *Wnt7b*+ fibroblasts - *Bcl11b*, *Mmp19*+ fibroblasts - *Mmp19*, *Col6a5*+ fibroblasts - *Col6a5*, *Col28a1*+ fibroblasts - *Col28a1*, *Irf1*+ fibroblasts - *Mgp*, Chondrocytes - *Col2a1*, Tenocytes - *Comp* and *Tnc*. (E) Predicted protein structure generated using ColabFold of unannotated axolotl genes from Fig. 6E (AMEX60DD021955 excluded because predicted to be non-coding) with some basic domain information. (F) Metazoan phylogeny showing the presence or absence of *Proxima* orthologs. For each clade, species with best BLAST hit to axolotl *Proxima* is shown, along with the corresponding E-value. Clade needed to have an ortholog with E-value < 1e-20 to be marked as having *Proxima*. All orthologs indicated in phylogeny are best mutual BLAST hits with *Proxima*. (G) Alignments of *Proxima* and orthologs labelled in (F). ***=p_adj_<0.001

**##Supplementary Fig. 10. Further characterization of epidermally-expressed genes.**

(A) Functional classification of top 200 differentially expressed genes along the PD axis in the bulk RNA-seq skin datasets of axolotl (left) and mouse (right), separated by whether gene shows proximal/medial enrichment (top) or distal enrichment (bottom). Genes were sorted by p_adj_ and for genes that appeared in multiple comparisons (proximal v. distal, proximal v. medial, medial v. distal), only data from the lowest p-value result was used. (B) MERFISH visualization of genes from Fig. 6K in medial axolotl (top) and mouse (middle) skin. Visium HD visualization of *Lce* and *Sprr* gene expression in mouse medial skin (bottom). *Lce1a1*, *Lce1d*, *Lce1e, Lce1f, Lce1k,* and *Lce1l* transcripts are shown. Epidermis is marked by *Lgals7* in mouse and *Scel* in axolotl. (C) Expression of select genes that show strong PD-bias within isolated mouse epidermis (top) and expression of these genes within mouse epidermal subtypes (middle). Visium HD visualization of these genes in proximal, medial, and distal mouse skin (bottom). (E-F) Expression of select genes from Fig. 6J that show significant PD- bias in isolated epidermis scRNA-seq data. Expression shown in proximal, medial, and distal skin (top) as well as within different epidermal subsets (bottom) for axolotl (D) and mouse (E) hindlimb skin. (F) Visium HD visualization of transcripts for genes with PD-biased epidermal expression, shown within Visium HD images (top) and computationally processed Visium HD UMAPs (bottom). Grey backdrop indicates a statistically significant difference in expression in CT from at least two positions along the PD axis (p_adj_<0.05). e=epidermis, d=dermis. Scale bars are 100 µm.

**##Supplementary Fig. 11. Characterization of a regional axolotl skin domain.**

(A) MERFISH visualization of genes with distally-biased expression within sequencing data that localize to a particular domain of the anterior, ventral, and distal axolotl epidermis in both forelimb and hindlimb. Scale bars are 1 mm. (B) Expression of genes in (A) within axolotl epidermal scRNA-seq data. (C) Expression of genes in (A) and literature markers of the nuptial pad in *Rana chensinensis*^152^, identified below the plot, within subclustered axolotl epidermis, including a nuptial pad cluster identified by expression of homologs of nuptial pad markers.

**##Supplementary Fig. 12. Further characterization of DV-biased gene expression in vertebrate limbs.**

(A) Heatmap bulk RNA-seq expression of genes with DV expression bias in mouse tissue that encode signaling factors, TFs, or ECM regulators. *En1* is highlighted with an arrowhead. (B) Visium HD visualization of *En1* and *Bmp7* expression in mouse dorsal and ventral skin. Scale bars, 100 µm. (C) Expression of genes from Fig. 7F within different cell types present in scRNA-seq atlases (left) and within CT subsets (right) for the axolotl (top) and mouse (mouse) limb. *=p_adj_<0.05, **=p_adj_<0.01, ***=p_adj_<0.001. The axolotl *Hand2\** locus refers to a distinct gene in the axolotl genome assembly from the canonical *Hand2* referenced in other figures (see Supplementary Data 16).

**##Supplementary Fig. 13. Further characterization of AP-biased gene expression in vertebrate limbs.**

(A) MERFISH visualization of *Lhx2, Hoxd13,* and *Grem1* transcripts in anterior and posterior distal hindlimb tissue. *Lhx2* is not significantly enriched in anterior CT by quantification (see Methods, Supplementary Data 17). It is displayed because this expression enrichment was published^16^ and also clearly visualized in regenerated tissue in Fig. 9. Scale bars are 500 µm. (B) Expression of genes from Fig. 8E within different cell types present in scRNA-seq atlases (left) and within CT subsets (right) for the axolotl (top) and mouse (mouse) limb. *=p_adj_<0.05, **=p_adj_<0.01, ***=p_adj_<0.001. ^†^=best axolotl BLAST hit.

**##Supplementary Fig. 14. Further characterization of positionally restricted gene expression during and after regeneration.**

(A) MERFISH visualization of forelimb and hindlimb-specific gene expression in regenerating and regenerated axolotl limbs. 14 dpa CT image is the same as shown in Fig. 9D. (B) MERFISH visualization of transcripts for genes with PD-biased expression in uninjured tissue in 58 dpa regenerated axolotl hindlimb. (C) MERFISH visualization of genes with AP-biased expression in 58 dpa regenerated axolotl hindlimb.(D) MERFISH visualization of transcripts for genes that had positionally restricted epidermal expression in uninjured mouse tissue in 14 dpa regenerating forelimb and hindlimb mouse digit tips. CT marked by *Pdgfra*, *Dpt*, and *Col6a1* in axolotl. Epidermis marked by *Lgals7* and *Calm4* in mouse. Images of 8 dpa and 11 dpa axolotl limbs are from juvenile regenerating axolotls. Scale bars, 100 µm for 8 dpa and 11 dpa axolotl images and 500 µm for 58 dpa and 68 dpa axolotl. Scale bars, 100 µm for mouse 14 dpa images.

**##Supplementary Fig. 15. Further characterization of genes with PD-biased expression within mouse digit tip regeneration datasets.**

(A) UMAP of all cells (top) and subsclustered CT cells (bottom) present in digit tip regeneration datasets, colored by cell type (top) and regeneration timepoint (bottom). (B) Expression of developmental patterning-related genes that displayed PD-biased expression in uninjured mouse CT within different cell types (top) and CT from different regenerative time points (bottom). (C) Expression of genes that displayed PD-biased expression in uninjured mouse epidermis within different cell types (top) and epithelial cells from different regenerative time points (bottom). (D) UMAP of all cells (top) and subsclustered CT cells (bottom) present in digit tip regeneration datasets, colored by cell type (top) and regeneration timepoint (bottom). (E) Expression of *Hox* genes that displayed PD-biased expression in uninjured mouse CT within different cell types (top) and CT from different regenerative time points (bottom). (F) Expression of developmental patterning-related genes that displayed PD-biased expression in uninjured mouse CT within different cell types (top) and CT from different regenerative time points (bottom). (H) Expression of genes that displayed PD-biased expression in uninjured mouse epidermis within different cell types (top) and epithelial cells from different regenerative time points (bottom). UA = unamputated. (A-C) display data from Storer et al., 2020^33^ and (D-H) display data from Johnson et al., 2020^32^.

**##Supplementary Data 1. Single cell atlas markers for vertebrate limbs.**

Results of Seurat FindAllMarkers (binom) shown for axolotl and mouse. Gene name, p-value, adjusted p-value, average log_2_FC, cluster name, percent 1 (percent of cells within cluster that express labelled gene), and percent 2 (percent of cells within other clusters that express labelled gene) shown.

**##Supplementary Data 2. CT markers for vertebrate limbs.**

Results of FindAllMarkers (binom) for CT in both axolotl and mouse limbs. Full list, as well as list filtered for ECM components and TFs shown for each organism. Gene name, p-value, adjusted p-value, average log_2_FC, cluster name, percent 1 (percent of cells within cluster that express labelled gene), and percent 2 (percent of cells within other clusters that express labelled gene) shown.

**##Supplementary Data 3. Differentially expressed genes between vertebrate forelimb and hindlimb CT from single-cell RNA-seq datasets.**

Results of FindMarkers (binom) for CT from forelimb and hindlimb of axolotl and mouse. Gene name, p-value, adjusted p-value, average log_2_FC, percent 1 (percent of cells within cluster that express labelled gene), and percent 2 (percent of cells within other clusters that express labelled gene) shown.

**##Supplementary Data 4. Single-cell atlas markers for mouse forelimb and hindlimb medial sections.**

Results of FindAllMarkers (binom) for full atlas and additionally for CT subsets within the GEM-X datasets of mouse forelimb and hindlimb medial tissue. These were clustered separately due to lack of easy integration with other datasets, likely because of differences in kit chemistry. Gene name, p-value, adjusted p-value, average log_2_FC, percent 1 (percent of cells within cluster that express labelled gene), and percent 2 (percent of cells within other clusters that express labelled gene) shown.

**##Supplementary Data 5. Differentially expressed genes between forelimb and hindlimb and along the PD axis in mouse Visium HD data.**

Results of FindMarkers (binom) for CT and keratinocytes (Kcytes) from forelimb medial and hindlimb proximal, medial, and distal computationally processed Visium HD bins (see Methods). For PD axis data the following is shown: raw results of comparisons between any two samples, the same data with duplicates removed (only highest p_adj_ result is kept), and then further filtered for both CT and keratinocytes. For forelimb- versus-hindlimb data, raw results are shown. Gene name, p-value, adjusted p-value, average log_2_FC, percent 1 (percent of cells within cluster that express labelled gene), and percent 2 (percent of cells within other clusters that express labelled gene) are shown. For PD comparisons, the number of pairwise comparisons that showed a significant difference as well as the pairwise comparison with the lowest p_adj_ are noted.

**##Supplementary Data 6. Differentially expressed genes along the PD axis in skin bulk RNA-seq data for mouse and axolotl limb.**

DESeq2 results constituting differentially expressed genes between proximal, medial, and distal skin bulk RNA-seq samples in mouse and axolotl limb. For each organism, the following data is presented: raw DESeq2 results of comparisons between any two samples, the same data with duplicates removed (only highest p_adj_ result is kept), the top 200 overall hits annotated with functional and localization information, and a list of the genes categorized as potentially influencing positional information based on broad functional characterization. Gene name, gene ID, p-value, adjusted p-value, comparison of origin, base mean, Wald stat, log_2_FC,and log_2_FC standard error is included for each gene comparison. Also shown is the number of pairwise comparisons that showed a significant difference as well as the pairwise comparison with the lowest p_adj_ value.

**##Supplementary Data 7. Differentially expressed genes along the PD axis in vertebrate limb CT from single-cell RNA-seq datasets.**

Results of FindMarkers (binom) for CT from proximal, medial, and distal hindlimb of axolotl and mouse. For each organism the following is shown: raw results of comparisons between any two samples, the same data with duplicates removed (only highest p_adj_ result is kept), duplicate removed data filtered for just ECM components and TF encoding genes, the top overall hits annotated with functional and localization information, and a list of the genes categorized as potentially influencing positional information based on broad functional characterization. Gene name, p-value, adjusted p-value, average log_2_FC, percent 1 (percent of cells within cluster that express labelled gene), and percent 2 (percent of cells within other clusters that express labelled gene) shown. Also shown is the number of pairwise comparisons that showed a significant difference as well as the pairwise comparison with the lowest p_adj_ value.

**##Supplementary Data 8. Epidermal markers and PD-biased gene expression in vertebrate limb epidermis.**

Results of FindMarkers (binom) for epidermis subclustering and comparisons of epidermis for proximal, medial, and distal limb in mouse and axolotl. For each organism, the following is shown: FindAllMarkers results for epidermis subclustering providing markers for each subset, results showing the top differentially expressed genes between PD comparisons of skin, and these same results filtered for ECM components and TFs only. Gene name, p-value, adjusted p-value, average log_2_FC, cluster name, percent 1 (percent of cells within cluster that express labelled gene), and percent 2 (percent of cells within other clusters that express labelled gene) shown. Also shown is the number of pairwise comparisons that showed a significant difference as well as the pairwise comparison with the lowest p_adj_ value.

**##Supplementary Data 9. Positionally restricted gene expression in other axolotl cell types.**

Results of FindMarkers (binom) for endothelium (Endo), satellite cells (SatCells), smooth muscle (SMusc), and Schwann cells across the forelimb-hindlimb, PD, AP, and DV axes in axolotl limb tissue. PD results include raw results and results filtered for duplicates from pairwise comparisons. Gene name, p-value, adjusted p-value, average log_2_FC, percent 1 (percent of cells within cluster that express labelled gene), and percent 2 (percent of cells within other clusters that express labelled gene) are shown. For PD comparisons, the number of pairwise comparisons that showed a significant difference as well as the pairwise comparison with the lowest p_adj_ value are noted. Summary tab displays any positionally restricted expression of developmentally relevant genes in the appropriate axes within any of these cell types.

**##Supplementary Data 10. Positionally restricted gene expression in other mouse cell types.**

Results of FindMarkers (binom) for endothelium (Endo), satellite cells (SatCells), smooth muscle (SMusc), and Schwann cells across the forelimb-hindlimb, PD, AP, and DV axes in mouse limb tissue. PD results include raw results and results filtered for duplicates from pairwise comparisons. Gene name, p-value, adjusted p-value, average log_2_FC, percent 1 (percent of cells within cluster that express labelled gene), and percent 2 (percent of cells within other clusters that express labelled gene) are shown. For PD comparisons, the number of pairwise comparisons that showed a significant difference as well as the pairwise comparison with the lowest p_adj_ value are noted. Summary tab displays any positionally-restricted expression of developmentally-relevant genes in the appropriate axes within any of these cell types.

**##Supplementary Data 11. Characterization of top differentially expressed unannotated axolotl genes along the PD axis.**

Information about top differentially expressed unannotated axolotl genes across the PD axis including assay ranks, BLAST results, domain search results, and DeepLoc results. Detailed information regarding the *Proxima* ortholog phylogeny also included.

**##Supplementary Data 12. Differentially expressed genes along the DV axis in vertebrate limb CT from single-cell and bulk RNA-seq datasets**

Results of FindMarkers (binom) for CT from dorsal and ventral hindlimb of axolotl and mouse. For each organism the following is shown: raw results of comparisons positions, the top overall hits annotated with functional and localization information, and a list of the genes categorized as potentially influencing positional information based on broad functional characterization. Gene name, p-value, adjusted p-value, average log_2_FC, cluster name, percent 1 (percent of cells within cluster that express labelled gene), and percent 2 (percent of cells within other clusters that express labelled gene) shown. Results of DESeq2 differential gene expression analysis for the bulk RNA-seq dorsal and ventral mouse limb tissue datasets also shown. Gene name, gene ID, p-value, adjusted p-value, base mean, Wald stat, log_2_FC,and log_2_FC standard error is included for each gene comparison.

**##Supplementary Data 13. Differentially expressed genes along the AP axis in vertebrate limb CT from single-cell RNA-seq datasets.**

Results of FindMarkers (binom) for CT from anterior and posterior hindlimb of axolotl and mouse. For each organism the following is shown: raw results of comparisons between positions, the top overall hits annotated with functional and localization information, and a list of the genes categorized as potentially influencing positional information based on broad functional characterization. Gene name, p-value, adjusted p-value, average log_2_FC, percent 1 (percent of cells within cluster that express labelled gene), and percent 2 (percent of cells within other clusters that express labelled gene) shown.

**##Supplementary Data 14. Characterization of top differentially expressed unannotated axolotl genes along other major limb axes.**

Information about top differentially expressed unannotated axolotl genes across the AP, DV, and forelimb-hindlimb axes. This includes assay ranks, BLAST results, domain search results, and DeepLoc results.

**##Supplementary Data 15. Data summary for gene expression patterns shown in the model figure.**

Detailed evidence shown in this work for each expression pattern shown in Fig. 9. Key shown in table. Also includes analysis of the axolotl genes shown in the model figure when scRNA-seq data was mapped to another genome assembly.

**##Supplementary Data 16. Annotation of axolotl genes referenced in figures.** Shown are the genome IDs and annotations for each axolotl gene referenced in this work, organized by figure. Also included is a reciprocal BLAST analysis for mouse CT subpopulation markers queried in axolotl and information about axolotl genes excluded in this work because of annotation discrepancies.

**##Supplementary Data 17. Quantification of MERFISH expression patterns.**

Results of FindMarkers (binom) for computationally segmented and processed cells (see Methods) from MERFISH experiments from different positions along the forelimb-hindlimb, PD, DV, and AP axes in mouse and axolotl limb. Cell and gene filtering criteria as well as notes on genes excluded from analysis are included within the document. Gene name, adjusted p-value, average log_2_FC, percent 1 (percent of cells within cluster that express labelled gene), and percent 2 (percent of cells within other clusters that express labelled gene) are shown.

## Materials and Methods

### Mouse Sourcing and Tissue Acquisition

Mice were acquired by, cared for, and handled by the Preclinical Modeling Facility at the Koch Institute. CD-1 IGS adult male mice were used, all fully sexually developed and at least 3 months old. For single-cell sequencing experiments, animals were perfused with PBS with 5IU/mL heparin sulfate at 5mL/min for a total of 40mL in order to avoid overrepresentation of blood cells in single-cell RNA-sequencing datasets. All experiments conducted with right hindlimbs.

### Axolotl Sourcing and Tissue Acquisition

Axolotls (Ambystoma mexicanum) were bred in and shipped from the Ambystoma Genetic Stock Center located in the College of Medicine at the University of Kentucky. Male adult axolotls were used in all experiments, all fully sexually developed and at least 14 months old. Animals used were between 20 cm and 25 cm from head tip to tail tip. Leucistic male axolotls were used for all single-cell RNA-sequencing experiments, while wild type males were used for bulk RNA-sequencing experiments due to the higher replicate requirements and greater ease of wild type adult acquisition. Animals were anesthetized in a pH adjusted solution of 0.1-0.2% tricaine (MS-222) for 15 minutes and checked for responsiveness prior to amputations. Experiments typically required the harvesting of the full limb up to the intersection with the trunk and in these cases, axolotls were euthanized to avoid recovery from extreme amputation. Amputations were done in accordance with protocols described in Kragl and Tanaka, 2009^187^. All experiments conducted with right hindlimbs.

### Mouse Bulk RNA-Sequencing

For mouse bulk RNA-sequencing experiments, all tissue was harvested from recently euthanized mice. Hair was removed from the right hindlimb using a razor blade. Hindlimb was then surgically separated from the rest of the mouse using a scalpel. This was done by extending the limb, identifying the hip joint by touch, making a skin incision all the way around the hip joint to separate limb skin from trunk skin, and then making an incision separating the femur at the hip to separate bone, muscle, and internal fascia associated with the hindlimb. For the proximal-distal axis dataset, tissue was then segmented at the knee joint and ankle joint to form proximal, medial, and distal sections. Skin (this included epidermis, dermis, and hypodermis while excluding skeletal muscle, bone, and deeper connective tissue and vasculature) was peeled from each section and placed into Trizol (Life Technologies) into an Eppendorf Safe-Lock Tube with a metal bead. In the distal (paw) section, skin was difficult to remove entirely from the digits without contaminating with other tissues; therefore skin was harvested from the ventral and dorsal hindpaw, as well as the from the base of the digits to capture as much uncontaminated skin sample as possible. For the dorsal-ventral axis dataset, after the hindlimb was separated at the hip joint as described above, the midline between the dorsal and ventral limb surface was marked with a shallow scalpel incision using the dorsal-ventral orientation of the paw, in which the back of the paw marks the dorsal side and the sole marks the ventral side. The incision was then deepened to the bone and tissue from each side (including skin, skeletal muscle, and deeper fascia and vasculature), excluding bone, and placed into separate Eppendorf Safe-Lock Tube with a metal bead. The samples in the tubes were then disrupted using a TissueLyser II (Qiagen) to ensure RNA could be captured from the entire sample. Total RNA was extracted using standard Trizol RNA extraction protocol. RNA was extracted from each leg section in biological quintuplicate. Libraries were prepared using the KAPA mRNA HyperPrep Kit with the KAPA Dual-Indexed Adapter Kit (KapaBiosystems) and sequenced on an Illumina HiSeq 2000.

### Axolotl Bulk RNA-Sequencing

For axolotl, right hindlimbs were surgically removed from the axolotl after anesthetization at the location where the limb met the trunk surgical scissors. Incisions were made with a sterile scalpel at the knee and ankle joints to segment the limb into proximal, medial, and distal sections. Skin (this included epidermis, dermis, and hypodermis while excluding skeletal muscle, bone, and deeper connective tissue and vasculature) was peeled from each section and placed into Trizol (Life Technologies) into an Eppendorf Safe-Lock Tube with a metal bead. Skin was fairly cleanly separable from the digits in this context and so distal skin samples were harvested in their entirety. Processing and extraction of RNA proceeded from this point similarly to the process described above for mouse, except libraries were sequenced on an Illumina NovaSeq 6000 SP. RNA was extracted from each leg section in biological quintuplicate.

### Bulk RNA-Sequencing Analysis

Reads were mapped to the GRCm39 genome assembly for mice and axolotl v6.0-DD genome assembly for axolotl using CellRanger. Differential expression analysis was performed using DESeq2^188^.

### Mouse Single-Cell RNA Sequencing

Prior to single-cell RNA-sequencing experimentation, mice were perfused as described above to eliminate circulating blood cells and then euthanized. Mouse limbs were shaved with a razor blade and the right hindlimb was removed from the rest of the body with a sterile scalpel, as described in Mouse Bulk RNA-Sequencing. Proximal, medial, and distal sections were segmented as described in Mouse Bulk RNA-Sequencing. This was done with two replicate limbs for single-cell RNA-sequencing, each segmented into proximal, medial, and distal sections. For both replicates, the bone tissue was included in the dissociation and sequencing submission for the distal section. In the first replicate, bone was included in the dissociation and sequencing submission for the proximal and medial sections as well, but appeared to contribute to the final processed dataset being heavily biased toward inclusion of leukocytes. With this in mind, bone was excluded from the proximal and medial samples in the second replicate. All other tissue, including skin, skeletal muscle, as well as deeper fascia and vasculature, were all included in the sample preparation. For the dorsal and ventral datasets, the full limb was separated from the body and dorsal and ventral tissue was collected as described in Mouse Bulk RNA-Sequencing. For the anterior and posterior datasets, the hindlimb was first separated from the body as described in Mouse Bulk RNA-Sequencing. Using the clear anterior-posterior orientation of the paw, a shallow incision was made along the midline of the anterior-posterior axis along the length of the limb. The incision was then deepened to the bone and tissue on the anterior and posterior side (including skin, skeletal muscle, and deeper fascia and vasculature) was harvested separately. Bone was excluded except in the paw. The paw was cut so that the two anterior-most digits were included in the anterior dataset and the two poster-most digits were included in the posterior dataset. The middle digit was excluded. One cell preparation each of dorsal, ventral, anterior, and posterior tissue was isolated and processed for single-cell RNA-sequencing in this manner. For the forelimb-versus-hindlimb comparisons, only the medial section of each limb was processed for sequencing, as described for medial sections above. One cell preparation each of forelimb medial and hindlimb medial tissue was used for the generation of these datasets.

Dissociation began with the dicing of the mouse tissue into very small pieces with a sterile scalpel in a small amount of PBS to keep the tissue hydrated. Tissue was then transferred to a solution of 100μg/mL Liberase TM (Sigma) and nutated at 37C for 4 hours with hourly manual mixing of the tissue to ensure penetration of the enzymatic solution. The tissue solution was then filtered through a 100μm filter and centrifuged at 500xg for 5 minutes. The supernatant was removed and the pellet resuspended in 1% bovine serum albumin (BSA) (Sigma) in PBS. The solution was then centrifuged again at 500xg for 5 minutes. The supernatant was removed and the pellet was resuspended in 1% BSA in PBS. The cellular suspension was then filtered through a 40μm filter and stained 1:1000 with calcein AM (Invitrogen) and 1:200 propidium iodide (PI).

Suspension was then submitted to FACS, where cells that were calcein+ and PI- were captured in 0.01% BSA in PBS. Cells were then counted on a hemocytometer and resuspended in 0.01% BSA in PBS at a concentration of 1000 cells/μl for 10X single cell sequencing.

### Axolotl Single-Cell RNA Sequencing

After limb amputation, the axolotl limb was placed in 0.8X PBS to prevent drying out. For proximal, medial, and distal datasets, the limb was removed and segmented as described in Axolotl Bulk RNA-Sequencing. All tissue, including skin, bone, skeletal muscle, as well as deeper fascia and vasculature, were all included in the sample preparation. For the dorsal and ventral samples, the hindlimb was isolated as described in Axolotl Bulk RNA-Sequencing. Using a sterile scalpel, a shallow incision was made along the midline of the dorsal-ventral axis along the length of the limb, using the clear dorsal-ventral orientation of the foot for guidance. Proximal, medial, and distal sections were then segmented as described in Axolotl Bulk RNA-Sequencing. For ease of dissection and comparison, only the medial section was used. On the medial section, the incision was then deepened to the bone and tissue on the dorsal and ventral side (including skin, skeletal muscle, and deeper fascia and vasculature) was harvested separately. Bone was not included due to difficulty of dissection. For anterior and posterior samples, tissue was harvested in a similar way, with the incision following the midline of the anterior-posterior axis instead, oriented using the obvious anterior-posterior axis of the foot. For ease of dissection and comparison, only the medial section was used and bone was included for these samples. Three biological replicate limbs were processed to generate proximal, medial, and distal samples. Two biological replicate limbs were processed for anterior and posterior samples and another two for anterior and posterior samples. One limb was processed to generate a medial forelimb sample.

Dissociation began with dicing all the collected tissue from the desired section into very small pieces with a sterile scalpel in a small amount of 0.8X PBS. Tissue was then transferred to a solution of 50μg/mL Liberase TM (Sigma) and 0.5U/mL DNAse I (Worthington) in 0.8X PBS and nutated for 55 minutes at room temperature. The tissue was mechanically disturbed with a pipette once every ∼15 minutes during the nutation. The tissue solution was then filtered through a 100μm filter and spun down at 300xg for 5 minutes. The supernatant was removed and the pellet was resuspended in 1% BSA in 0.8X PBS. The cellular suspension was then filtered through a 30μm filter and spun down once more at 300xg for 5 minutes. The supernatant was removed and the pellet was resuspended in 0.01% BSA in 0.8X PBS and filtered through a 30μm filter. The solution was spun down at 300xg for 5 minutes, supernatant removed, and the pellet resuspended in 0.01% BSA in 0.8X PBS once more. This solution was then filtered through a 30μm filter. Cells were then counted on a hemocytometer and resuspended in 0.01% BSA in 0.8X PBS at a concentration of 1000 cells/μl for 10X single-cell sequencing.

### Single-Cell RNA Sequencing Analysis

After dissociation and resuspension, cells were processed either by either the Genome Technology Core at the Whitehead Institute or through internal lab processing using either 10X Genomics Chromium Controller, 10X Genomics Chromium iX Controller, or 10X Genomics Chromium X Controller. Following kits were used in accordance with manufacturer instructions and protocols: Chromium Next GEM Single Cell 3’ Kit v3.1t (PN-1000268), Chromium Next GEM Single Cell 3’ HT Reagent Kits v3.1 (PN- 1000348), and Chromium GEM-X Universal 3’ Gene Expression v4 (PN-1000691). All mouse samples and replicates were processed with Chromium Next GEM Single Cell 3’ Kits, except the forelimb medial and hindlimb medial datasets which were processed with Chromium GEM-X Universal kits. The three biological replicates each for the proximal, medial, and distal axolotl hindlimb datasets were processed as follows: two were processed using Chromium Next GEM Single Cell 3’ Kits and one was split into two technical replicates that were processed with Chromium Next GEM Single Cell 3’

HT Reagent Kits. The two biological replicates each for axolotl hindlimb anterior, posterior, dorsal, and ventral datasets were processed as follows: one was processed using Chromium Next GEM Single Cell 3’ Kits and one was split into two technical replicates that were processed with Chromium Next GEM Single Cell 3’ HT Reagent Kits. The one biological preparation for medial axolotl forelimb was split into two technical replicates that were processed with Chromium Next GEM Single Cell 3’ HT Reagent Kits. Recommended cell numbers were loaded to maximize the number of cells analyzed. Amplified cDNA libraries were submitted to bioanalyzer analysis for quality control and size selected using magnetic beads in accordance with manufacturer’s recommendations. Mouse samples were sequenced on an Illumina HiSeq 2000 and axolotl samples were sequenced on either an Illumina HiSeq 2000 or an Illumina NovaSeq 6000 S4. Sequencing reads were mapped to the GRCm39 genome assembly (NCBI: GCF_000001635.27) for mouse and AmbMex60DD genome assembly (GenBank: GCA_002915635.3) for axolotl using Cell Ranger. Reads that represented ambient RNA or hybridization artifacts were computationally modelled and removed using CellBender^189^. Cell doublets were removed using scDblFinder^190^. Cells were assessed for UMI and gene counts, which are represented by violin plots in Supplementary Fig. 1. For axolotl, cells with <400 or >75,000 UMIs were excluded, as were cells with <100 or >10,000 unique genes. For mouse, cells with <400 or >40,000 UMIs were excluded, as were cells with <100 or >4,000 unique genes. Further analysis was performed using Seurat^191^, which was used to cluster (Leiden) and visualize cells using uniform manifold approximation and projection (UMAP). Cluster identities were assigned using known markers of mouse and axolotl cell types, detailed in the figures above. Processed Seurat objects for these datasets are available on Zenodo: 10.5281/zenodo.18316185. Differential expression analysis was performed between groups of computationally isolated cells using the binomial test within the FindMarkers function in Seurat analysis for both datasets. For mouse, this analysis was done separately for forelimb-versus-hindlimb comparisons within CT in a separate dataset, as these did not integrate well into the rest of the atlas due to differences in GEM-X and Next GEM chemistry. When filtering for ECM components, annotated lists of components in mouse and human from The Matrisome Project^192^ were used. When filtering for TFs, the compiled list of mouse and human TFs from Ravasi et al., 2010^193^ was used. For fibroblast subtype marker analysis (see Supplementary Data 16, Supplementary Fig. 2C), the longest annotated protein isoform encoded by each mouse subtype marker was queried against the predicted axolotl proteome using BLASTP. The longest annotated protein isoform of each highest scoring hit was then queried to the mouse proteome to look for the best reciprocal BLASTP hit for each mouse subtype marker. We additionally mapped all axolotl scRNA-seq datasets to the UKY_AmexF1_1 genome assembly (NCBI: GCF_040938575.1) using Cell Ranger. To check that the axolotl genes presented in our model (Fig. 10) were expressed in a similar manner when mapped to this new assembly, we isolated the subset of this newly mapped dataset that corresponded to our CT and epidermal subclustered datasets. We then performed differential expression analysis as described above and assessed the expression patterns of the genes in our model. Results are found in Supplementary Data 15.

### MERFISH Spatial Transcriptomics

Tissue samples from mouse and axolotl limb were surgically dissected out and fixed overnight in a 4% PFA solution. Distal sections that included bone were additionally treated with a 10% formic acid solution overnight to enable decalcification. Distal sections in both organisms were taken at the position along the proximal-distal axis where the digits meet the hand/foot/paw, taken perpendicularly to the proximal-distal axis, capturing the complete anterior-posterior and dorsal-ventral axes. In both organisms, proximal samples were taken from the ventral side of the limb, slightly proximal to the knee. In both organisms, medial sections were taken from the ventral side of the limb midway between the knee and ankle joints. All sections should include and enable the visualization of all major cell types present in the limb with the exception of bone in proximal and medial sections, as it was excluded. Samples were then embedded in paraffin and sectioned to 5μm thickness by the Histology Core in the Koch Institute. Sections were captured on Vizgen MERFISH coverslips and processed according to the manufacturer Vizgen MERFISH 2.0 Sample Preparation User Guide for Sectioned Tissue Sample with a modified clearing protocol (5mL Vizgen Clearing Premix + 100μl Proteinase K (NEB #P8107S) at 47C for 19 hours, then 200ul Vizgen Clearing Premix + 50μl Proteinase K at 47C for 5 hours, then 5mL Vizgen Clearing Premix + 50μl Proteinase K at 37C for 72 hours). Probesets were generated by Vizgen. Some highly expressed genes were imaged with sequential FISH, done alongside MERFISH, identified by the lack of discrete dots in the presented images. MERFISH 1.0 images are shown for some other highly expressed genes. All imaging was done on a Vizgen MERSCOPE according to the manufacturer instructions. Segmentation was performed using CellPose^194^ implemented through the Vizgen Post-processing Tool.

Cellpose segmented cells were imported into Seurat for quantification. Cells with <3 unique genes or <8 detected transcripts were excluded. Remaining cells were clustered (Louvain) and visualized using UMAP. Cluster identities were assigned using known markers of mouse and axolotl cell types included in the MERFISH probeset for the purposes of cell-type identification. Differential expression analysis was performed between groups of computationally isolated cells using the binomial test within the FindMarkers function in Seurat. Quantification results and details can be found in Supplementary Data 17.

### 10X Genomics Visium HD Spatial Transcriptomics

Tissue samples from mouse limb were surgically dissected out and fixed overnight in a 4% PFA solution. All sections were taken perpendicular to the proximal-distal axis of the limb to include the full anterior-posterior and dorsal-ventral axes of the limb at that position and therefore all included bone. Because of the inclusion of bone, samples were additionally treated with 10% formic acid solution overnight to enable decalcification. Samples were then embedded in paraffin and sectioned to 5μm thickness by the Histology Core in the Koch Institute and subsequently captured on glass slides. Relevant tissue sections were identified, processed, and hybridized to transcriptomic probes by the Spatial Technology Platform Team at the Broad Institute using a Visium CytAssist following manufacturer instructions. Libraries were generated following standard Visium workflow and sequenced by NovaSeq 6000 S4. Visium Mouse Transcriptome Probe Set v2.0 was used to assess gene expression in the mouse samples. For computational analysis, filtered feature matrices of transcript counts for 8μm Visium HD bins were imported into Seurat. Bins with <100 UMIs and/or <50 unique genes detected were excluded. Clustering, cell-type identification, visualization, and differential expression analyses were done in the same way as for scRNA-seq datasets, described above.

### Predicted Protein Structures

Predicted structures of proteins encoded by spatially expressed unannotated genes were generated by ColabFold^195^. Highest ranked folding predictions were used. None of the unannotated genes had characterized hits by BLAST and none had any results with E-values < 1e-10 when searching for structure homology by FoldSeek^196^ and were therefore not shown. *Proxima* ortholog structures were also predicted and generated by Colabfold. Visualizations and alignments were done in PyMOL.

## Acknowledgements

We thank all members of the Reddien lab for continuous input and M.L. Scimone, Y. Huang, and K.D. Atabay for comments on the manuscript. Many thanks to O. Paugois for establishing and maintaining the axolotl housing system. We are appreciative of P. Aoude and K.D. Atabay for their input on data processing. We are grateful to A. Connor and the Preclinical Modeling Facility, as well as R. Flannery, for mouse samples. Many thanks to S. McCallum and A. Nutter-Upham for their extensive work implementing digilimb.wi.mit.edu. We thank T. Lopez for tissue processing input, S. Gupta and the Whitehead Institute Genome Technology Core for assistance with sequencing and data processing, S. Tsai for axolotl surgery training, as well as C. Rausch for illustrations. We are additionally grateful to the University of Kentucky Ambystoma Genetic Stock Center, and the NIH grant (P40-OD019794) that supports the center, for supplying axolotls. Thank you to the Koch Histology Core for help in tissue processing, as well as the members of the Broad Spatial Technology Platform for assistance with Visium HD. We thank the LEO Foundation, Eleanor Schwartz Charitable Foundation, and Howard Hughes Medical Institute for their financial support.

## Data availability

Bulk RNA-seq and scRNA-seq data have been deposited in the NCBI Sequence Read Archive under the BioProject accession IDs of PRJNA1358705 and PRJNA1363939, respectively. Processed scRNA-seq Seurat objects are available on Zenodo: 10.5281/zenodo.18316185. Interactive versions of the scRNA-seq and select MERFISH data are available at digilimb.wi.mit.edu.

## References

1 Brockes, J. P., Kumar, A. & Velloso, C. P. Regeneration as an evolutionary variable. J Anat 199, 3–11 (2001).

2 Sánchez Alvarado, A. Regeneration in the Metazoans: Why does it happen? BioEssays 22, 578–590 (2000).

3 Srivastava, M. Beyond Casual Resemblance: Rigorous Frameworks for Comparing Regeneration Across Species. Annu Rev Cell Dev Biol 37, 415–440 (2021). 10.1146/annurev-cellbio-120319-114716

4 Raz, A. A., Srivastava, M., Salvamoser, R. & Reddien, P. W. Acoel regeneration mechanisms indicate an ancient role for muscle in regenerative patterning. Nat Commun 8, 1260 (2017). 10.1038/s41467-017-01148-5

5 Srivastava, M., Mazza-Curll, K. L., van Wolfswinkel, J. C. & Reddien, P. W. Whole-body acoel regeneration is controlled by Wnt and Bmp-Admp signaling. Current biology : CB 24, 1107–1113 (2014). 10.1016/j.cub.2014.03.042

6 Gehrke, A. R. & Srivastava, M. Neoblasts and the evolution of whole-body regeneration. Curr Opin Genet Dev 40, 131–137 (2016). 10.1016/j.gde.2016.07.009

7 Fumagalli, M. R., Zapperi, S. & La Porta, C. A. M. Regeneration in distantly related species: common strategies and pathways. npj Systems Biology and Applications **4**, 5 (2018). 10.1038/s41540-017-0042-z

8 Darnet, S. et al. Deep evolutionary origin of limb and fin regeneration. Proceedings of the National Academy of Sciences 116, 15106–15115 (2019). doi:10.1073/pnas.1900475116

9 Wolpert, L. Positional information and the spatial pattern of cellular differentiation. Journal of theoretical biology 25, 1–47 (1969).

10 Wolpert, L. Positional information revisited. *Development* **107 Suppl**, 3–12 (1989). 10.1242/dev.107.Supplement.3

11 Reddien, P. W. The Cellular and Molecular Basis for Planarian Regeneration. Cell 175, 327–345 (2018). 10.1016/j.cell.2018.09.021

12 Broun, M., Gee, L., Reinhardt, B. & Bode, H. R. Formation of the head organizer in hydra involves the canonical Wnt pathway. Development 132, 2907–2916 (2005). 10.1242/dev.01848

13 Echeverri, K. & Tanaka, E. M. Proximodistal patterning during limb regeneration. Dev Biol 279, 391–401 (2005). 10.1016/j.ydbio.2004.12.029

14 Nacu, E., Gromberg, E., Oliveira, C. R., Drechsel, D. & Tanaka, E. M. FGF8 and SHH substitute for anterior-posterior tissue interactions to induce limb regeneration. Nature 533, 407–410 (2016). 10.1038/nature17972

15 Oliveira, C. R. et al. Tig1 regulates proximo-distal identity during salamander limb regeneration. Nature Communications 13, 1141 (2022). 10.1038/s41467-022-28755-1

16 Otsuki, L., Plattner, S. A., Taniguchi-Sugiura, Y., Falcon, F. & Tanaka, E. M. Molecular basis of positional memory in limb regeneration. Nature 642, 730–738 (2025). 10.1038/s41586-025-09036-5

17 Billingham, R. E. & Silvers, W. K. Studies on the conservation of epidermal specifies of skin and certain mucosas in adult mammals. J Exp Med 125, 429–446 (1967). 10.1084/jem.125.3.429

18 Dhouailly, D. Formation of cutaneous appendages in dermo-epidermal recombinations between reptiles, birds and mammals. Wilehm Roux Arch Dev Biol 177, 323–340 (1975). 10.1007/bf00848183

19 Sengel, P. Feather pattern development. Ciba Found Symp **0**, 51–70 (1975). 10.1002/9780470720110.ch4

20 Bohnert, A., Hornung, J., Mackenzie, I. C. & Fusenig, N. E. Epithelial-mesenchymal interactions control basement membrane production and differentiation in cultured and transplanted mouse keratinocytes. Cell and Tissue Research 244, 413–429 (1986). 10.1007/bf00219217

21 Sengel, P. Pattern formation in skin development. Int J Dev Biol 34, 33–50 (1990).

22 Yamaguchi, Y., Morita, A., Maeda, A. & Hearing, V. J. Regulation of skin pigmentation and thickness by Dickkopf 1 (DKK1). J Investig Dermatol Symp Proc 14, 73–75 (2009). 10.1038/jidsymp.2009.4

23 Yamaguchi, Y. et al. Regulation of keratin 9 in nonpalmoplantar keratinocytes by palmoplantar fibroblasts through epithelial-mesenchymal interactions. J Invest Dermatol 112, 483–488 (1999). 10.1046/j.1523-1747.1999.00544.x

24 Szabowski, A. et al. c-Jun and JunB antagonistically control cytokine-regulated mesenchymal-epidermal interaction in skin. Cell 103, 745–755 (2000). 10.1016/s0092-8674(00)00178-1

25 Shoshkes-Carmel, M. et al. Subepithelial telocytes are an important source of Wnts that supports intestinal crypts. Nature 557, 242–246 (2018). 10.1038/s41586-018-0084-4

26 McCarthy, N. et al. Distinct Mesenchymal Cell Populations Generate the Essential Intestinal BMP Signaling Gradient. Cell Stem Cell 26, 391–402.e395 (2020). 10.1016/j.stem.2020.01.008

27 Higuchi, Y. et al. Gastrointestinal Fibroblasts Have Specialized, Diverse Transcriptional Phenotypes: A Comprehensive Gene Expression Analysis of Human Fibroblasts. PLoS ONE 10, e0129241 (2015).

28 Chang, H. Y. et al. Diversity, topographic differentiation, and positional memory in human fibroblasts. Proc Natl Acad Sci U S A 99, 12877–12882 (2002). 10.1073/pnas.162488599

29 Rinn, J. L., Bondre, C., Gladstone, H. B., Brown, P. O. & Chang, H. Y. Anatomic demarcation by positional variation in fibroblast gene expression programs. PLoS genetics 2, e119 (2006). 10.1371/journal.pgen.0020119

30 Rinn, J. L. et al. Functional demarcation of active and silent chromatin domains in human HOX loci by noncoding RNAs. Cell 129, 1311–1323 (2007). 10.1016/j.cell.2007.05.022

31 Rinn, J. L. et al. A dermal HOX transcriptional program regulates site-specific epidermal fate. Genes & development 22, 303–307 (2008). 10.1101/gad.1610508

32 Johnson, G. L., Masias, E. J. & Lehoczky, J. A. Cellular Heterogeneity and Lineage Restriction during Mouse Digit Tip Regeneration at Single-Cell Resolution. Dev Cell 52, 525–540.e525 (2020). 10.1016/j.devcel.2020.01.026

33 Storer, M. A. et al. Acquisition of a Unique Mesenchymal Precursor-like Blastema State Underlies Successful Adult Mammalian Digit Tip Regeneration. Dev Cell 52, 509–524.e509 (2020). 10.1016/j.devcel.2019.12.004

34 Mahmud, N. et al. Nail-associated mesenchymal cells contribute to and are essential for dorsal digit tip regeneration. Cell Rep 41, 111853 (2022). 10.1016/j.celrep.2022.111853

35 Nachtrab, G., Kikuchi, K., Tornini, V. A. & Poss, K. D. Transcriptional components of anteroposterior positional information during zebrafish fin regeneration. Development 140, 3754–3764 (2013). 10.1242/dev.098798

36 Kragl, M. et al. Cells keep a memory of their tissue origin during axolotl limb regeneration. Nature 460, 60–65 (2009). nature08152 [pii] 10.1038/nature08152

37 Nacu, E. et al. Connective tissue cells, but not muscle cells, are involved in establishing the proximo-distal outcome of limb regeneration in the axolotl. Development 140, 513–518 (2013). 10.1242/dev.081752

38 Kawaguchi, A. et al. A chromatin code for limb segment identity in axolotl limb regeneration. Developmental Cell 59, 2239–2253.e2239 (2024). 10.1016/j.devcel.2024.05.002

39 Reddien, P. W. Constitutive gene expression and the specification of tissue identity in adult planarian biology. Trends Genet 27, 277–285 (2011). S0168-9525(11)00060-6 [pii] 10.1016/j.tig.2011.04.004

40 Witchley, J. N., Mayer, M., Wagner, D. E., Owen, J. H. & Reddien, P. W. Muscle cells provide instructions for planarian regeneration. Cell Reports 4, 633–641 (2013). 10.1016/j.celrep.2013.07.022

41 Scimone, M. L., Cote, L. E. & Reddien, P. W. Orthogonal muscle fibres have different instructive roles in planarian regeneration. Nature 551, 623–628 (2017). 10.1038/nature24660

42 Tewari, A. G., Stern, S. R., Oderberg, I. M. & Reddien, P. W. Cellular and Molecular Responses Unique to Major Injury Are Dispensable for Planarian Regeneration. Cell Reports 25, 2577–2590 e2573 (2018). 10.1016/j.celrep.2018.11.004

43 Scimone, M. L., Cloutier, J. K., Maybrun, C. L. & Reddien, P. W. The planarian wound epidermis gene equinox is required for blastema formation in regeneration. Nat Commun 13, 2726 (2022). 10.1038/s41467-022-30412-6

44 Owlarn, S. & Bartscherer, K. Go ahead, grow a head! A planarian’s guide to anterior regeneration. Regeneration 3, 139–155 (2016).

45 Ruiz-Trillo, I., Riutort, M., Littlewood, D. T., Herniou, E. A. & Baguna, J. Acoel flatworms: earliest extant bilaterian Metazoans, not members of Platyhelminthes. Science 283, 1919–1923. (1999).

46 Hejnol, A. et al. Assessing the root of bilaterian animals with scalable phylogenomic methods. Proc Biol Sci 276, 4261–4270 (2009). 10.1098/rspb.2009.0896

47 Cannon, J. T. et al. Xenacoelomorpha is the sister group to Nephrozoa. Nature 530, 89–93 (2016). 10.1038/nature16520

48 Arroyo, A. S., López-Escardó, D., Vargas, C. d. & Ruiz-Trillo, I. Hidden diversity of Acoelomorpha revealed through metabarcoding. Biol. Lett. 12, 20160674 (2016).

49 Rouse, G. W., Wilson, N. G., Carvajal, J. I. & Vrijenhoek, R. C. New deep-sea species of Xenoturbella and the position of Xenacoelomorpha. Nature 530, 94–97 (2016). 10.1038/nature16545

50 Cote, L. E., Simental, E. & Reddien, P. W. Muscle functions as a connective tissue and source of extracellular matrix in planarians. Nat Commun 10, 1592 (2019). 10.1038/s41467-019-09539-6

51 Bryant, D. M. et al. A Tissue-Mapped Axolotl De Novo Transcriptome Enables Identification of Limb Regeneration Factors. Cell Rep 18, 762–776 (2017). 10.1016/j.celrep.2016.12.063

52 Plikus, M. V. et al. Fibroblasts: Origins, definitions, and functions in health and disease. Cell 184, 3852–3872 (2021). 10.1016/j.cell.2021.06.024

53 Carr, M. J. et al. Mesenchymal Precursor Cells in Adult Nerves Contribute to Mammalian Tissue Repair and Regeneration. Cell Stem Cell 24, 240–256.e249 (2019). 10.1016/j.stem.2018.10.024

54 Muhl, L. et al. Single-cell analysis uncovers fibroblast heterogeneity and criteria for fibroblast and mural cell identification and discrimination. Nature Communications 11, 3953 (2020). 10.1038/s41467-020-17740-1

55 He, H. et al. Single-cell transcriptome analysis of human skin identifies novel fibroblast subpopulation and enrichment of immune subsets in atopic dermatitis. J Allergy Clin Immunol 145, 1615–1628 (2020). 10.1016/j.jaci.2020.01.042

56 Ascensión, A. M., Fuertes-Álvarez, S., Ibañez-Solé, O., Izeta, A. & Araúzo-Bravo, M. J. Human Dermal Fibroblast Subpopulations Are Conserved across Single-Cell RNA Sequencing Studies. J Invest Dermatol 141, 1735–1744.e1735 (2021). 10.1016/j.jid.2020.11.028

57 Buechler, M. B. et al. Cross-tissue organization of the fibroblast lineage. Nature 593, 575–579 (2021). 10.1038/s41586-021-03549-5

58 Deng, C. C. et al. Single-cell RNA-seq reveals fibroblast heterogeneity and increased mesenchymal fibroblasts in human fibrotic skin diseases. Nat Commun 12, 3709 (2021). 10.1038/s41467-021-24110-y

59 Solé-Boldo, L. et al. Single-cell transcriptomes of the human skin reveal age-related loss of fibroblast priming. Communications Biology 3, 188 (2020). 10.1038/s42003-020-0922-4

60 McKee, T. J., Perlman, G., Morris, M. & Komarova, S. V. Extracellular matrix composition of connective tissues: a systematic review and meta-analysis. Sci Rep 9, 10542 (2019). 10.1038/s41598-019-46896-0

61 Moffitt, J. R. et al. High-throughput single-cell gene-expression profiling with multiplexed error-robust fluorescence in situ hybridization. Proceedings of the National Academy of Sciences 113, 11046–11051 (2016). doi:10.1073/pnas.1612826113

62 Chen, K. H., Boettiger, A. N., Moffitt, J. R., Wang, S. & Zhuang, X. Spatially resolved, highly multiplexed RNA profiling in single cells. Science 348, aaa6090 (2015). doi:10.1126/science.aaa6090

63 Philippeos, C. et al. Spatial and Single-Cell Transcriptional Profiling Identifies Functionally Distinct Human Dermal Fibroblast Subpopulations. J Invest Dermatol 138, 811–825 (2018). 10.1016/j.jid.2018.01.016

64 Driskell, R. R. & Watt, F. M. Understanding fibroblast heterogeneity in the skin. Trends Cell Biol 25, 92–99 (2015). 10.1016/j.tcb.2014.10.001

65 Martino, P. A., Heitman, N. & Rendl, M. The dermal sheath: An emerging component of the hair follicle stem cell niche. Exp Dermatol 30, 512–521 (2021). 10.1111/exd.14204

66 Chou, C.-H. et al. Synovial cell cross-talk with cartilage plays a major role in the pathogenesis of osteoarthritis. Scientific Reports 10, 10868 (2020). 10.1038/s41598-020-67730-y

67 Zhang, Z. et al. Synovial fibroblast derived small extracellular vesicles miRNA15- 29148 promotes articular chondrocyte apoptosis in rheumatoid arthritis. Bone Research 13, 61 (2025). 10.1038/s41413-025-00430-3

68 Sebastian, A. et al. Single-Cell RNA-Seq Reveals Transcriptomic Heterogeneity and Post-Traumatic Osteoarthritis-Associated Early Molecular Changes in Mouse Articular Chondrocytes. Cells 10 (2021). 10.3390/cells10061462

69 Logan, M. & Tabin, C. J. Role of Pitx1 upstream of Tbx4 in specification of hindlimb identity. Science 283, 1736–1739 (1999). 10.1126/science.283.5408.1736

70 Logan, M., Simon, H. & Tabin, C. Differential regulation of T-box and homeobox transcription factors suggests roles in controlling chick limb-type identity. Development 125, 2825–2835 (1998).

71 Rallis, C. et al. Tbx5 is required for forelimb bud formation and continued outgrowth. Development 130, 2741–2751 (2003). 10.1242/dev.00473

72 Nemec, S. et al. Pitx1 directly modulates the core limb development program to implement hindlimb identity. Development 144, 3325–3335 (2017). 10.1242/dev.154864

73 Simon, H.-G. et al. A novel family of T-box genes in urodele amphibian limb development and regeneration: candidate genes involved in vertebrate forelimb/hindlimb patterning. Development 124, 1355–1366 (1997). 10.1242/dev.124.7.1355

74 Gibson-Brown, J. J. et al. Evidence of a role for T-box genes in the evolution of limb morphogenesis and the specification of forelimb/hindlimb identity. Mech Dev 56, 93–101 (1996). 10.1016/0925-4773(96)00514-x

75 Takeuchi, J. K. et al. Tbx5 and Tbx4 trigger limb initiation through activation of the Wnt/Fgf signaling cascade. Development 130, 2729–2739 (2003). 10.1242/dev.00474

76 Naiche, L. A. & Papaioannou, V. E. Loss of Tbx4 blocks hindlimb development and affects vascularization and fusion of the allantois. Development 130, 2681–2693 (2003). 10.1242/dev.00504

77 Carlson, M. R., Komine, Y., Bryant, S. V. & Gardiner, D. M. Expression of Hoxb13 and Hoxc10 in developing and regenerating Axolotl limbs and tails. Dev Biol 229, 396–406 (2001). 10.1006/dbio.2000.0104

78 Chen, L. et al. Heterozygous deletion of *HOXC10-HOXC9* causes lower limb abnormalities in congenital vertical talus. Journal of Medical Genetics 61, 777–779 (2024). 10.1136/jmg-2023-109656

79 Peterson, R. L., Papenbrock, T., Davda, M. M. & Awgulewitsch, A. The murine Hoxc cluster contains five neighboring AbdB-related Hox genes that show unique spatially coordinated expression in posterior embryonic subregions. Mech Dev 47, 253–260 (1994). 10.1016/0925-4773(94)90043-4

80 Hubert, K. A. & Wellik, D. M. Hox genes in development and beyond. Development 150 (2023). 10.1242/dev.192476

81 Krumlauf, R. Hox genes in vertebrate development. Cell 78, 191–201 (1994). 10.1016/0092-8674(94)90290-9

82 Peraldi, R. & Kmita, M. 40 years of the homeobox: mechanisms of Hox spatial-temporal collinearity in vertebrates. Development 151 (2024). 10.1242/dev.202508

83 Wellik, D. M. & Capecchi, M. R. Hox10 and Hox11 Genes Are Required to Globally Pattern the Mammalian Skeleton. Science 301, 363–367 (2003). doi:10.1126/science.1085672

84 Wellik, D. M., Hawkes, P. J. & Capecchi, M. R. Hox11 paralogous genes are essential for metanephric kidney induction. Genes Dev 16, 1423–1432 (2002). 10.1101/gad.993302

85 Boulet, A. M. & Capecchi, M. R. Multiple roles of Hoxa11 and Hoxd11 in the formation of the mammalian forelimb zeugopod. Development 131, 299–309 (2004). 10.1242/dev.00936

86 Davis, A. P., Witte, D. P., Hsieh-Li, H. M., Potter, S. S. & Capecchi, M. R. Absence of radius and ulna in mice lacking hoxa-11 and hoxd-11. Nature 375, 791–795 (1995). 10.1038/375791a0

87 Fromental-Ramain, C. et al. Hoxa-13 and Hoxd-13 play a crucial role in the patterning of the limb autopod. Development 122, 2997–3011 (1996). 10.1242/dev.122.10.2997

88 Kmita, M., Fraudeau, N., Hérault, Y. & Duboule, D. Serial deletions and duplications suggest a mechanism for the collinearity of Hoxd genes in limbs. Nature 420, 145–150 (2002). 10.1038/nature01189

89. Tickle, C. in *HOX Gene Expression* 42-52 (Springer New York, 2007).

90 Cooper, K. L. et al. Initiation of Proximal-Distal Patterning in the Vertebrate Limb by Signals and Growth. Science 332, 1083–1086 (2011). doi:10.1126/science.1199499

91 Fromental-Ramain, C. et al. Specific and redundant functions of the paralogous Hoxa-9 and Hoxd-9 genes in forelimb and axial skeleton patterning. Development 122, 461–472 (1996). 10.1242/dev.122.2.461

92 Kmita, M. et al. Early developmental arrest of mammalian limbs lacking HoxA/HoxD gene function. Nature 435, 1113–1116 (2005). 10.1038/nature03648

93 Tarchini, B., Duboule, D. & Kmita, M. Regulatory constraints in the evolution of the tetrapod limb anterior–posterior polarity. Nature 443, 985–988 (2006). 10.1038/nature05247

94 Roensch, K., Tazaki, A., Chara, O. & Tanaka, E. M. Progressive specification rather than intercalation of segments during limb regeneration. Science 342, 1375–1379 (2013). 10.1126/science.1241796

95 Quigley, I. K., Stubbs, J. L. & Kintner, C. Specification of ion transport cells in the Xenopus larval skin. Development 138, 705–714 (2011). 10.1242/dev.055699

96 Fallon, J. F. et al. FGF-2: Apical Ectodermal Ridge Growth Signal for Chick Limb Development. Science 264, 104–107 (1994). doi:10.1126/science.7908145

97 Niswander, L. & Martin, G. R. FGF-4 and BMP-2 have opposite effects on limb growth. Nature 361, 68–71 (1993). 10.1038/361068a0

98 Niswander, L., Tickle, C., Vogel, A., Booth, I. & Martin, G. R. FGF-4 replaces the apical ectodermal ridge and directs outgrowth and patterning of the limb. Cell 75, 579–587 (1993). 10.1016/0092-8674(93)90391-3

99 Capdevila, J., Tsukui, T., Rodríquez Esteban, C., Zappavigna, V. & Izpisúa Belmonte, J. C. Control of vertebrate limb outgrowth by the proximal factor Meis2 and distal antagonism of BMPs by Gremlin. Mol Cell 4, 839–849 (1999). 10.1016/s1097-2765(00)80393-7

100 Mercader, N. et al. Opposing RA and FGF signals control proximodistal vertebrate limb development through regulation of Meis genes. Development 127, 3961–3970 (2000). 10.1242/dev.127.18.3961

101 Mariani, F. V., Ahn, C. P. & Martin, G. R. Genetic evidence that FGFs have an instructive role in limb proximal–distal patterning. Nature 453, 401–405 (2008). 10.1038/nature06876

102 ten Berge, D., Brugmann, S. A., Helms, J. A. & Nusse, R. Wnt and FGF signals interact to coordinate growth with cell fate specification during limb development. Development 135, 3247–3257 (2008). 10.1242/dev.023176

103 Sekine, K. et al. Fgf10 is essential for limb and lung formation. Nat Genet 21, 138–141 (1999). 10.1038/5096

104 Ohuchi, H. et al. The mesenchymal factor, FGF10, initiates and maintains the outgrowth of the chick limb bud through interaction with FGF8, an apical ectodermal factor. Development 124, 2235–2244 (1997). 10.1242/dev.124.11.2235

105 Kawakami, Y. et al. WNT signals control FGF-dependent limb initiation and AER induction in the chick embryo. Cell 104, 891–900 (2001). 10.1016/s0092-8674(01)00285-9

106 Lewandoski, M., Sun, X. & Martin, G. R. Fgf8 signalling from the AER is essential for normal limb development. Nat Genet 26, 460–463 (2000). 10.1038/82609

107 Barrow, J. R. et al. Ectodermal Wnt3/beta-catenin signaling is required for the establishment and maintenance of the apical ectodermal ridge. Genes Dev 17, 394–409 (2003). 10.1101/gad.1044903

108 Kengaku, M. et al. Distinct WNT Pathways Regulating AER Formation and Dorsoventral Polarity in the Chick Limb Bud. Science 280, 1274–1277 (1998). doi:10.1126/science.280.5367.1274

109 Dealy, C. N., Roth, A., Ferrari, D., Brown, A. M. & Kosher, R. A. Wnt-5a and Wnt- 7a are expressed in the developing chick limb bud in a manner suggesting roles in pattern formation along the proximodistal and dorsoventral axes. Mech Dev 43, 175–186 (1993). 10.1016/0925-4773(93)90034-u

110 Yamaguchi, T. P., Bradley, A., McMahon, A. P. & Jones, S. A Wnt5a pathway underlies outgrowth of multiple structures in the vertebrate embryo. Development 126, 1211–1223 (1999). 10.1242/dev.126.6.1211

111 Saunders, J. W., Jr. The proximo-distal sequence of origin of the parts of the chick wing and the role of the ectoderm. J Exp Zool 108, 363–403 (1948). 10.1002/jez.1401080304

112 Zúñiga, A. Next generation limb development and evolution: old questions, new perspectives. Development 142, 3810–3820 (2015). 10.1242/dev.125757

113 McQueen, C. & Towers, M. Establishing the pattern of the vertebrate limb. Development 147 (2020). 10.1242/dev.177956

114 Kawakami, Y. et al. Sp8 and Sp9, two closely related buttonhead-like transcription factors, regulate Fgf8expression and limb outgrowth in vertebrate embryos. Development 131, 4763–4774 (2004). 10.1242/dev.01331

115 Haro, E. et al. Sp6 and Sp8 transcription factors control AER formation and dorsal-ventral patterning in limb development. PLoS Genet 10, e1004468 (2014). 10.1371/journal.pgen.1004468

116 Zhong, J. et al. Multi-species atlas resolves an axolotl limb development and regeneration paradox. Nature Communications 14, 6346 (2023). 10.1038/s41467-023-41944-w

117 Glotzer, G. L., Tardivo, P. & Tanaka, E. M. Canonical Wnt signaling and the regulation of divergent mesenchymal Fgf8 expression in axolotl limb development and regeneration. Elife 11 (2022). 10.7554/eLife.79762

118 Christensen, R. N., Weinstein, M. & Tassava, R. A. Expression of fibroblast growth factors 4, 8, and 10 in limbs, flanks, and blastemas of Ambystoma. Dev Dyn 223, 193–203 (2002). 10.1002/dvdy.10049

119 Purushothaman, S., Elewa, A. & Seifert, A. W. Fgf-signaling is compartmentalized within the mesenchyme and controls proliferation during salamander limb development. Elife 8 (2019). 10.7554/eLife.48507

120 Zúñiga, A., Haramis, A. P., McMahon, A. P. & Zeller, R. Signal relay by BMP antagonism controls the SHH/FGF4 feedback loop in vertebrate limb buds. Nature 401, 598–602 (1999). 10.1038/44157

121 Michos, O. et al. Gremlin-mediated BMP antagonism induces the epithelial-mesenchymal feedback signaling controlling metanephric kidney and limb organogenesis. Development 131, 3401–3410 (2004). 10.1242/dev.01251

122 Khokha, M. K., Hsu, D., Brunet, L. J., Dionne, M. S. & Harland, R. M. Gremlin is the BMP antagonist required for maintenance of Shh and Fgf signals during limb patterning. Nature Genetics 34, 303–307 (2003). 10.1038/ng1178

123 Maden, M. Vitamin A and pattern formation in the regenerating limb. Nature 295, 672–675 (1982). 10.1038/295672a0

124 Roselló-Díez, A., Ros, M. A. & Torres, M. Diffusible signals, not autonomous mechanisms, determine the main proximodistal limb subdivision. Science 332, 1086–1088 (2011). 10.1126/science.1199489

125 Niederreither, K., Subbarayan, V., Dollé, P. & Chambon, P. Embryonic retinoic acid synthesis is essential for early mouse post-implantation development. Nat Genet 21, 444–448 (1999). 10.1038/7788

126 Mic, F. A., Sirbu, I. O. & Duester, G. Retinoic acid synthesis controlled by Raldh2 is required early for limb bud initiation and then later as a proximodistal signal during apical ectodermal ridge formation. J Biol Chem 279, 26698–26706 (2004). 10.1074/jbc.M401920200

127 Duester, G. Retinoic acid synthesis and signaling during early organogenesis. Cell 134, 921–931 (2008). 10.1016/j.cell.2008.09.002

128 Cunningham, T. J., Chatzi, C., Sandell, L. L., Trainor, P. A. & Duester, G. Rdh10 mutants deficient in limb field retinoic acid signaling exhibit normal limb patterning but display interdigital webbing. Dev Dyn 240, 1142–1150 (2011). 10.1002/dvdy.22583

129 Sandell, L. L. et al. RDH10 is essential for synthesis of embryonic retinoic acid and is required for limb, craniofacial, and organ development. Genes Dev 21, 1113–1124 (2007). 10.1101/gad.1533407

130 Yashiro, K. et al. Regulation of retinoic acid distribution is required for proximodistal patterning and outgrowth of the developing mouse limb. Dev Cell 6, 411–422 (2004). 10.1016/s1534-5807(04)00062-0

131 Roselló-Díez, A., Arques, C. G., Delgado, I., Giovinazzo, G. & Torres, M. Diffusible signals and epigenetic timing cooperate in late proximo-distal limb patterning. Development 141, 1534–1543 (2014). 10.1242/dev.106831

132 Duerr, T. J. et al. Retinoic acid breakdown is required for proximodistal positional identity during axolotl limb regeneration. Nature Communications 16, 4798 (2025). 10.1038/s41467-025-59497-5

133 Mercader, N. et al. Conserved regulation of proximodistal limb axis development by Meis1/Hth. Nature 402, 425–429 (1999). 10.1038/46580

134 Mercader, N. et al. Ectopic Meis1 expression in the mouse limb bud alters P-D patterning in a Pbx1-independent manner. Int J Dev Biol 53, 1483–1494 (2009). 10.1387/ijdb.072430nm

135 Delgado, I. et al. Proximo-distal positional information encoded by an Fgf-regulated gradient of homeodomain transcription factors in the vertebrate limb. Science Advances 6, eaaz0742 (2020). doi:10.1126/sciadv.aaz0742

136 Mercader, N., Tanaka, E. M. & Torres, M. Proximodistal identity during vertebrate limb regeneration is regulated by Meis homeodomain proteins. Development 132, 4131–4142 (2005). 10.1242/dev.01976

137 Gerber, T. et al. Single-cell analysis uncovers convergence of cell identities during axolotl limb regeneration. Science 362 (2018). 10.1126/science.aaq0681

138 Satoh, W., Gotoh, T., Tsunematsu, Y., Aizawa, S. & Shimono, A. Sfrp1 and Sfrp2 regulate anteroposterior axis elongation and somite segmentation during mouse embryogenesis. Development 133, 989–999 (2006). 10.1242/dev.02274

139 Ikegawa, M. et al. Syndactyly and preaxial synpolydactyly in the single Sfrp2 deleted mutant mice. Dev Dyn 237, 2506–2517 (2008). 10.1002/dvdy.21655

140 Morello, R. et al. Brachy-syndactyly caused by loss of Sfrp2 function. J Cell Physiol 217, 127–137 (2008). 10.1002/jcp.21483

141 Singh, M. K. et al. The T-box transcription factor Tbx15 is required for skeletal development. Mech Dev 122, 131–144 (2005). 10.1016/j.mod.2004.10.011

142 Allen, J. M., McGlinn, E., Hill, A. & Warman, M. L. Autopodial development is selectively impaired by misexpression of chordin-like 1 in the chick limb. Dev Biol 381, 159–169 (2013). 10.1016/j.ydbio.2013.06.003

143 da Silva, S. M., Gates, P. B. & Brockes, J. P. The newt ortholog of CD59 is implicated in proximodistal identity during amphibian limb regeneration. Dev Cell 3, 547–555 (2002). 10.1016/s1534-5807(02)00288-5

144 Kumar, A., Godwin, J. W., Gates, P. B., Garza-Garcia, A. A. & Brockes, J. P. Molecular basis for the nerve dependence of limb regeneration in an adult vertebrate. Science 318, 772–777 (2007). 10.1126/science.1147710

145 Shaikh, N., Gates, P. B. & Brockes, J. P. The Meis homeoprotein regulates the axolotl Prod 1 promoter during limb regeneration. Gene 484, 69–74 (2011). 10.1016/j.gene.2011.06.003

146 Steinert, P. M., Parry, D. A. & Marekov, L. N. Trichohyalin mechanically strengthens the hair follicle: multiple cross-bridging roles in the inner root shealth. J Biol Chem 278, 41409–41419 (2003). 10.1074/jbc.M302037200

147 Kościuczuk, E. M. et al. Cathelicidins: family of antimicrobial peptides. A review. Mol Biol Rep 39, 10957–10970 (2012). 10.1007/s11033-012-1997-x

148 Jackson, B. et al. Late cornified envelope family in differentiating epithelia-- response to calcium and ultraviolet irradiation. J Invest Dermatol 124, 1062–1070 (2005). 10.1111/j.0022-202X.2005.23699.x

149 Archer, N. K., Dilolli, M. N. & Miller, L. S. Pushing the Envelope in Psoriasis: Late Cornified Envelope Proteins Possess Antimicrobial Activity. J Invest Dermatol 137, 2257–2259 (2017). 10.1016/j.jid.2017.08.026

150 Zhang, C. et al. Small proline-rich proteins (SPRRs) are epidermally produced antimicrobial proteins that defend the cutaneous barrier by direct bacterial membrane disruption. Elife 11 (2022). 10.7554/eLife.76729

151 Luna, M. C., Mcdiarmid, R. W. & Faivovich, J. From erotic excrescences to pheromone shots: structure and diversity of nuptial pads in anurans. Biological Journal of the Linnean Society 124, 403–446 (2018). 10.1093/biolinnean/bly048

152 Qiu, F., Ai, Q., Li, J. & Wu, H. Transcriptome analysis reveals the genetic basis underlying the formation and seasonal changes of nuptial pads in Rana chensinensis. BMC Genomics 25, 1254 (2024). 10.1186/s12864-024-11176-3

153 Stocum, D. L. Regeneration of symmetrical hindlimbs in larval salamanders. Science 200, 790–793 (1978). 10.1126/science.644323

154 Tank, P. W. The failure of double-half forelimbs to undergo distal transformation following amputation in the axolotl, Ambystoma mexicanum. J Exp Zool 204, 325–336 (1978). 10.1002/jez.1402040303

155 Slack, J. M. & Savage, S. Regeneration of mirror symmetrical limbs in the axolotl. Cell 14, 1–8 (1978). 10.1016/0092-8674(78)90295-7

156 Meinhardt, H. A boundary model for pattern formation in vertebrate limbs. J Embryol Exp Morphol 76, 115–137 (1983).

157 Riddle, R. D. et al. Induction of the LIM homeobox gene Lmx1 by WNT7a establishes dorsoventral pattern in the vertebrate limb. Cell 83, 631–640 (1995). 10.1016/0092-8674(95)90103-5

158 Vogel, A., Rodriguez, C., Warnken, W. & Izpisúa Belmonte, J. C. Dorsal cell fate specified by chick Lmx1 during vertebrate limb development. Nature 378, 716–720 (1995). 10.1038/378716a0

159 Chen, H. et al. Limb and kidney defects in Lmx1b mutant mice suggest an involvement of LMX1B in human nail patella syndrome. Nat Genet 19, 51–55 (1998). 10.1038/ng0598-51

160 Maatouk, D. M., Choi, K. S., Bouldin, C. M. & Harfe, B. D. In the limb AER Bmp2 and Bmp4 are required for dorsal-ventral patterning and interdigital cell death but not limb outgrowth. Dev Biol 327, 516–523 (2009). 10.1016/j.ydbio.2009.01.004

161 Qiu, Q., Chen, H. & Johnson, R. L. Lmx1b-expressing cells in the mouse limb bud define a dorsal mesenchymal lineage compartment. Genesis 47, 224–233 (2009). 10.1002/dvg.20430

162 Loomis, C. A. et al. The mouse Engrailed-1 gene and ventral limb patterning. Nature 382, 360–363 (1996). 10.1038/382360a0

163 Loomis, C. A., Kimmel, R. A., Tong, C. X., Michaud, J. & Joyner, A. L. Analysis of the genetic pathway leading to formation of ectopic apical ectodermal ridges in mouse Engrailed-1 mutant limbs. Development 125, 1137–1148 (1998). 10.1242/dev.125.6.1137

164 Logan, C., Hornbruch, A., Campbell, I. & Lumsden, A. The role of Engrailed in establishing the dorsoventral axis of the chick limb. Development 124, 2317–2324 (1997). 10.1242/dev.124.12.2317

165 Pizette, S., Abate-Shen, C. & Niswander, L. BMP controls proximodistal outgrowth, via induction of the apical ectodermal ridge, and dorsoventral patterning in the vertebrate limb. Development 128, 4463–4474 (2001). 10.1242/dev.128.22.4463

166 Yamamoto, S., Kashimoto, R., Furukawa, S., Ohashi, A. & Satoh, A. Lmx1b activation in axolotl limb regeneration. Dev Dyn 251, 1509–1523 (2022). 10.1002/dvdy.476

167 Johnson, G. L., Glasser, M. B., Charles, J. F., Duryea, J. & Lehoczky, J. A. En1 and Lmx1b do not recapitulate embryonic dorsal-ventral limb patterning functions during mouse digit tip regeneration. Cell Rep 41, 111701 (2022). 10.1016/j.celrep.2022.111701

168 Qu, S. et al. Polydactyly and ectopic ZPA formation in Alx-4 mutant mice. Development 124, 3999–4008 (1997). 10.1242/dev.124.20.3999

169 Qu, S. et al. Mutations in mouse Aristaless-like4 cause Strong’s luxoid polydactyly. Development 125, 2711–2721 (1998). 10.1242/dev.125.14.2711

170 Takahashi, M. et al. The role of Alx-4 in the establishment of anteroposterior polarity during vertebrate limb development. Development 125, 4417–4425 (1998). 10.1242/dev.125.22.4417

171 Kuijper, S. et al. Function and regulation of Alx4 in limb development: complex genetic interactions with Gli3 and Shh. Dev Biol 285, 533–544 (2005). 10.1016/j.ydbio.2005.06.017

172 Echelard, Y. et al. Sonic hedgehog, a member of a family of putative signaling molecules, is implicated in the regulation of CNS polarity. Cell 75, 1417–1430 (1993). 10.1016/0092-8674(93)90627-3

173 Riddle, R. D., Johnson, R. L., Laufer, E. & Tabin, C. Sonic hedgehog mediates the polarizing activity of the ZPA. Cell 75, 1401–1416 (1993). 10.1016/0092-8674(93)90626-2

174 Chiang, C. et al. Cyclopia and defective axial patterning in mice lacking Sonic hedgehog gene function. Nature 383, 407–413 (1996). 10.1038/383407a0

175 Yang, Y. et al. Relationship between dose, distance and time in Sonic Hedgehog-mediated regulation of anteroposterior polarity in the chick limb. Development 124, 4393–4404 (1997). 10.1242/dev.124.21.4393

176 Duprez, D., Lapointe, F., Edom-Vovard, F., Kostakopoulou, K. & Robson, L. Sonic hedgehog (SHH) specifies muscle pattern at tissue and cellular chick level, in the chick limb bud. Mech Dev 82, 151–163 (1999). 10.1016/s0925-4773(99)00040-4

177 Charité, J., McFadden, D. G. & Olson, E. N. The bHLH transcription factor dHAND controls Sonic hedgehog expression and establishment of the zone of polarizing activity during limb development. Development 127, 2461–2470 (2000). 10.1242/dev.127.11.2461

178 Fernandez-Teran, M. et al. Role of dHAND in the anterior-posterior polarization of the limb bud: implications for the Sonic hedgehog pathway. Development 127, 2133–2142 (2000). 10.1242/dev.127.10.2133

179 Yelon, D. et al. The bHLH transcription factor hand2 plays parallel roles in zebrafish heart and pectoral fin development. Development 127, 2573–2582 (2000). 10.1242/dev.127.12.2573

180 Nelson, C. E. et al. Analysis of Hox gene expression in the chick limb bud. Development 122, 1449–1466 (1996). 10.1242/dev.122.5.1449

181 Imokawa, Y. & Yoshizato, K. Expression of Sonic hedgehog gene in regenerating newt limb blastemas recapitulates that in developing limb buds. Proc Natl Acad Sci U S A 94, 9159–9164 (1997). 10.1073/pnas.94.17.9159

182 Torok, M. A., Gardiner, D. M., Izpisúa-Belmonte, J. C. & Bryant, S. V. Sonic hedgehog (shh) expression in developing and regenerating axolotl limbs. J Exp Zool 284, 197–206 (1999).

183 Lapan, S. W. & Reddien, P. W. Transcriptome Analysis of the Planarian Eye Identifies ovo as a Specific Regulator of Eye Regeneration. Cell Reports 2, 294–307 (2012). 10.1016/j.celrep.2012.06.018

184 Storer, M. A. & Miller, F. D. Cellular and molecular mechanisms that regulate mammalian digit tip regeneration. Open Biol 10, 200194 (2020). 10.1098/rsob.200194

185 Borgens, R. B. Mice regrow the tips of their foretoes. Science 217, 747–750 (1982).

186 Johnson, G. L. & Lehoczky, J. A. Mammalian Digit Tip Regeneration: Moving from Phenomenon to Molecular Mechanism. Cold Spring Harb Perspect Biol 14 (2022). 10.1101/cshperspect.a040857

187 Kragl, M. & Tanaka, E. M. Axolotl (Ambystoma mexicanum) limb and tail amputation. Cold Spring Harb Protoc 2009, pdb.prot5267 (2009). 10.1101/pdb.prot5267

188 Love, M. I., Huber, W. & Anders, S. Moderated estimation of fold change and dispersion for RNA-seq data with DESeq2. Genome Biol 15, 550 (2014). 10.1186/s13059-014-0550-8

189 Fleming, S. J. et al. Unsupervised removal of systematic background noise from droplet-based single-cell experiments using CellBender. Nature Methods 20, 1323–1335 (2023). 10.1038/s41592-023-01943-7

190 Germain, P. L., Lun, A., Garcia Meixide, C., Macnair, W. & Robinson, M. D. Doublet identification in single-cell sequencing data using scDblFinder. F1000Res 10, 979 (2021). 10.12688/f1000research.73600.2

191 Satija, R., Farrell, J. A., Gennert, D., Schier, A. F. & Regev, A. Spatial reconstruction of single-cell gene expression data. Nat Biotechnol 33, 495–502 (2015). 10.1038/nbt.3192

192 Naba, A. et al. The matrisome: in silico definition and in vivo characterization by proteomics of normal and tumor extracellular matrices. Mol Cell Proteomics 11, M111.014647 (2012). 10.1074/mcp.M111.014647

193 Ravasi, T. et al. An atlas of combinatorial transcriptional regulation in mouse and man. Cell 140, 744–752 (2010). 10.1016/j.cell.2010.01.044

194 Stringer, C., Wang, T., Michaelos, M. & Pachitariu, M. Cellpose: a generalist algorithm for cellular segmentation. Nature Methods 18, 100–106 (2021). 10.1038/s41592-020-01018-x

195 Mirdita, M. et al. ColabFold: making protein folding accessible to all. Nature Methods 19, 679–682 (2022). 10.1038/s41592-022-01488-1

196 van Kempen, M. et al. Fast and accurate protein structure search with Foldseek. Nature Biotechnology 42, 243–246 (2024). 10.1038/s41587-023-01773-0

