## Supplementary Figures for "Spatial gene expression maps in vertebrate limbs display conserved and regenerative species-specific features within connective tissue"

### Supplementary Fig. 1 (Fig. 1S1)

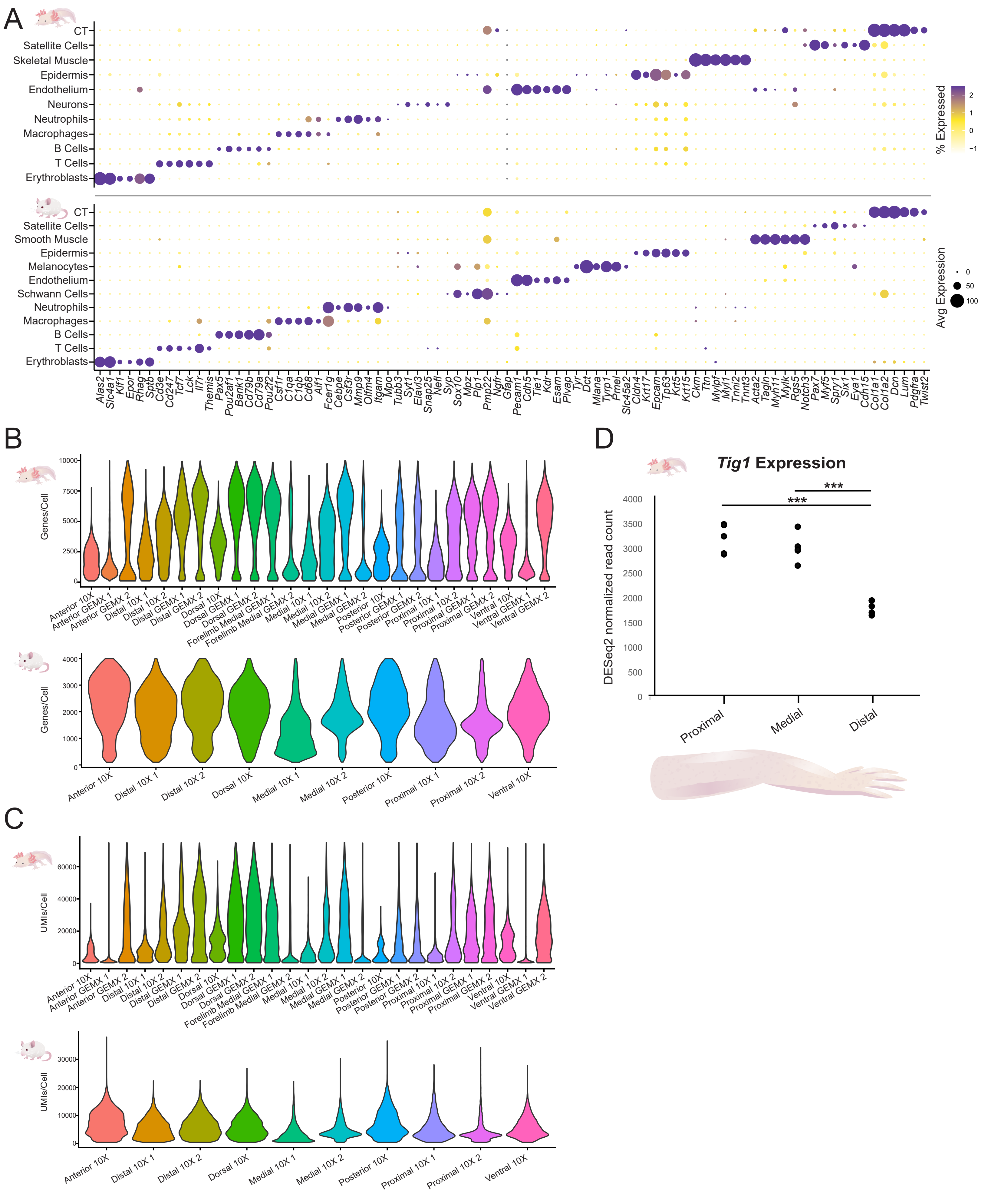

### Supplementary Fig. 2 (Fig. 2S1)

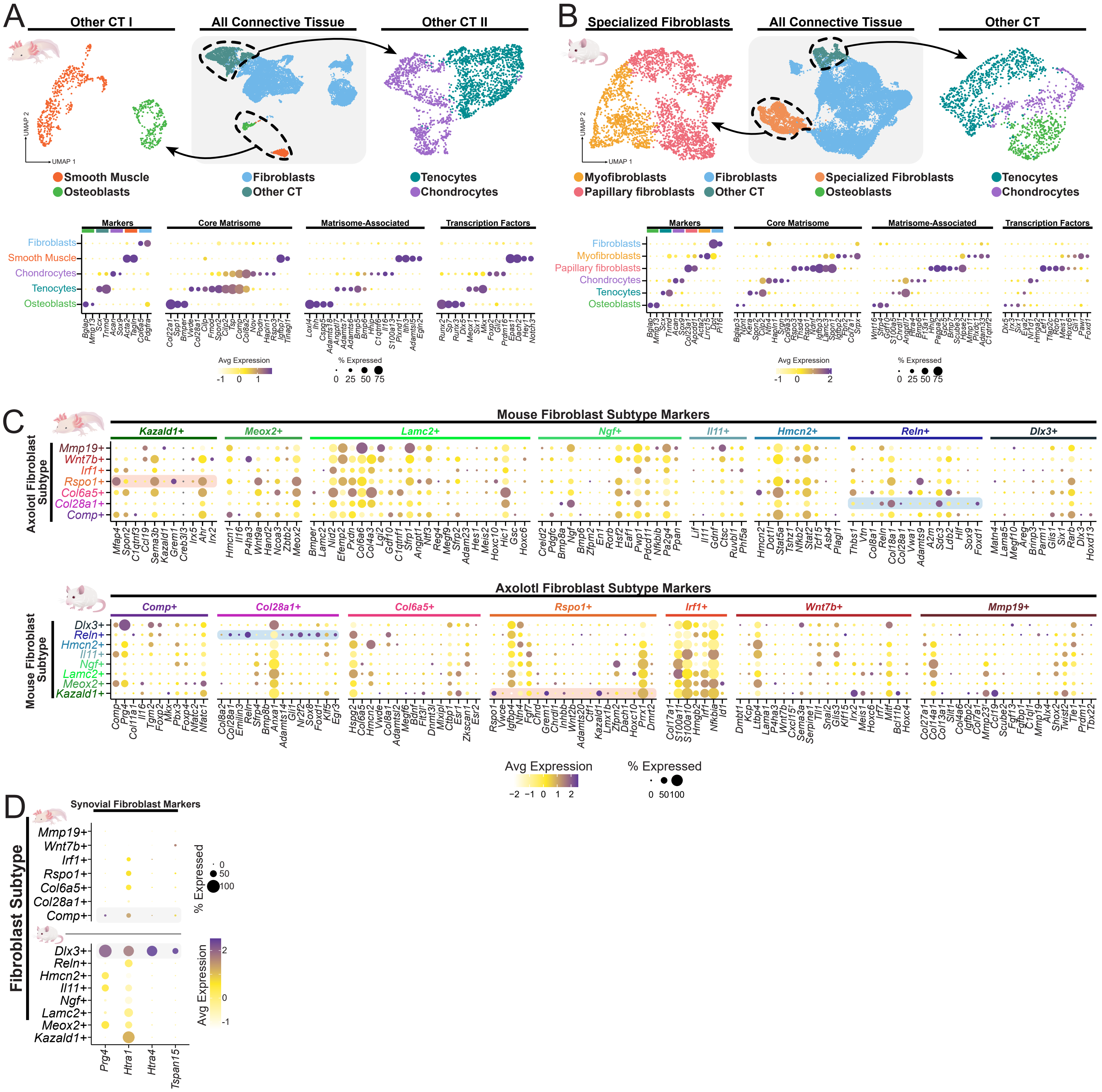

### Supplementary Fig. 3 (Fig. 3S1)

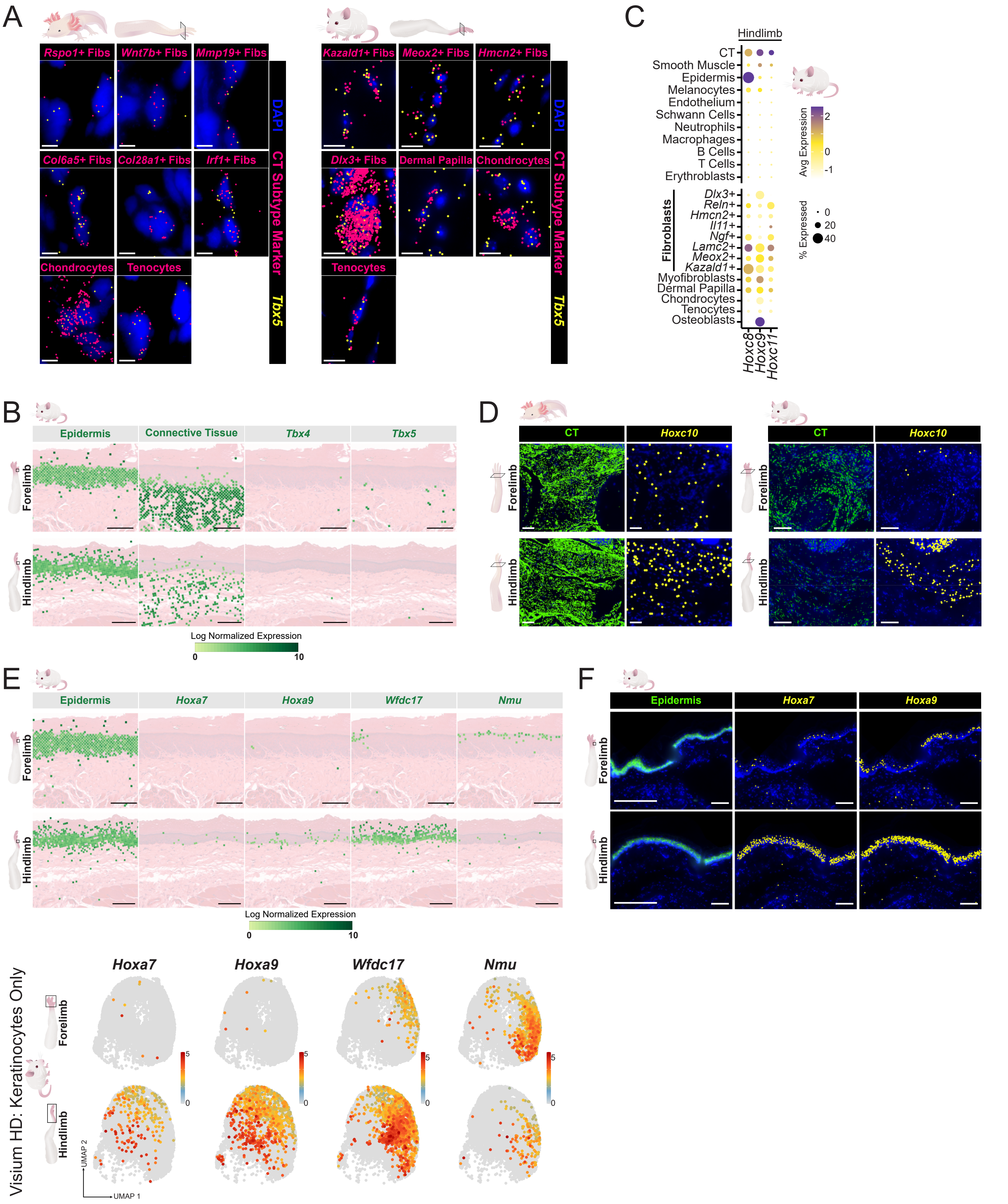

### Supplementary Fig. 4 (Fig. 3S2)

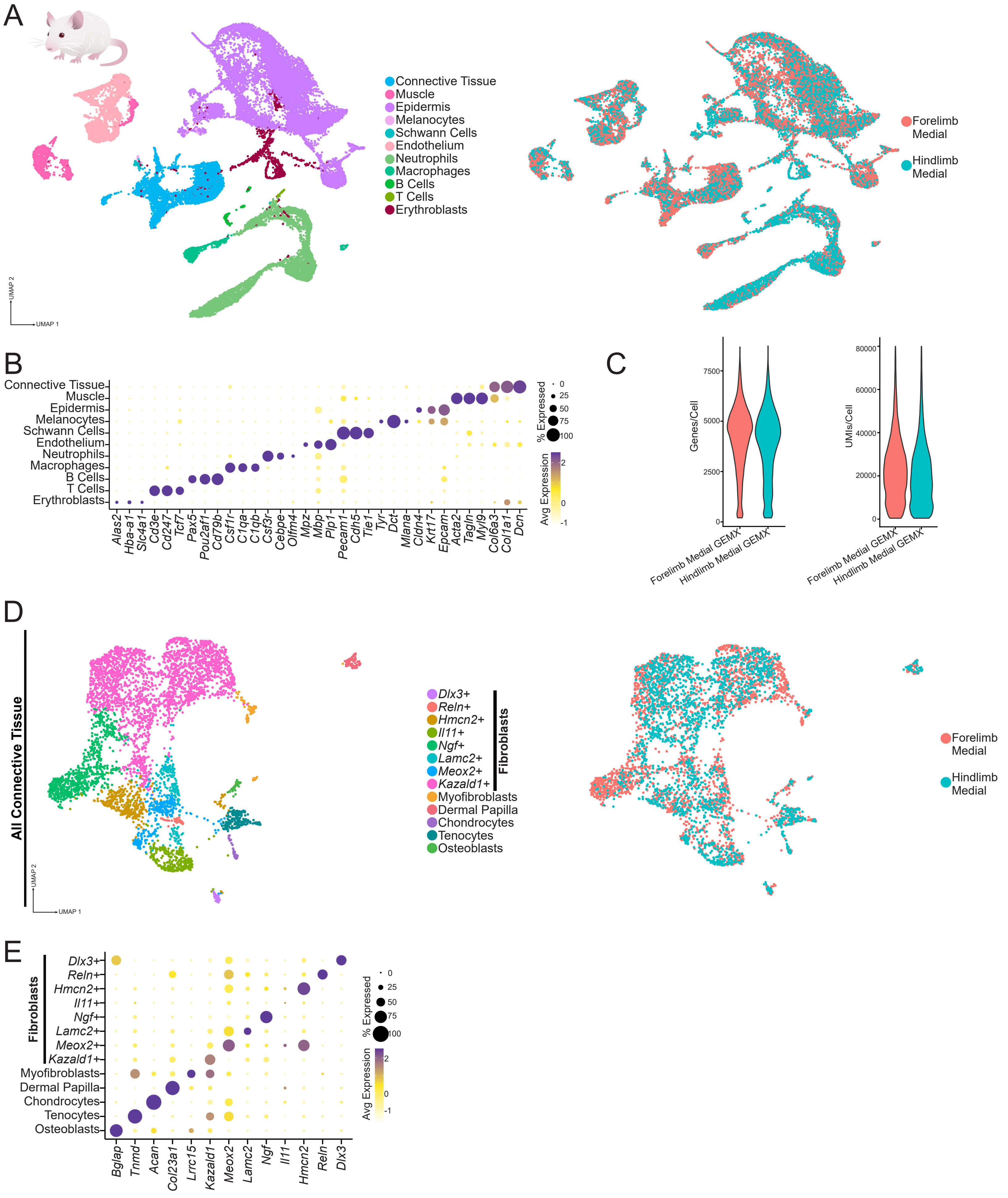

### Supplementary Fig. 5 (Fig. 4S1)

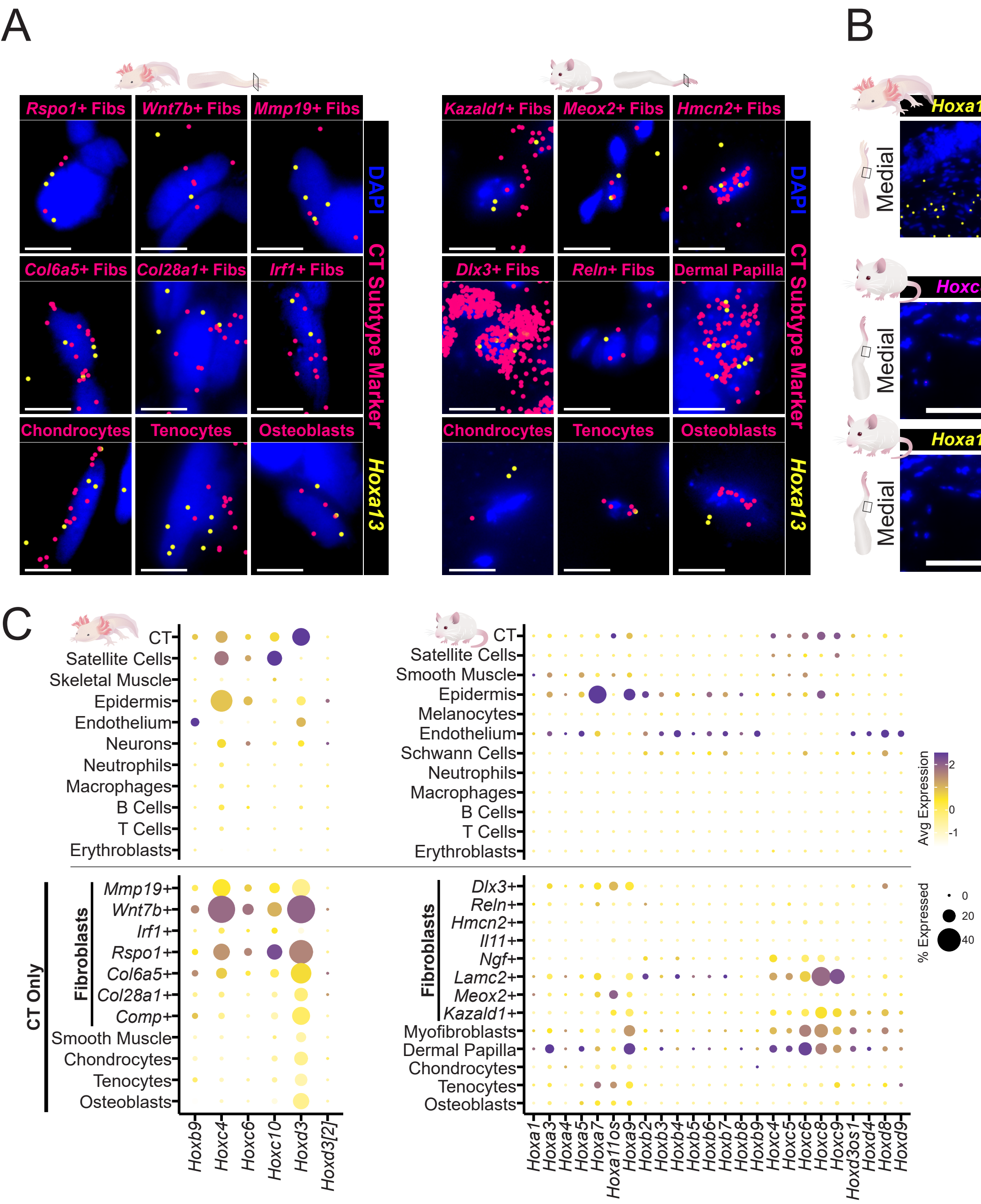

### Supplementary Fig. 6 (Fig. 4S2)

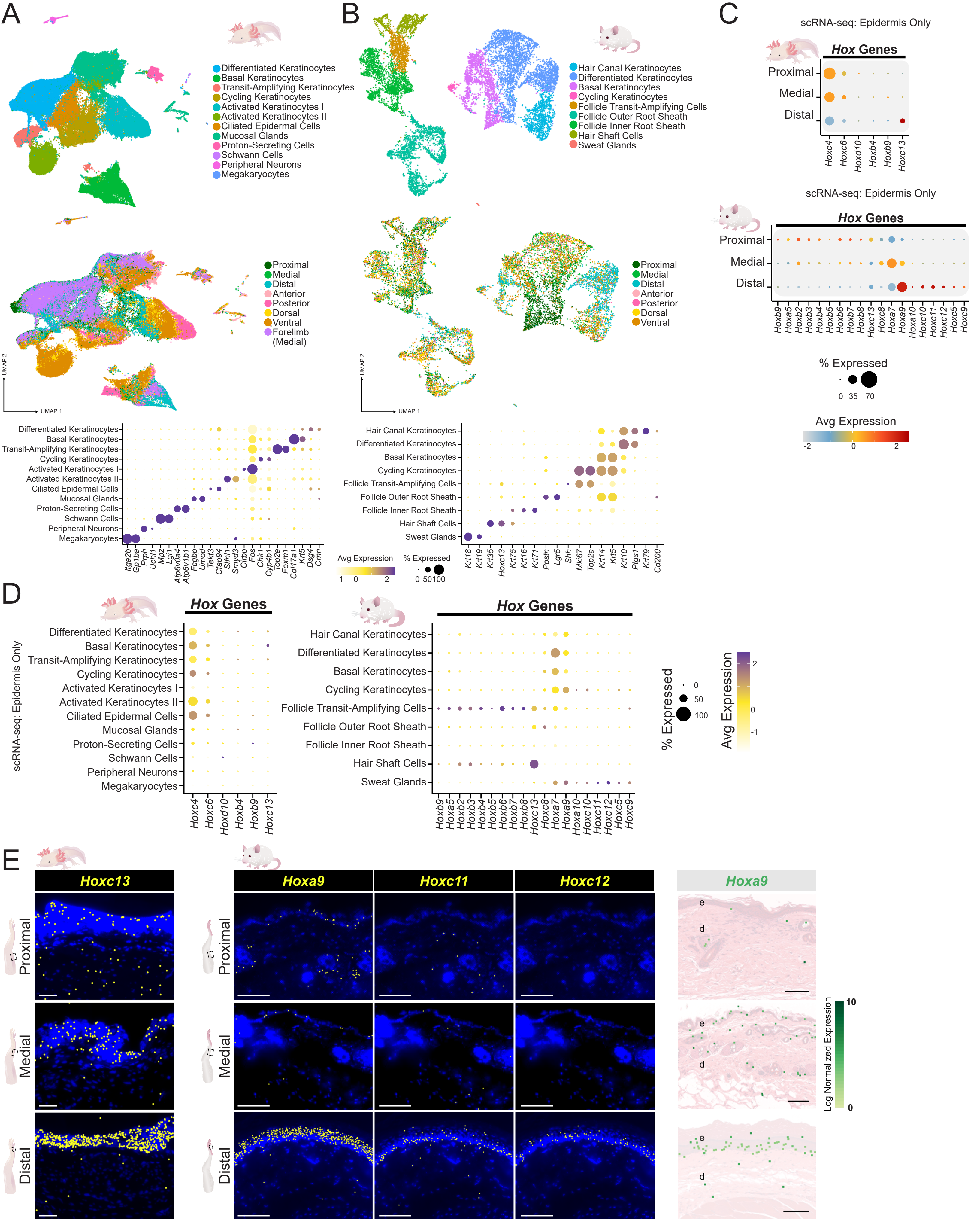

### Supplementary Fig. 7 (Fig. 5S1)

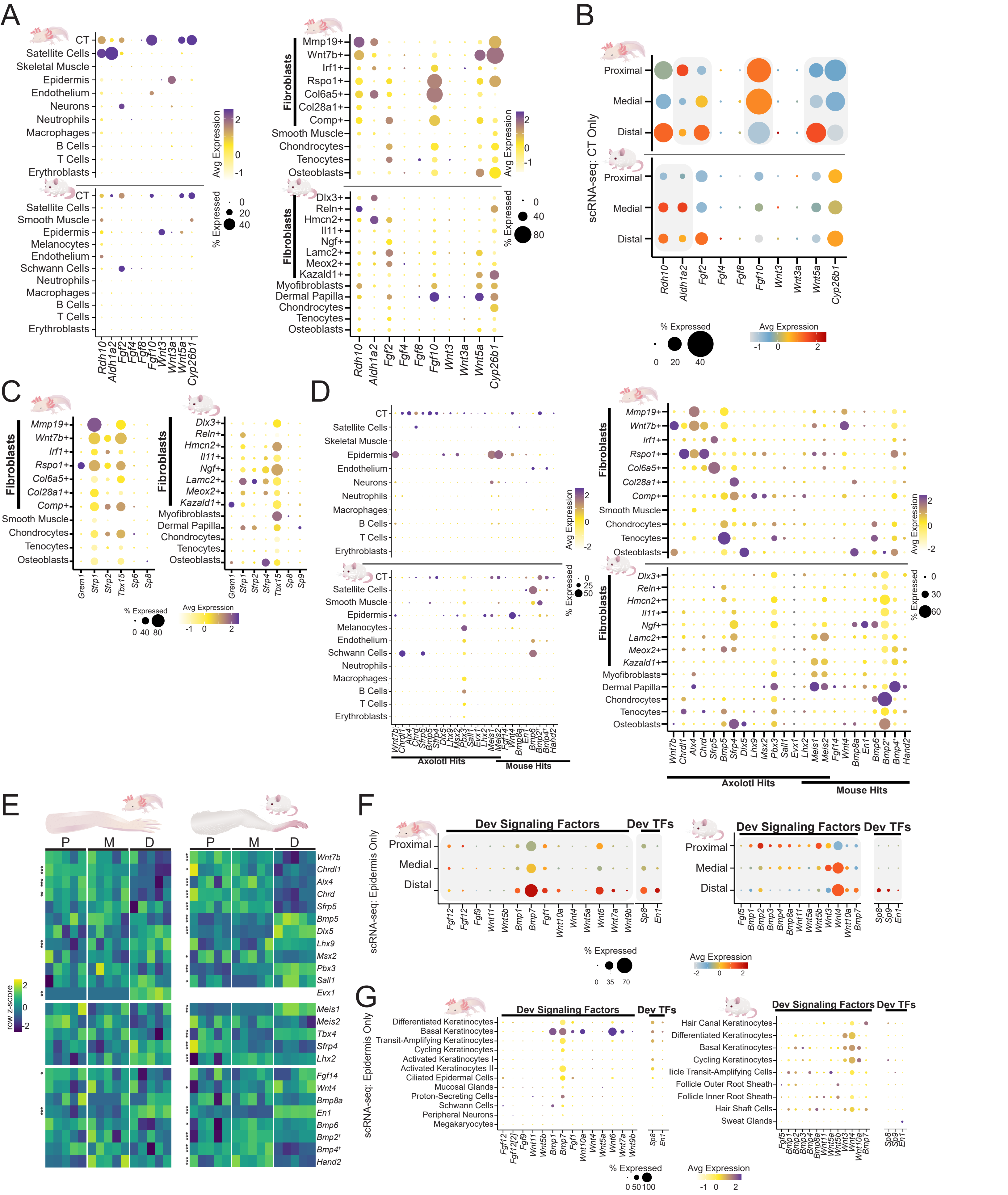

### Supplementary Fig. 8 (Fig. 6S1)

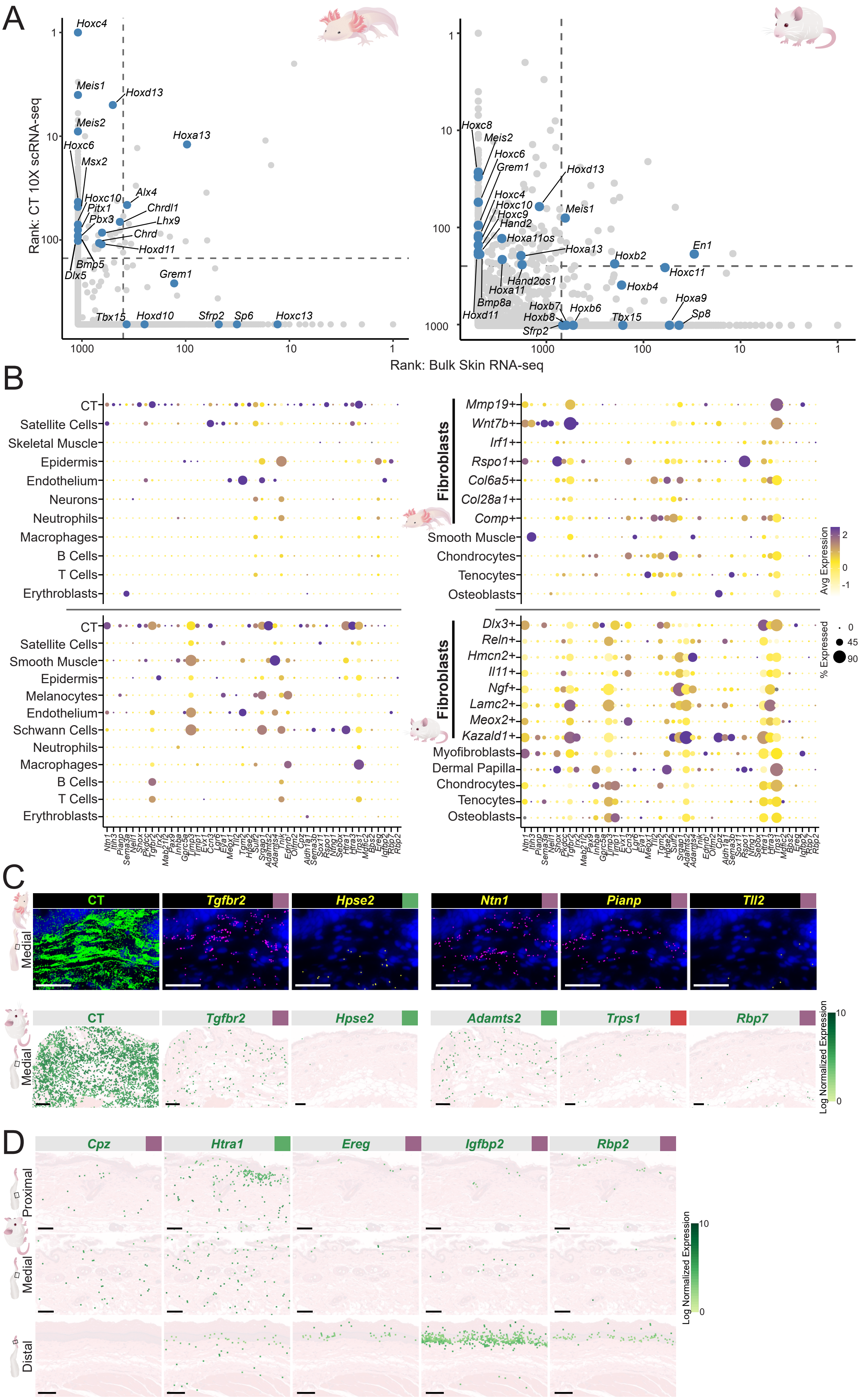

### Supplementary Fig. 9 (Fig. 6S2)

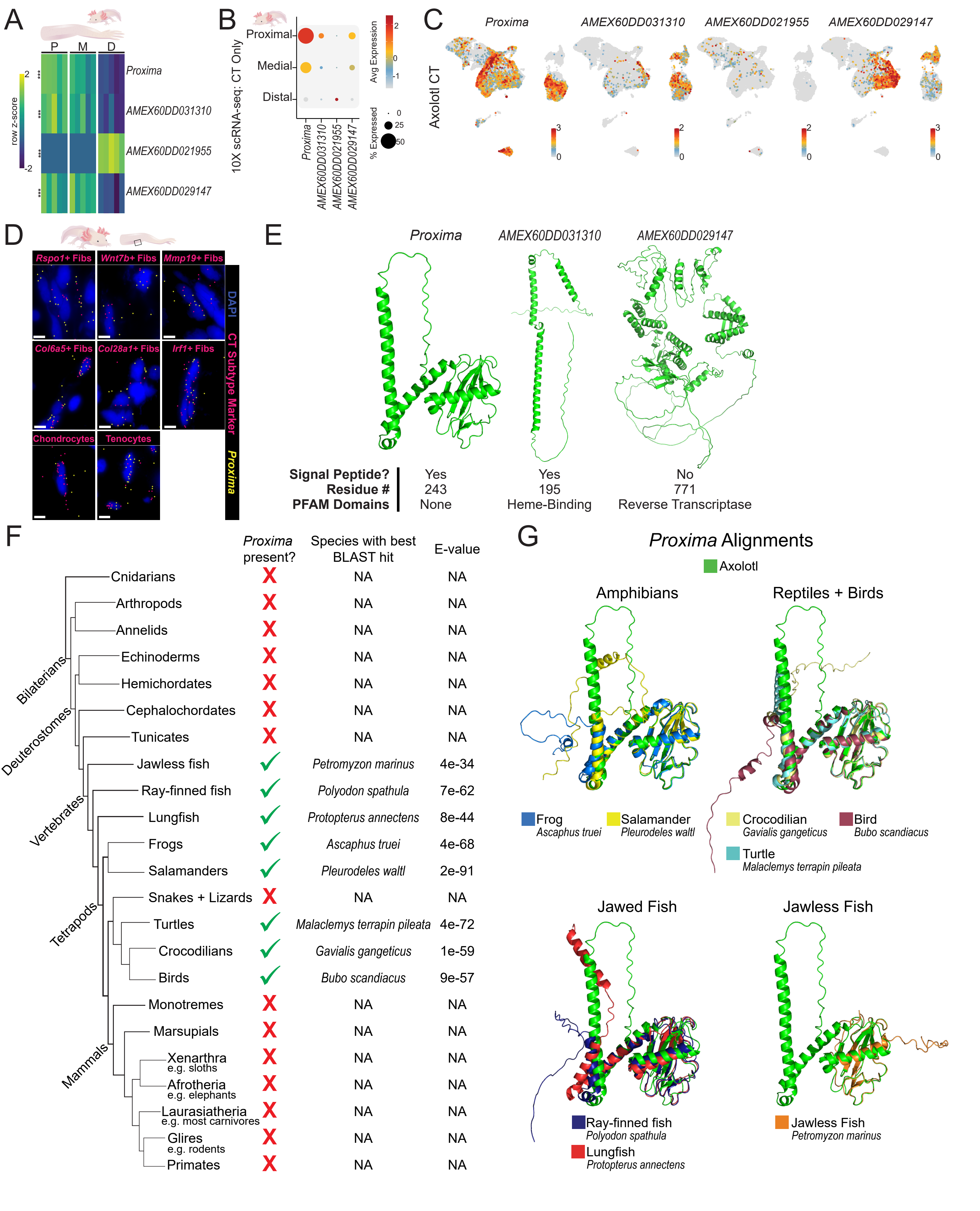

### Supplementary Fig. 10 (Fig. 6S3)

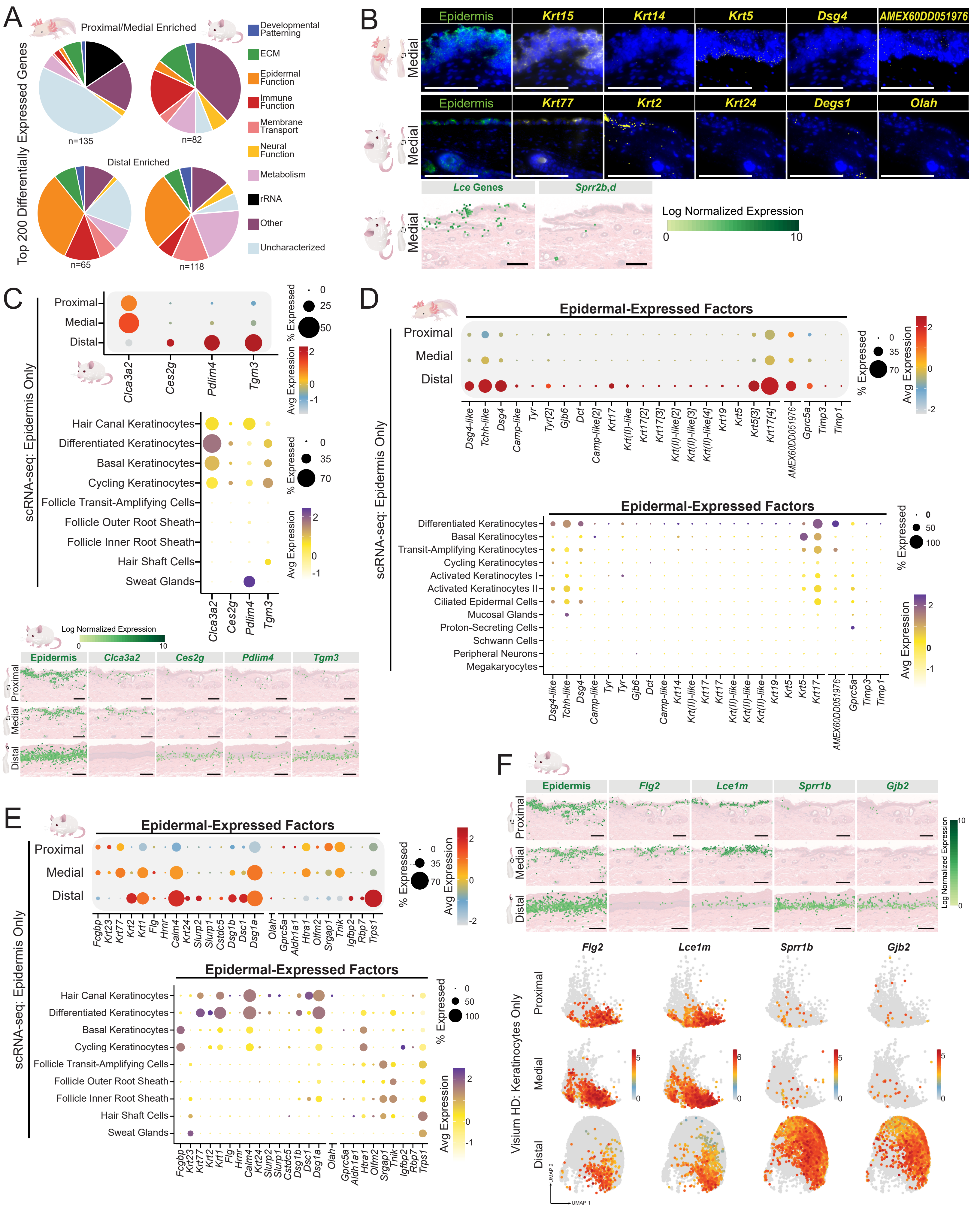

### Supplementary Fig. 11 (Fig. 6S4)

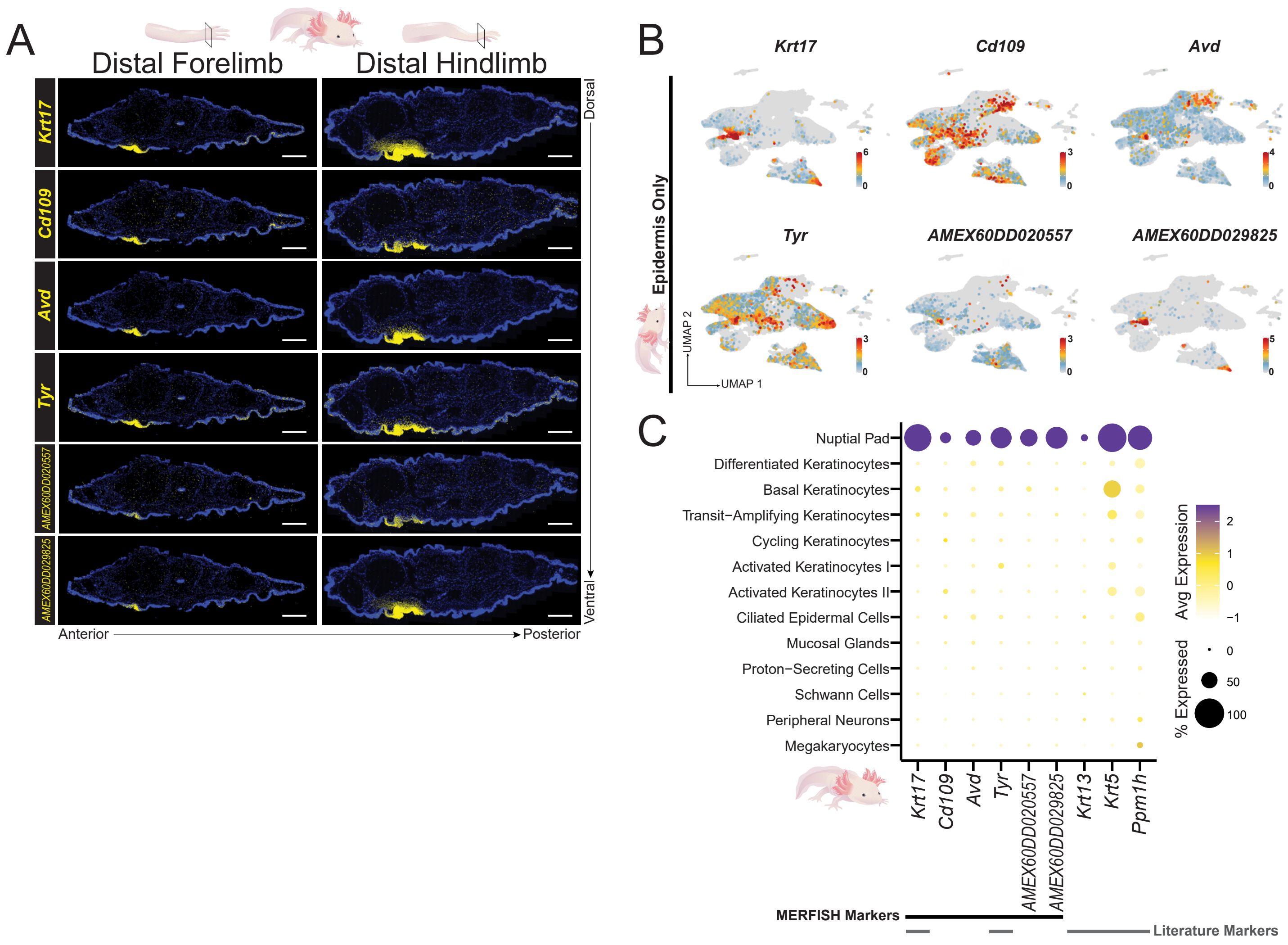

### Supplementary Fig. 12 (Fig. 7S1)

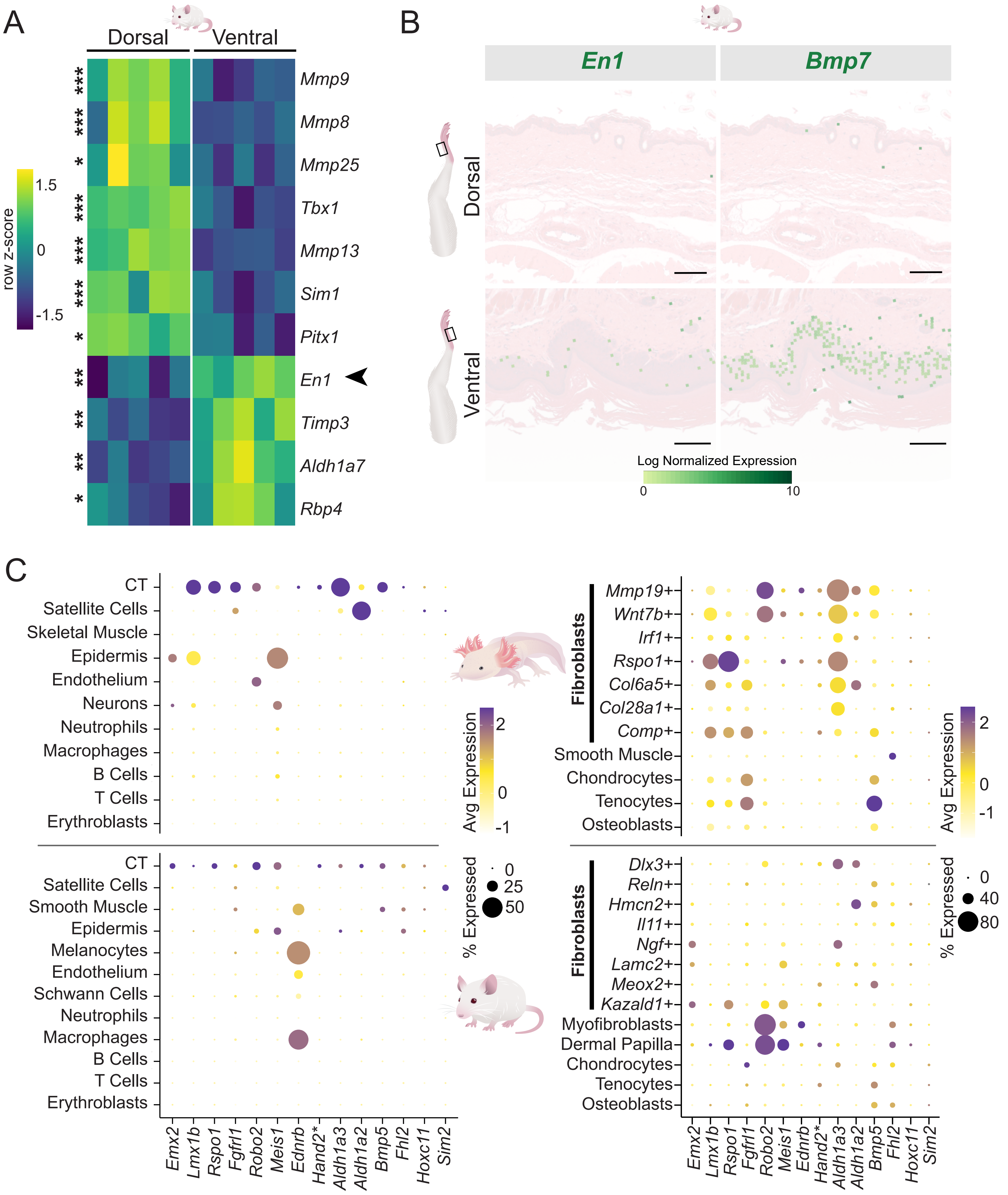

### Supplementary Fig. 13 (Fig. 8S1)

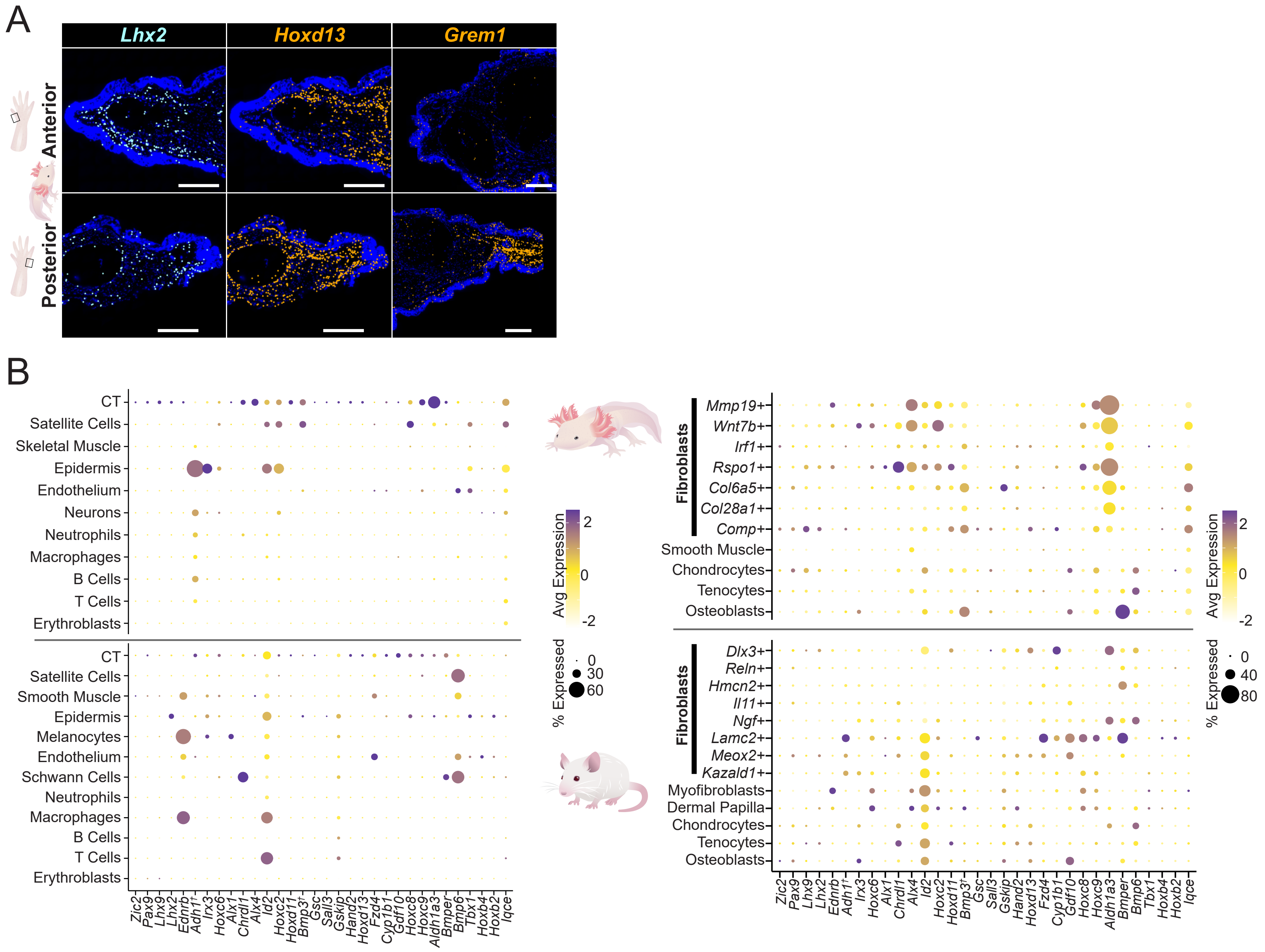

### Supplementary Fig. 14 (Fig. 9S1)

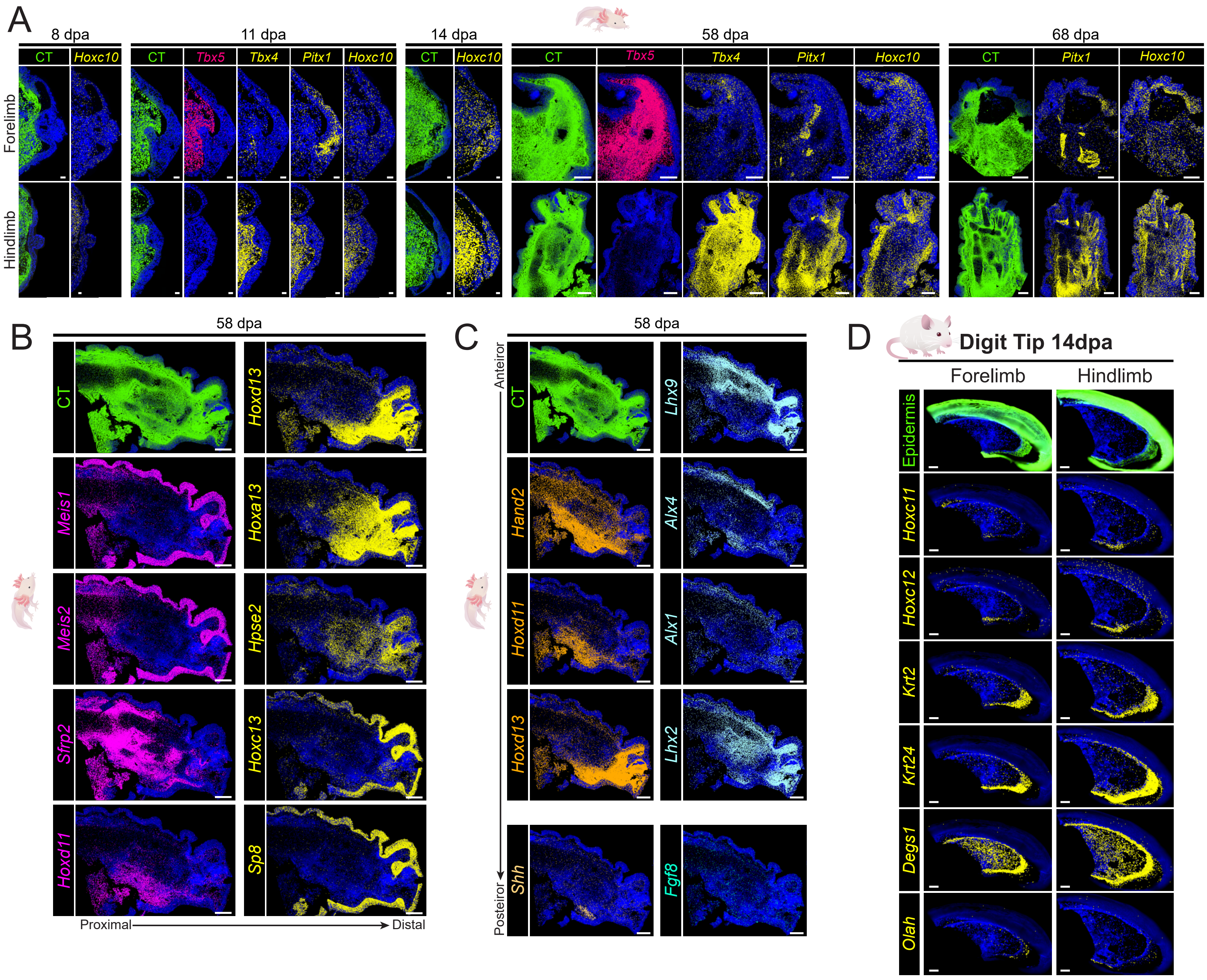

### Supplementary Fig. 15 (Fig. 9S2)

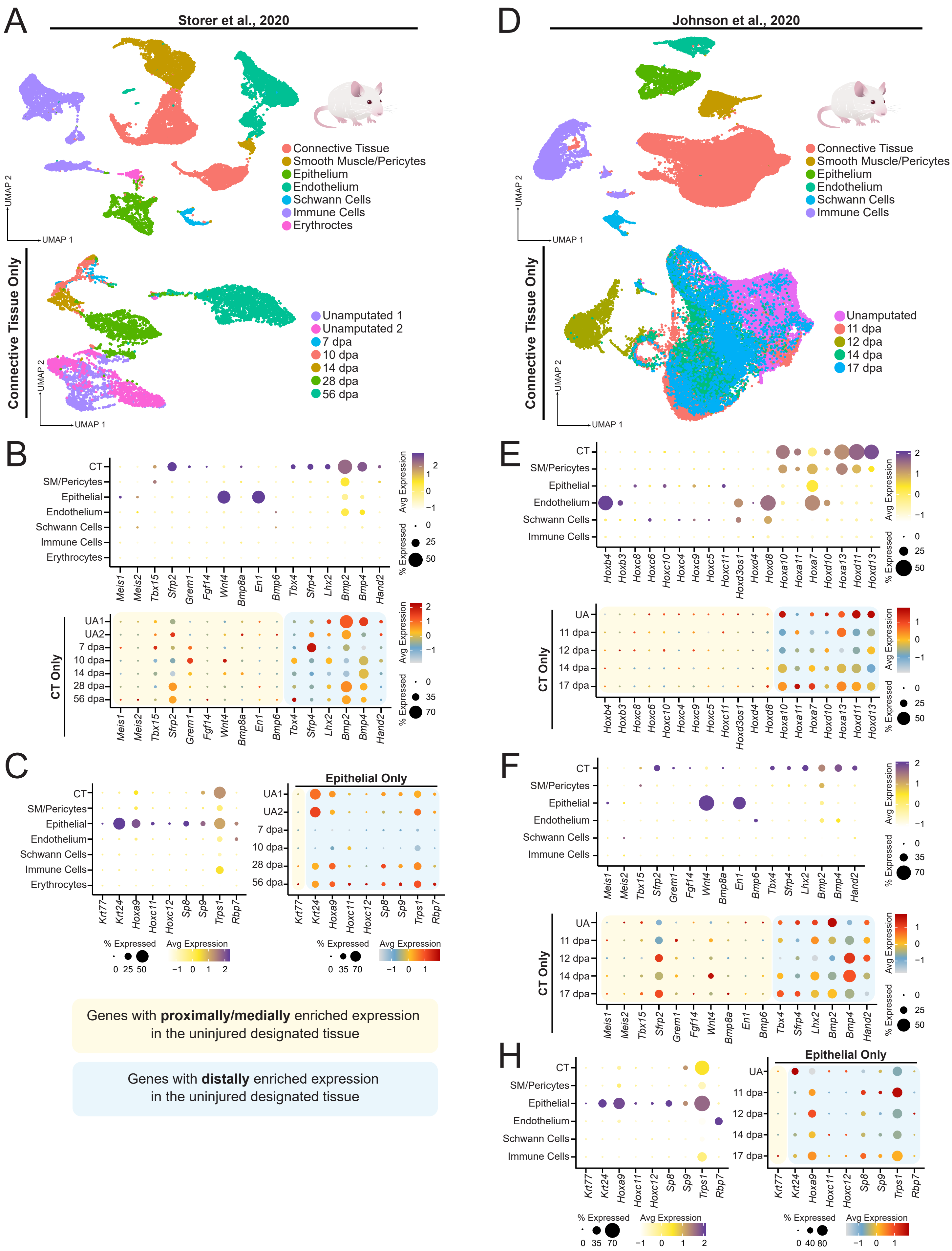
